# The Influence of Demographic History and Genetic Architecture on Complex Traits via Runs of Homozygosity

**DOI:** 10.64898/2025.12.17.694908

**Authors:** Mingzuyu Pan, Zachary A Szpiech

## Abstract

Runs of homozygosity (ROH) are contiguous genomic regions where all sites are homozygous, inherited from identical haplotypes due to shared ancestry. The number and length of ROH in individuals varies based on population history and sociocultural behaviors. Although often discussed in the context of inbreeding, ROH are ubiquitous in putatively outbred human populations, and their prevalence are associated with multiple complex traits, including height and measures of lung function. Importantly, long ROH have been shown to be enriched for deleterious alleles, suggesting a mechanism by which ROH prevalence can influence traits. Here we employ realistic forward-in-time population genetic simulations and a flexible quantitative model of a generic complex phenotype to explore how population history and genetic architecture influence ROH associations with a generic quantitative phenotype. We show that ROH are important for all simulated demographic histories and genetic architectures but especially when phenotypes have a recessive component. This is even more prominent when the rare-allele contribution to the phenotype is upweighted and in high-diversity populations (e.g. African). For a fully recessive phenotype, ROH can account for 25-55% of an individual’s total genotypic score, depending on demographic history and rare-allele weight. Our results emphasize the utility of ROH in helping to explain phenotype variation across different population histories and genetic architectures.

## Introduction

Runs of homozygosity (ROH) are contiguous genomic regions in an individual where all sites are homozygous, resulting from the inheritance of identical haplotypes from both parents due to common ancestry at some point in the past. In 1999, Broman and Weber^1^ first identified numerous long homozygous chromosomal segments using genomic data from the Centre d’Étude du Polymorphisme Humain (CEPH). At that time, researchers hypothesized that these long homozygous segments are prevalent even in outbred populations, potentially providing more insights into human health and broader population genetics than previously expected. Several years later, in 2006, Gibson^2^ analyzed the number, length and distribution of long homozygous chromosomal segments from outbred HapMap populations and identified three outlier individuals with extremely long and abundant homozygous segments. In the same year, Li et al^3^ reported similar observations in a Taiwanese population and suggested these long homozygous chromosomal segments likely represent autozygosity instead of the effects of gene deletion, recombination, or other genomic events. These findings prompted the formal definition of Runs of Homozygosity (ROH)^4^ and the associated metric F_ROH_,^5^ a novel estimator of the inbreeding coefficient.

Building upon this early work, researchers have accumulated increasingly large ROH datasets across diverse populations. In practice, ROH are not directly observed but are inferred from genotype array or sequence data using algorithmic approaches that apply thresholds for marker density, allowable missingness and heterozygosity, and minimum segment length. These studies have shown that ROH are ubiquitously distributed across putatively outbred global human populations.^6, 7^ For different populations, their unique demographic history and sociocultural preferences, for example, consanguineous marriage, can shape the length and number of ROH among the individuals of their contemporary populations.^8, 9^ Indeed, ROH prevalence has been used as a powerful tool in population genetics to provide insights about demographic events such as bottlenecks, founder events, and inbreeding in humans and other species.^10–15^

ROH analyses have deepened our understanding of the influence of inbreeding depression on complex traits in various species.^16–18^ For example, a study in wild Soay sheep^19^ found that long ROH reduce the likelihood of surviving the first winter more than short ROH. Across both wild and managed populations, ROH have also been used to study the impact of inbreeding and inbreeding depression on milk production traits^20–22^ in livestock. ROH also informs selection analyses in horse and dog breeds^23, 24^ and supports the design of management programs for endangered populations such as wild wolves,^25^ North American Thoroughbred horses,^26^ and Indian tigers.^27^

ROH are also informative in medical genomics studies to elucidate the genetic basis of human diseases.^28^ Previous studies have demonstrated the associations between ROH and increased risk of schizophrenia,^4, 29^ autism,^30^ human height,^31, 32^ cancer,^33, 34^ blood pressure,^35, 36^ depression,^37^ and cardiovascular diseases,^38, 39^ which highlights the value of ROH in assessing genetic health risks within populations and across populations.^40^ Total ROH levels have even been associated with direct fitness effects in humans.^41^ However, these associations have not been consistently replicated across populations, and the sources of this heterogeneity remain poorly understood.

Indeed, previous studies have suggested the mechanism by which ROH may be associated with traits. Szpiech et al.^42^ demonstrated that long runs of homozygosity are enriched for deleterious mutations worldwide. These results demonstrated the potential of long runs of homozygosity to harbor significant number of rare deleterious mutations, which are usually low frequency in the population but get paired into homozygotes within ROH. In a study of ROH in admixed populations, Szpiech et al.^43^ demonstrated that ROH overlapping African ancestry regions accumulated deleterious homozygotes at a higher rate compared to those overlapping European or Native American ancestry segments, demonstrating that demographic history influences the strength of enrichment. These results suggest that demographic history may play an important role in understanding the relationship between ROH and the genetic architecture of complex traits.

Unlike Mendelian traits driven primarily by variants at single loci, complex traits possess a polygenic architecture, arising from the cumulative effects of multiple genetic variants and their interplay with environmental factors. Genome-wide association studies (GWAS) and polygenic risk scores (PRS) have provided robust quantitative frameworks for studying complex traits, providing statistical tools to understand the genetic basis of complex traits. However, their effectiveness is still limited by their inability to detect very rare causal alleles^44^ and by population biases. Although many rare variant association tests (RVATs) have been developed,^45, 46^ these methods often fail to account for the evolutionary forces that help to shape genetic architecture. Existing studies have shown that when realistic evolutionary factors such as population growth and natural selection are included, the statistical performance of these methods diminishes.^44^ As a result, it remains difficult to detect associations between rare variants and phenotypes or to draw general conclusions. In contrast, ROH, by capturing homozygous regions enriched for rare, deleterious homozygotes^42^ and reflecting unique population demographic histories, offer a complementary perspective for studying genetic contributions to complex traits.

Previous studies on the relationship between ROH and complex traits have generally examined trait values directly, for example, by regressing phenotype values on total ROH burden, rather than decomposing phenotypic variance. We follow this approach and focus on decomposing individual genotypic scores rather than genetic variance. Importantly, the inconsistent associations observed across these empirical studies are precisely what our study aims to explain: by systematically varying genetic architecture and demographic history in simulation, we investigate the conditions under which ROH contribute to genotypic scores and the conditions under which they do not. Because ROH are reshuffled each generation, whether decomposing genetic variance into ROH and non-ROH components is meaningful remains an open question, though we still present variance decomposition as a supplementary analysis. To address these questions, we use forward-in-time simulations with SLiM combined with coalescent simulations with msprime. In empirical data, genetic architecture and demographic history are always confounded: it is not possible to determine whether differences in ROH-trait associations across populations are due to differences in dominance architecture, demographic history, or both. Simulation allows us to disentangle these factors by holding one constant while varying the other, providing complete ground truth including the location, effect size, and dominance coefficient of every causal variant. This theoretical framework provides a basis for interpreting the inconsistent empirical results observed across populations. Without it, these inconsistencies remain scattered observations that cannot inform each other.

Specifically, first, we employ forward-in-time simulations that incorporate realistic genome structure and an explicit out-of-Africa demographic model. Second, we couple natural selection to phenotypic effect sizes through a tunable parameter (τ) that controls the relative contribution of rare versus common variants. Third, we systematically examine four dominance regimes, including a mixed model informed by empirical inheritance-pattern annotations. The three single-dominance models (fully recessive, fully additive, and fully dominant) represent three qualitatively distinct regimes of how ROH interacts with genetic effects, and together they characterize the full parameter space: any intermediate dominance coefficient will produce behavior that falls between these three regimes. We believe this is a standard approach in theoretical population genetics, where boundary conditions are used to reveal the qualitative structure of a system. Simulating a continuous distribution of dominance coefficients would require specifying the shape of that distribution, which is itself empirically unknown, and would introduce additional unverifiable assumptions rather than increasing realism. Finally, we introduce ROH-class-specific metrics, including per-cM genotypic-score contributions and proportion-of-genotypic-score partitions, to quantify how different ROH classes contribute to complex traits. In addition, we conduct a comprehensive set of robustness analyses to evaluate the sensitivity of our conclusions to key methodological choices, including sparse versus non-sparse genetic architectures, conditional versus unconditional per-unit-length analyses, diagnostics of causal variant enrichment within ROH at both the allele and homozygote levels, alternative frequency-independent ROH-calling methods, and cM-based versus Mb-based ROH classification thresholds, among others.

Instead of considering a specific phenotype, we simulated the genetic component under an omnigenic model. The omnigenic model proposes that complex traits are determined by a vast number of genes distributed across gene regulatory networks and the whole genome, rather than by a small number of specific genes.^47^ The existence of phenotype correlations with total ROH levels is consistent with a broadly distributed recessive load across the genome, motivating the use of an omnigenic framework for modeling complex-trait architecture.^48^ Thus, by simulating these genetic effects across the entire genome, our work establishes a generic quantitative framework for assessing the relationship between ROH and complex phenotypes. Additionally, we investigate the influence of dominance coefficients and assess the significance of rare allele contributions to genotypic scores (the total genetic contribution to the phenotype per individual), offering potential guidance for future methodological developments in population and medical genetics.

## Results

Figures in the main text are numbered in pairs: each analysis based on deleterious mutations is paired with a neutral counterpart (Figures 1 & 3, 2 & 4, 5 & 6, 7 & 8). We present all deleterious results first, followed by the neutral results, so figures are not cited in strict numerical order.

**Figure 1.**
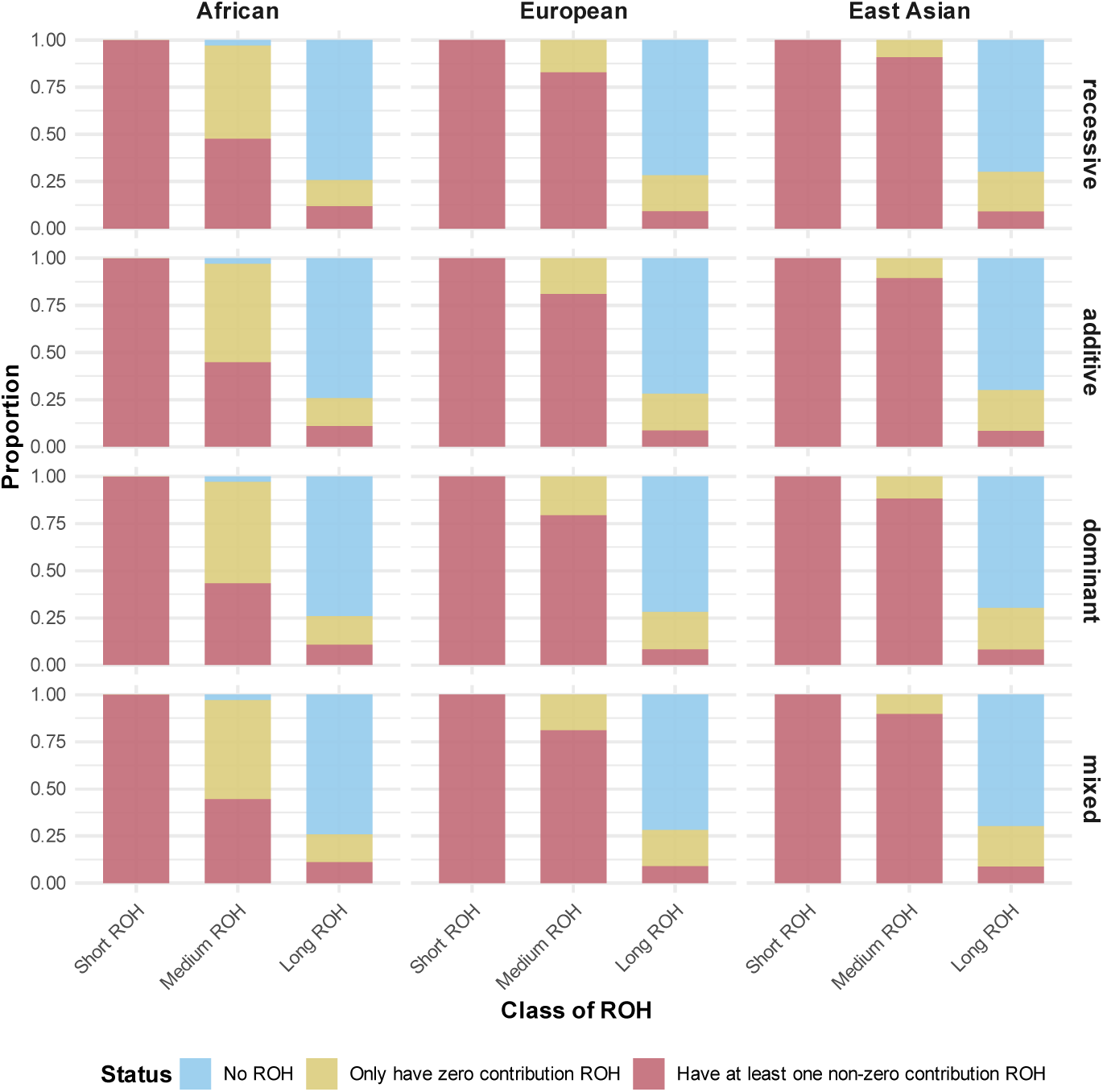
Proportion of individuals with ROH contributing to genotypic score across various population histories and genetic architectures. Genotypic scores are only contributed by deleterious mutations. Blue bars represent the individuals lacking ROH (thus no genotypic score contribution); red bars represent the individuals having at least one ROH with a genotypic score contribution; yellow bars represent the individuals having ROH with no genotypic score contribution. Region definitions: Short ROH (< 0.25cM); Medium ROH (0.25-1cM); Long ROH (> 1cM).

### Proportion of individuals with ROH contributing to the genotypic score

Runs of Homozygosity were called in each individual and categorized into three size classes, Short ROH (<0.25 cM), Medium ROH (0.25-1.0 cM), and Long ROH (>1.0 cM). Figure 1 shows the mean proportion of individuals across simulation replicates for each ROH size category, where blue represents the proportion of individuals that have no ROH of a particular size, red represents the proportion of individuals with at least 1 ROH of that size that also contributes to the genotypic score, and yellow represents the proportion of individuals that have ROH of that size but none contribute to the genotypic score.

We find that phenotype architecture doesn’t have strong influence on these proportions, but demographic history does influence the proportion of individuals with ROH > 0.25 cM that contribute to the genotypic score. In the African population, about 50% of individuals have at least one medium ROH with non-zero genotypic score contribution. The proportion of individuals in the European population having medium ROH with non-zero genotypic score contribution is higher, reaching 75%, and the proportion in the East Asian population is higher still, reaching 87.5%.

Interestingly, for long ROH, the differences between populations are small than those of medium ROH. As these populations are simulated with no explicit inbreeding, the proportion of individuals with long ROH are low, and approximately 10% of individuals from each of these three populations have long ROH with non-zero genotypic score contribution. Interestingly, the proportion of individuals in the African population with at least one long ROH contributing to the genotypic score is slightly larger than that in the European and East Asian populations (approximately 12% vs. 10% and 10%, respectively), even though the total proportion of individuals with long ROH is lower (approximately 25% vs. 27% and 28%, respectively).

### Genotypic score contribution per centimorgan

Next, we consider the relative contributions of ROH to the genotypic score across a range of architectures. To fairly compare ROH of different sizes, we normalize by ROH length, analyzing the genotypic-score contribution per centimorgan. This per-unit calculation is performed at the individual level: each individual’s genotypic score attributable to a given ROH class is divided by that individual’s total length of ROH in that class, rather than computed per segment. We refer to the analysis that considers only individuals with at least one ROH carrying a non-zero genotypic-score contribution as the “conditional” analysis, and the analysis that considers all individuals as the “unconditional” analysis. These two scenarios are labeled accordingly in all figures and captions. The main-text figures present conditional results; unconditional methods, results, and figures are presented in the robustness analysis section (Figures S14–S15).

#### Additive Model

Under the additive model the functional allele has a phenotype effect regardless of zygosity, with heterozygotes having half the effect of homozygotes. Figures 2 and S1 shows that short ROH contribute significantly more per cM to genotypic scores than non-ROH, regardless of demographic history and the value of tau. Interestingly, short ROH also contribute significantly more per cM to genotypic score than long ROH across these parameters, and long ROH contribute more than medium ROH, except for larger tau in the European and East Asian populations. In contrast, long ROH and medium ROH were more sensitive to the value of tau and demographic history. When emphasis is shifted to common alleles (small tau), long and medium ROH contributed significantly more to genotypic score than non-ROH, however, when emphasis is shifted to rare alleles (large tau), this pattern flips.

**Figure 2.**
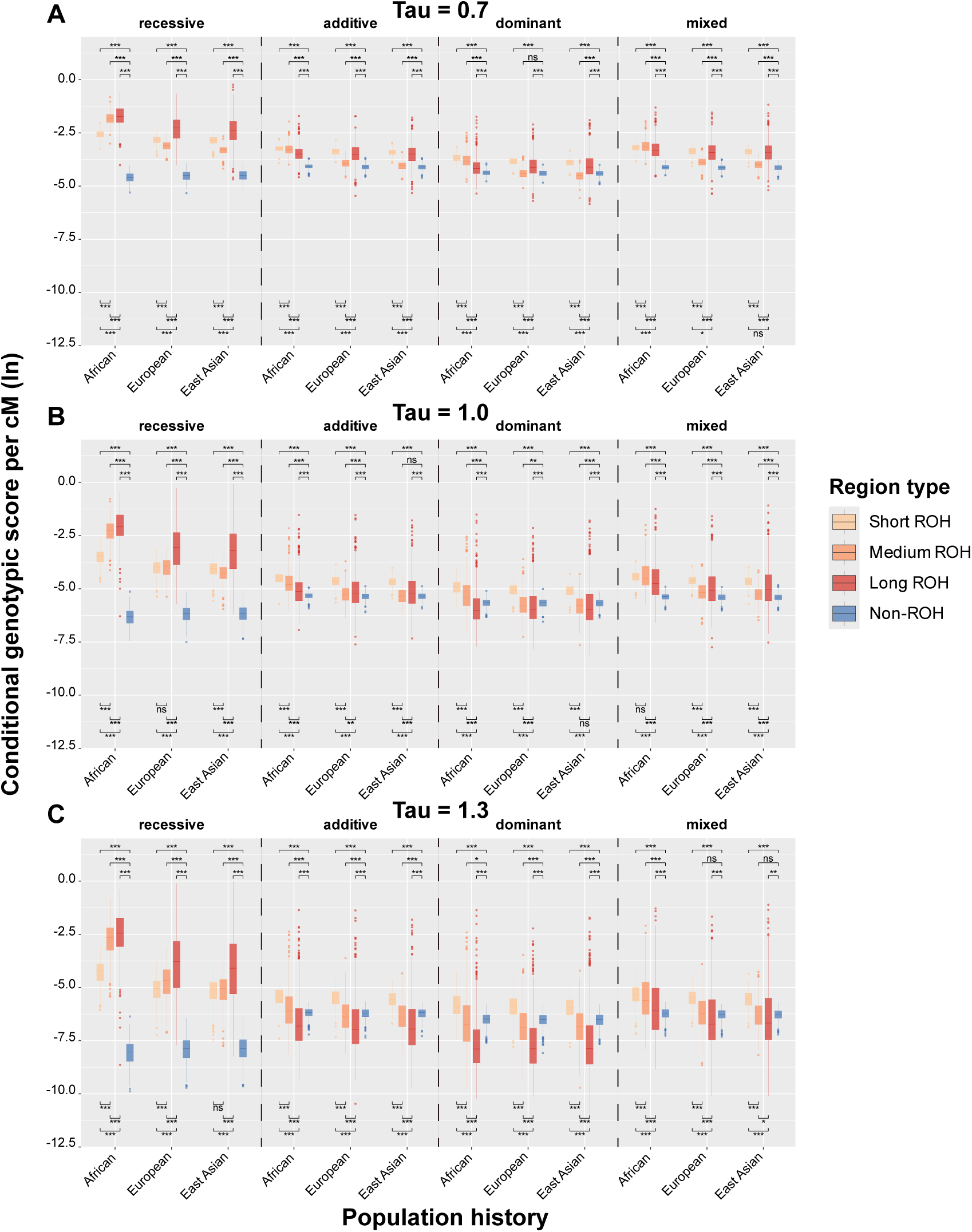
Conditional per-cM genotypic score contribution (log-transformed; deleterious mutations) for various regions, population histories, and genetic architectures. Regions are: Short ROH (< 0.25cM); Medium ROH (0.25-1cM); Long ROH (> 1cM); Non-ROH. genotypic scores are generated from deleterious mutations. The parameter tau is varied for different weighting of rare alleles in genotypic score calculation. (A) *τ* = 0.7, (B) *τ* = 1.0, (C) *τ* = 1.3. Significance levels for adjusted *P*-values: *** *P_ad_*_j_ < 0.001, ** *P_ad_*_j_ < 0.01, * *P_ad_*_j_ < 0.05, ns *P_ad_*_j_ ≥ 0.05.

#### Dominant Model

Under the dominant model the functional allele has a phenotype effect regardless of zygosity, however in this case both heterozygotes and homozygotes have full effect. Figures 2 and S1 shows that ROH contributions per cM to genotypic score follow similar patterns as in the additive model. Short ROH contribute significantly more per cM to genotypic scores than non-ROH and long ROH. For medium ROH, the influence of demographic history is clear. For the African population, medium ROH have a significantly higher contribution to genotypic score per cM compared to non-ROH except for the largest tau, however for European and East Asian populations, medium ROH generally contribute lower compared to non-ROH. Long ROH have significantly more contribution per cM compared to non-ROH only when emphasis is shifted to common alleles (small tau), but this pattern flips when emphasis is put on rare alleles (large tau).

#### Recessive Model

Under the recessive model, the functional allele has a phenotype effect only in homozygous form. Figures 2 and S1 shows that under the recessive model, all three ROH categories across the three populations exhibited significantly higher contributions to polygenic genotypic scores than non-ROH regions, with these differences being consistently statistically significant. Among the three ROH categories, long ROH regions exhibited an outstanding signal, having significantly higher genotypic score per cM than short, medium, or non ROH regions. This pattern was robust to changes in emphasis on common vs rare alleles (changing tau).

#### Mixed Dominance Model

Under the mixed dominance model, the relationship between functional allele phenotype effect and zygosity is randomly determined, with approximately 88.4% additive, 7.1% recessive, and 4.5% dominant (see Methods). Figures 2 and S1 shows trends that generally reflect the patterns of the single inheritance models mixed at the above proportions. Interestingly, as the genotypic score per cM for recessive inheritance models dramatically favors ROH over non-ROH regions, even a relatively small recessive component drives the importance of ROH regions across several parameters. For models with an emphasis on common alleles (small tau), short, medium, and large ROH show significantly higher genotypic score per cM compared to non-ROH. As emphasis shifts to rarer alleles (large tau), the influence of demographic history begins to differentiate patterns somewhat. Under an African demographic history, all ROH regions show significantly higher genotypic score per cM compared to non-ROH, but this pattern changes for European and East Asian demographic histories. For European histories only short and medium ROH have a significantly higher genotypic score per cM, and for East Asian histories only short ROH have a significantly higher genotypic score per cM.

#### Changing emphasis from common to rare alleles

Equation 1 transforms a mutation’s selection coefficient into a phenotype effect, and the parameter tau indirectly modulates whether a genotypic score is relatively more influenced by common variation (small tau) versus rare variation (large tau). This occurs since large tau accentuates large absolute selection coefficients more than small ones, and large absolute selection coefficients (i.e., highly deleterious alleles) are more likely to be rare. However, by changing the exponent tau, both the mean and variance of the phenotype effect distribution is changed, which may obscure important comparative patterns when looking across tau values. To address the changing variances, the total genotypic scores per individual are normalized by the standard deviation across all populations, however mean centering would disallow a log transform. To explore how changing tau influences the relative per-cM genotypic score distribution among ROH and non-ROH regions, figure 5 plots the difference between ROH and non-ROH log mean per-cM genotypic score contributions as a function of tau. In this figure, positive values indicate that the mean per-cM genotypic score contribution of the ROH is higher than the non-ROH, and negative values indicate that non-ROH per-cM genotypic score contribution is higher.

We find that genetic dominance has the largest effect on these patterns, with the largest positive values occurring with the recessive phenotype model across all ROH sizes. Furthermore, as tau increases weight on rare alleles, we find that ROH regions contain substantially more per-cM genotypic score contribution compared to non-ROH regions, with long ROH showing the highest differences and biggest changes. Demographic history also plays a role, with African demographies having the highest difference in ROH vs non-ROH per-cM genotypic score contribution.

In contrast to the recessive phenotype model, the additive and dominant phenotype models show more modest differences between ROH and non-ROH per-cM genotypic scores, with the dominant phenotype model generally showing the smallest positive differences. In some cases, especially for large tau among medium and long ROH, we find that non-ROH regions have an increased and higher per-cM genotypic score compared to ROH. Interestingly, changing tau has minimal effect on the difference between short ROH and non-ROH per-cM genotypic score contributions for these phenotype models, and, for non-African demographies under the dominant model, there is even a small increase in the per-cM genotypic score contribution difference for larger tau.

### The proportion of genotypic score explained by ROH

Next, we examine how much of the total genotypic score is explained by ROH versus non-ROH regions. Different from the per-centimorgan genotypic score calculation, we considered all the individuals here, with no regard to ROH content of the genomes.

Figure 7 illustrates these patterns divided by short, medium, long, and non-ROH regions. Generally, for dominant and additive phenotypes, non-ROH regions explain the vast majority of an individual’s total genotypic score. This is particularly notable for dominant phenotypes under the African demographic history, where ∼90% of the genotypic score is explained by mutations in non-ROH regions. European and East Asian demographic histories, which have experienced a historical bottleneck, see somewhat more total genotypic score explained by ROH for dominant and additive phenotypes. Notably, under a purely recessive phenotype model, total ROH explains a substantially larger proportion of the genotypic score, ranging from approximately 25% to 55%, depending on the demographic history and emphasis on common versus rare alleles (low versus high tau).

Indeed, while changing the emphasis of common versus rare variation doesn’t have an appreciable effect on the proportion of genotypic score explained for other inheritance regimes, for recessive phenotypes there is a clear trend. When the phenotype effect is shifted to more heavily favor rare alleles (large tau), ROH regions increasingly explain more of the genotypic score. In particular, the contribution proportion of short ROH regions to the genotypic scores ranged from 22 to 45%, medium ROH ranged from 4 to 10%, and Long ROH ranged from 0.5 to 2%.

### Phenotypes from neutral alleles

The previous results presented here considered a phenotype that is determined purely from deleterious alleles and whose total score is a function of the underlying selection coefficients of those alleles. We also considered the case where a phenotype was determined purely from neutral alleles (of similar number) simulated in the presence of deleterious alleles (i.e., background selection). In this case the genotypic score was determined by sampling from the same marginal distribution of selection coefficients and computing the phenotype effect using the same functional transformation, but with the actual selective effect of the alleles remaining neutral.

Similar to Figure 1, Figure 3 shows the mean proportion of individuals across simulation replicates for each ROH size category, where blue represents the proportion of individuals that have no ROH of a particular size, red represents the proportion of individuals with at least 1 ROH of that size that also contributes to the genotypic score, and yellow represents the proportion of individuals that have ROH of that size but none contribute to the genotypic score. As before, the differences within the results are primarily observed across the ROH size classes and population histories, while there are minimal differences across genetic architectures.

**Figure 3.**
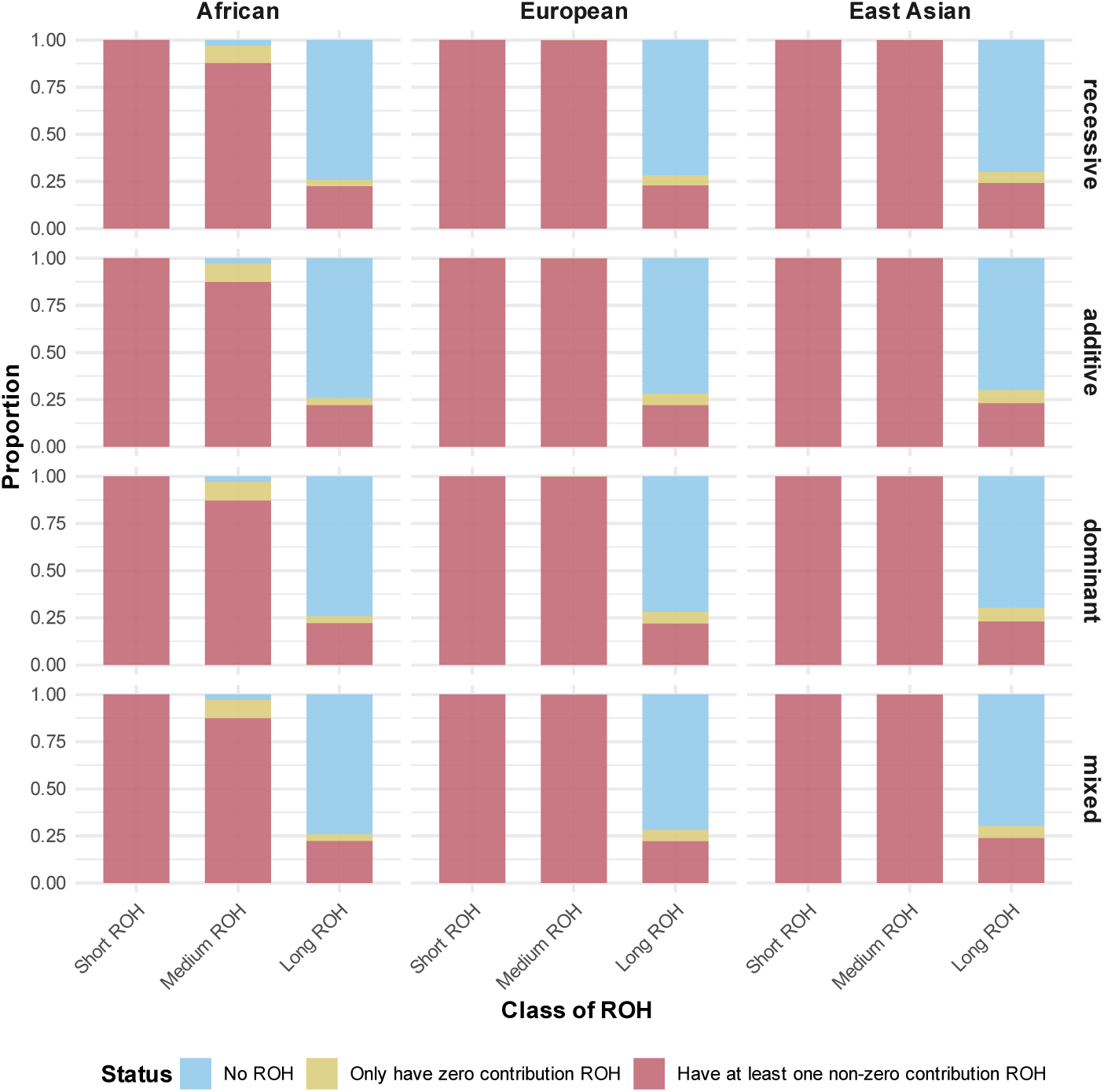
Proportion of individuals with ROH contributing to genotypic score across various population histories and genetic architectures. Genotypic scores are only contributed by neutral mutations. Blue bars represent the individuals lacking ROH (thus no genotypic score contribution); red bars represent the individuals having at least one ROH with a genotypic score contribution; yellow bars represent the individuals having ROH with no genotypic score contribution. Region definitions: Short ROH (< 0.25cM); Medium ROH (0.25-1cM); Long ROH (> 1cM).

Notably, although the proportion of individuals with ROH are approximately the same, the proportion of individuals with at least 1 ROH contributing to the genotypic score is higher for medium and long ROH when phenotypes are determined from neutral alleles. In the African population, 87.5% of individuals have medium ROH with non-zero genotypic score contribution. About 10% of individuals have medium ROH with zero genotypic score contribution, while a very small proportion do not have medium ROH. All individuals in the European and East Asian populations have medium ROH with non-zero genotypic score contribution. Slightly below 25% of individuals in each of these three populations have long ROH with non-zero genotypic score contribution. In all these populations, a small number of individuals having long ROH without genotypic score contribution. In particular, the proportion in the East Asian population is slightly higher than that in the European population, while the proportion in the European population is also slightly higher than that in the African population. Still, regardless of genetic architecture, all the individuals from these three populations have short ROH with non-zero genotypic score contribution.

Figures 4 and S2 show the genotypic score per cM for short, medium, long, and non ROH under this model for varying genetic architectures, demographic histories, and inheritance regimes. Generally, none of these variables have as conspicuous an effect compared to the case where phenotypes are determined by deleterious alleles.

**Figure 4.**
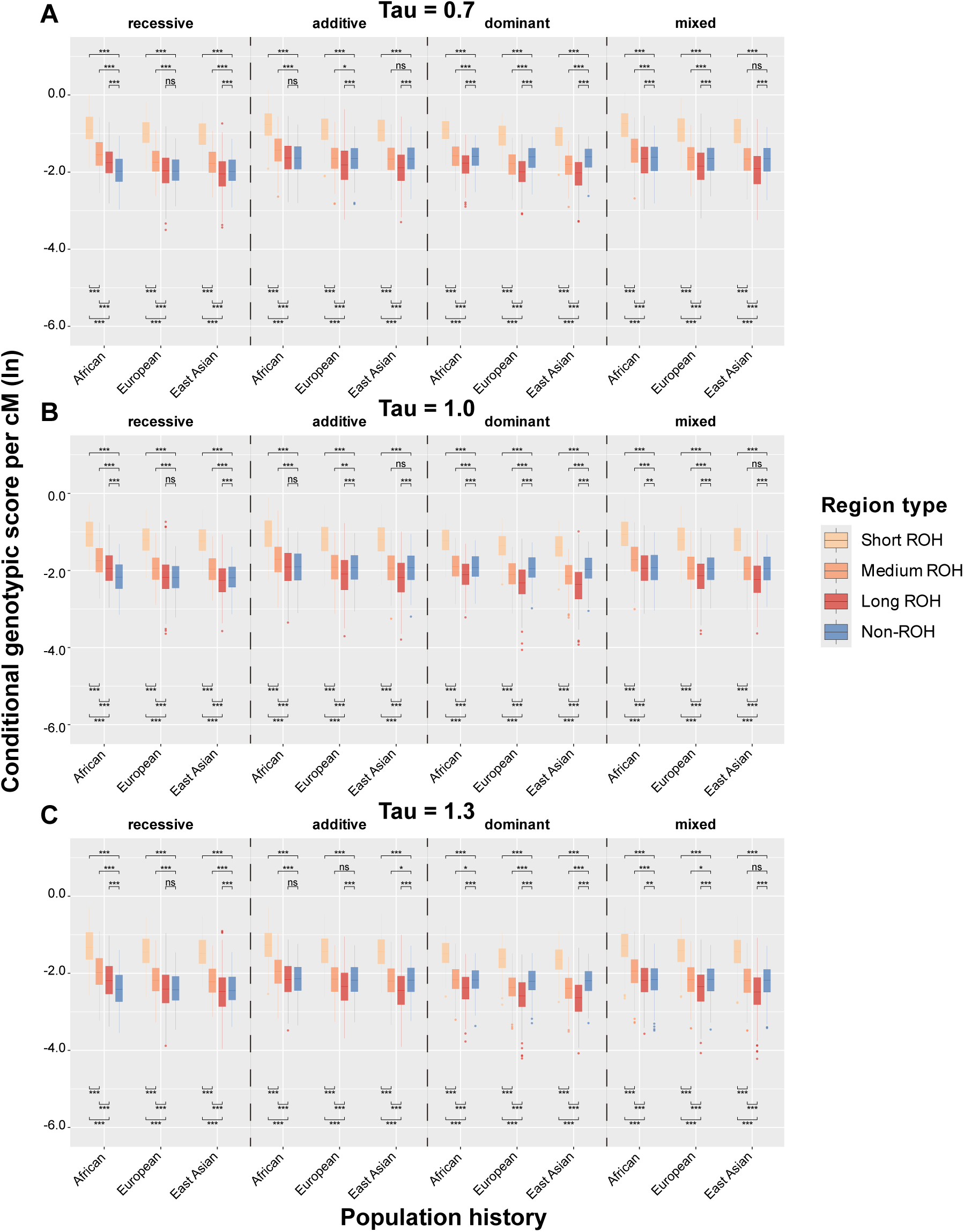
Conditional per-cM genotypic score contribution (log-transformed; neutral mutations) for various regions, population histories, and genetic architectures. Regions are: Short ROH (< 0.25cM); Medium ROH (0.25-1cM); Long ROH (> 1cM); Non-ROH. Genotypic scores are generated from neutral mutations. The parameter tau is varied for different weighting of rare alleles in genotypic score calculation. (A) *τ* = 0.7, (B) *τ* = 1.0, (C) *τ* = 1.3. Significance levels for adjusted *P*-values: *** *P_ad_*_j_ < 0.001, ** *P_ad_*_j_ < 0.01, * *P_ad_*_j_ < 0.05, ns *P_ad_*_j_ ≥ 0.05.

**Figure 5.**
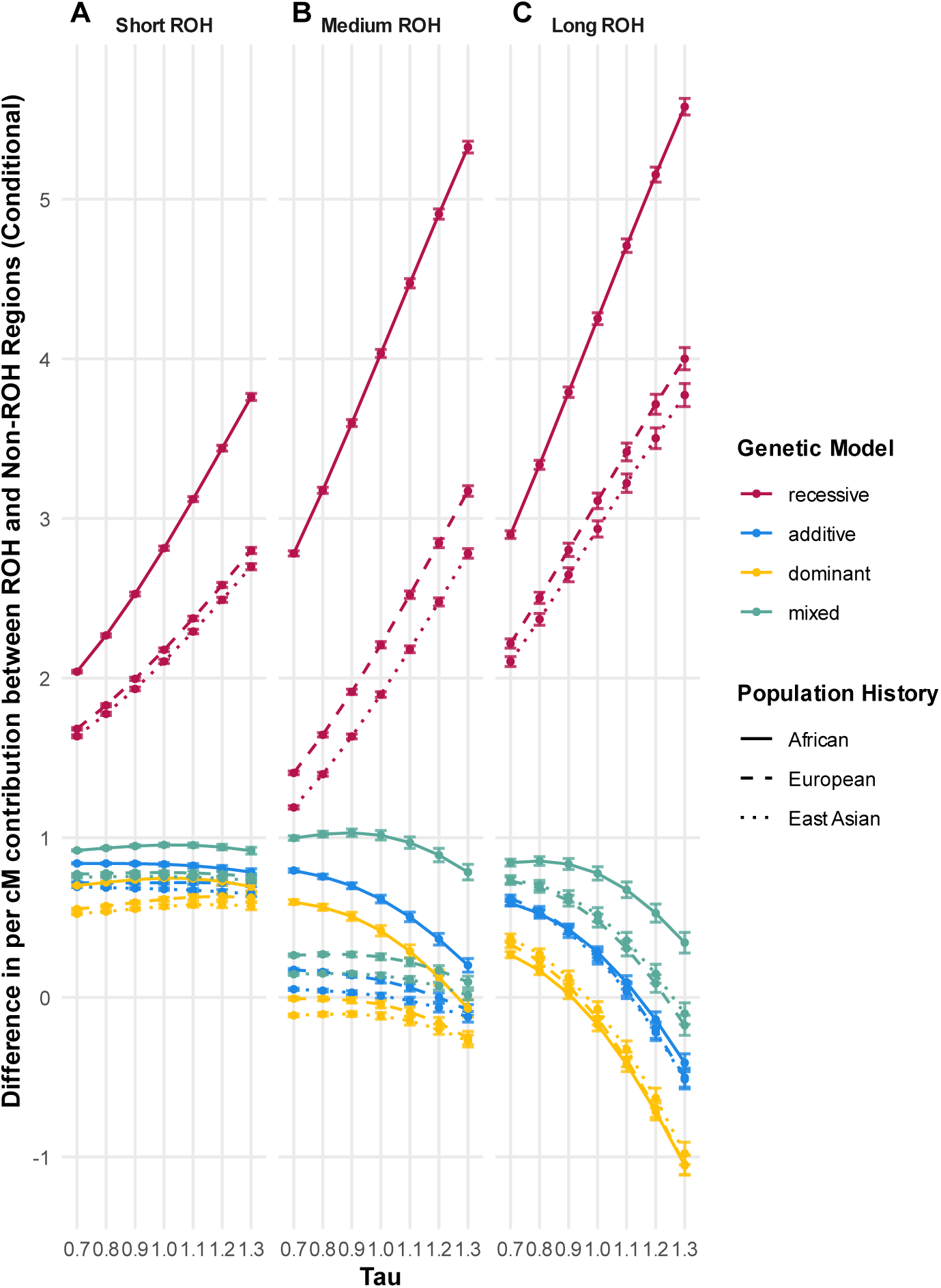
Difference (in log space) in conditional per-cM genotypic score contribution between ROH and non-ROH regions with varied tau, various population histories and genetic architectures. Genotypic scores are only contributed by deleterious mutations. Panels show the difference in contribution between the specific ROH regions and non-ROH regions: (A) Short ROH regions (< 0.25 cM); (B) Medium ROH regions (0.25-1cM); (C) Long ROH regions (> 1 cM).

**Figure 6.**
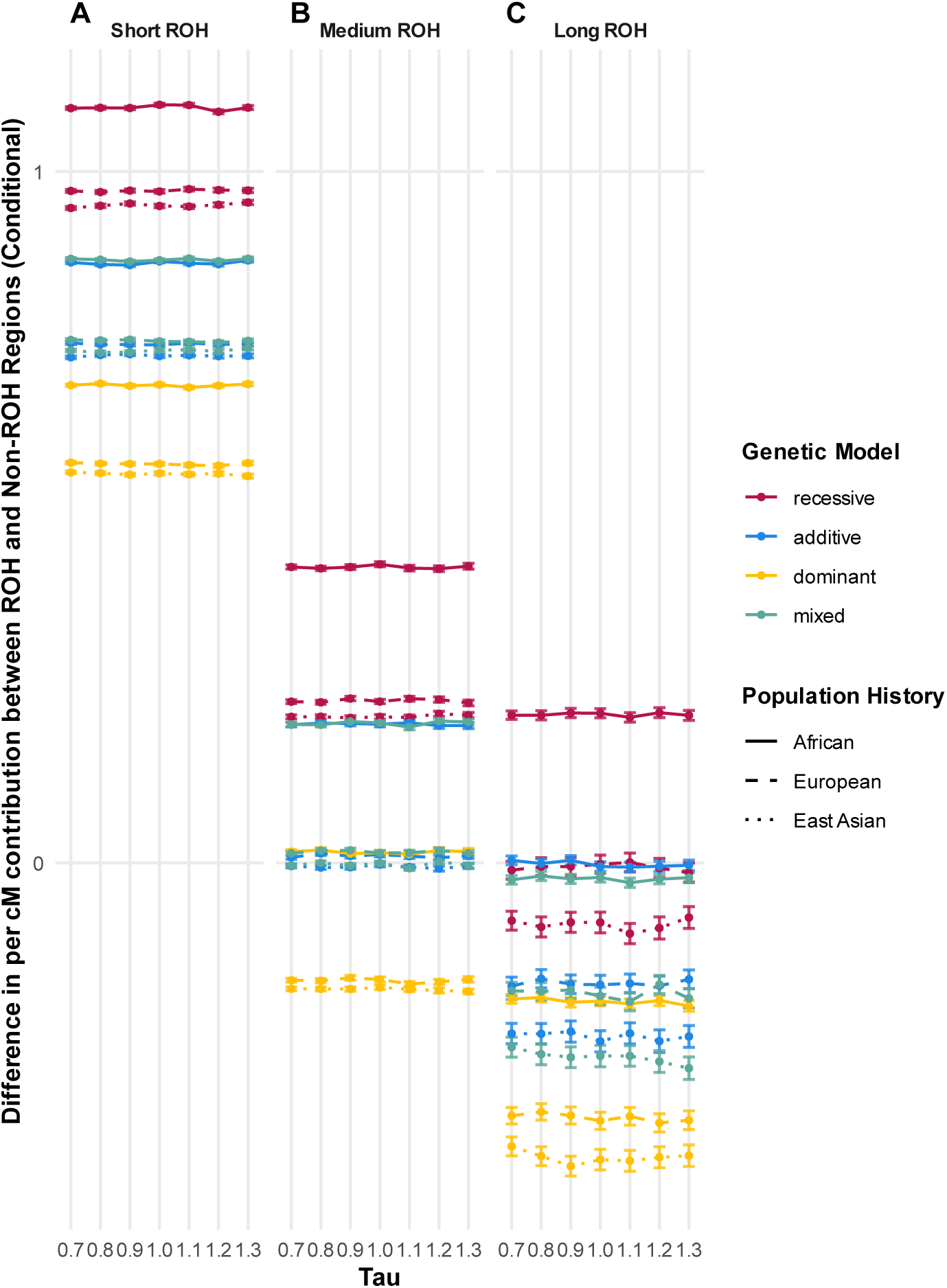
Difference (in log space) in conditional per-cM genotypic score contribution between ROH and non-ROH regions with varied tau, various population histories and genetic architectures. Genotypic scores are only contributed by neutral mutations. Panels show the difference in contribution between the specific ROH regions and non-ROH regions: (A) Short ROH regions (< 0.25 cM); (B) Medium ROH regions (0.25-1cM); (C) Long ROH regions (> 1 cM).

Under the recessive model, all ROH regions in the African demographic history had a significantly higher genotypic score per cM compared to non ROH regions, regardless of the emphasis on common or rare variant (value of tau). Under this model, in the European and East Asian demographic histories, only short ROH and medium ROH regions have a significantly higher genotypic score per cM compared to non ROH regions, regardless of tau. These patterns contrast with the patterns in Figures 2 and S1, where long ROH consistently had the highest genotypic score per cM.

Similarly, under the additive, dominant, and mixed models, short ROH were the only ROH region that consistently had a significantly higher genotypic score per cM compared to non ROH regions across all populations and tau values. Under these models, the genotypic score contribution per cM of long ROH regions were consistently either equal to or less than that of the non-ROH region. However, medium ROH was only significantly higher compared to non-ROH regions in the African population under the additive and mixed models, and this pattern was consistent across tau values.

Notably, the European and East Asian demographic histories showed similar patterns to each other across all models, but they differed from the African demographic history. Specifically, under all models and tau values, the per-cM genotypic score contribution of long ROH in the African population was slightly higher than that in European and East Asian populations (See Supplemental material. Table S1 and S2.). Although the per-cM genotypic score contribution of long ROH was equal to or significantly less than non-ROH regions in all models other than the recessive model.

Figure 8 shows the genotypic score contribution proportion by ROH and non-ROH for phenotypes with neutral effect alleles. The range of contribution proportion for ROH and non-ROH regions was similar to that in the deleterious scenario, with the exception of the recessive case. Not only do ROH contribute proportionally less to the genotypic score under the recessive neutral model compared to the recessive deleterious model (Figure 7), but the proportions do also not change with increasing tau, suggesting that shifting genotypic score contribution from common to rare variants doesn’t influence ROH importance.

**Figure 7.**
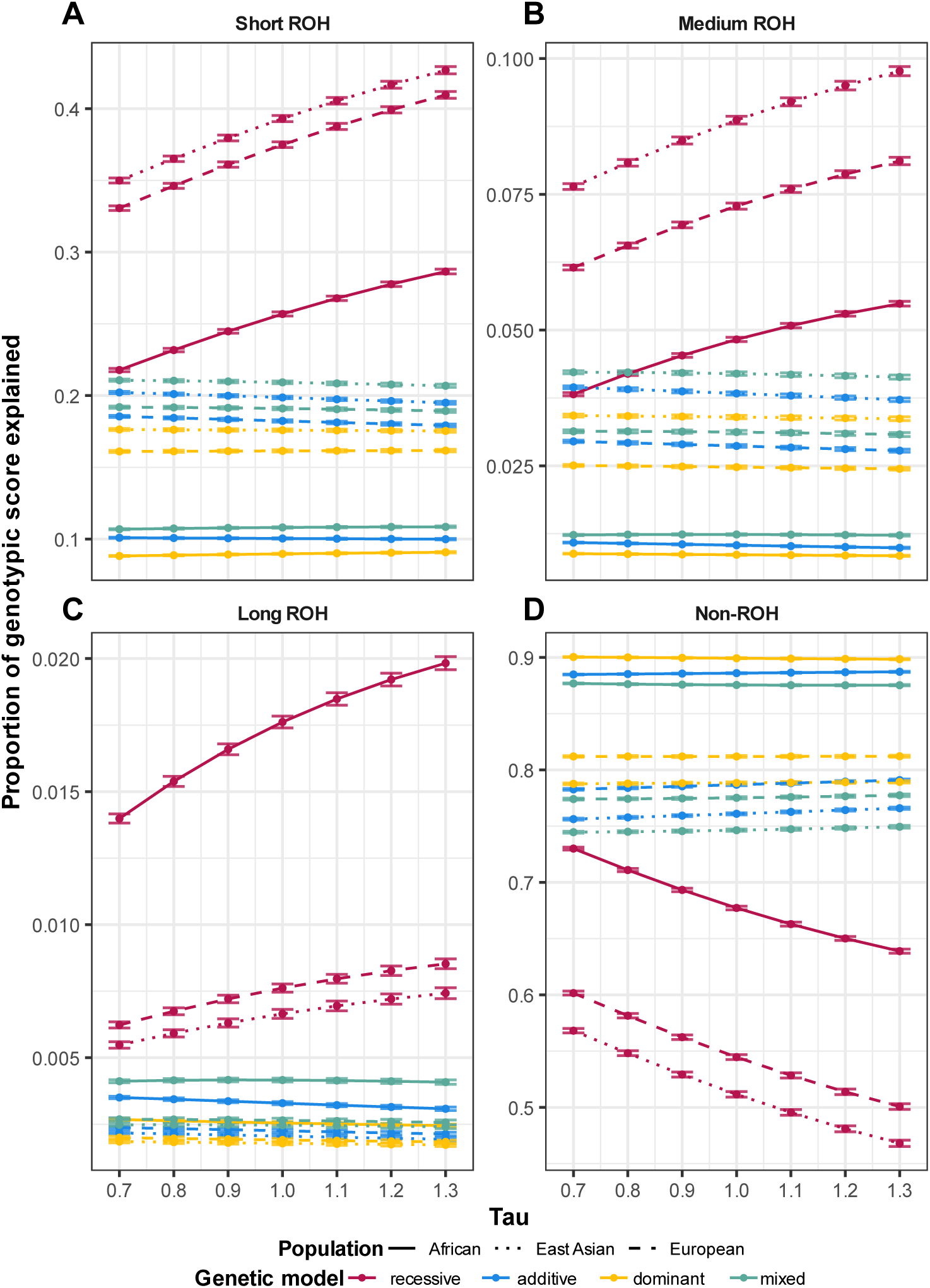
Proportion of total genotypic score per individual explained by different types of regions with varied tau, various population histories and genetic architectures. Genotypic scores are only contributed by deleterious mutations. Panels show the proportion of genotypic score explained by (A) Short ROH regions (< 0.25 cM); (B) Medium ROH regions (0.25-1cM); (C) Long ROH regions (> 1 cM); (D) Non-ROH.

**Figure 8.**
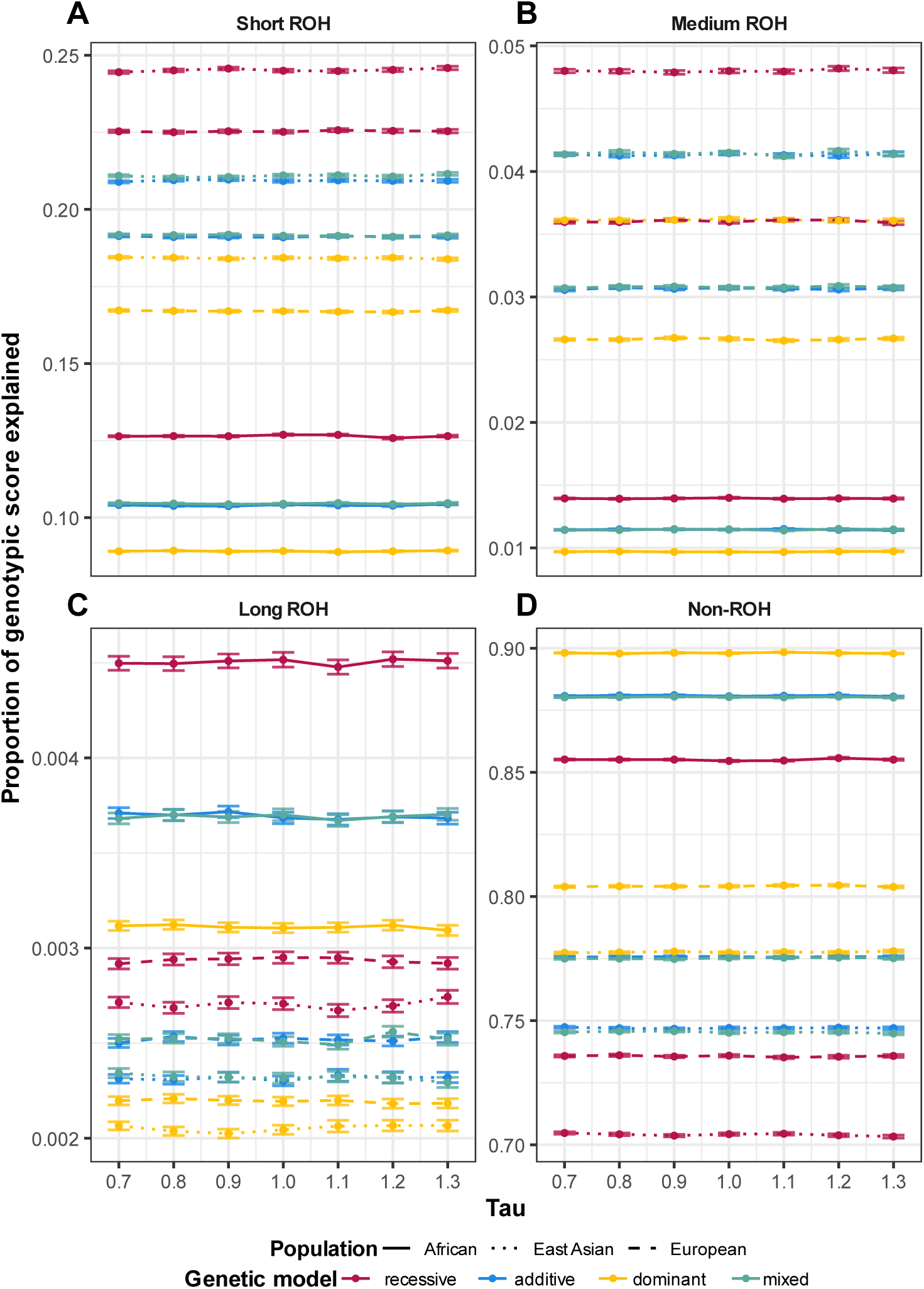
Proportion of total genotypic score per individual explained by different types of regions with varied tau, various population histories and genetic architectures. Genotypic scores are only contributed by neutral mutations. Panels show the proportion of genotypic score explained by (A) Short ROH regions (< 0.25 cM); (B) Medium ROH regions (0.25-1cM); (C) Long ROH regions (> 1 cM); (D) Non-ROH.

Under the neutral model, figure 6 shows that varying tau does not have much effect on the difference between ROH and non-ROH per-cM genotypic score contribution. This is likely due to the fact that under this model the phenotype effect and the selective effect of the allele are decoupled, therefore decoupling the relationship between phenotype effect and allele frequency. However, we still see that under the recessive model ROH generally have larger per-cM genotypic score contribution than non-ROH, although this difference is largest for short ROH and smallest for long ROH. In fact, for long ROH, it is only under the African demography that long ROH have higher per-cM genotypic score contribution compared to non-ROH.

### Robustness analysis

#### Comparison with 1000 Genomes

We considered three scenarios from the 1000 Genomes data: ROH called from genome-wide autosomes, ROH called from the whole chromosome 1, and ROH called from the first 100 Mb of chromosome 1. Our central conclusion is that the abundance and length distribution of ROH segments generated by our simulations are in close agreement with the empirical data.

Figures S3 and S4 show the average number of ROH segments per individual. Results from all three genetic architectures in the simulated data are highly consistent with the empirical data, particularly with the empirical scenario considering only the first 100 Mb of chromosome 1. This is expected, as the number of ROH per individual is positively correlated with the length of the genomic region considered. Figure S5 shows *F_R_*_0*H*_ for all simulated and empirical scenarios. The simulated *F_R_*_0*H*_ values closely match those from the empirical data (AFR: 0.061 in simulation versus 0.072, 0.070, and 0.068 in the three empirical scenarios; EUR: 0.138 (fully additive and fully dominant) or 0.137 (fully recessive) in simulation versus 0.157, 0.154, and 0.154 in empirical scenarios; EAS: 0.163 in simulation versus 0.160, 0.156, and 0.156 in empirical scenarios).

Figures S6 and S7 show the ROH length distributions across all six scenarios; lengths are in cM in Figure S6 and in Mb in Figure S7. For the simulated data, ROH segments were pooled across replicates within each population. In both figures, simulated ROH length distributions closely resemble those from the three empirical scenarios. We observe a strong similarity between East Asian and European distributions, contrasted with a clear difference between these two and the African population. Because of their shared bottleneck history, East Asian and European populations carry more medium and long ROH than the African population. Consequently, their length distributions are shifted toward larger ROH sizes relative to the African population, with peaks located at higher ROH lengths.

Figures S8 and S9 compare the complementary cumulative distribution functions (CCDFs) of ROH lengths between the simulated and empirical data; lengths are in cM in Figure S8 and in Mb in Figure S9. Taking Figure S8 as an illustrative example: as expected, the maximum detectable ROH length in the empirical data decreases with the size of the genomic region considered, and the distribution for the 100 Mb empirical scenario is truncated at approximately 5–10 cM. In contrast, although our simulations are also based on a 100 Mb genomic segment, the long tail of the simulated ROH length distribution closely matches that of the genome-wide autosomal empirical scenario, extending to 20–100 cM at very low frequencies (<10⁻⁶). These very long segments are observable because of the large number of replicates that are simulated (500 per scenario). These results indicate that a 100 Mb simulation with a large number of replicates is sufficient to generate ROH segments comparable in length spectrum to those observed at the full autosomal scale. The three dominance models produce highly similar distributions, indicating that this conclusion is not affected by the choice of dominance model.

### Genetic architecture diagnostics

To assess whether our simulations are biologically reasonable, we conducted the following diagnostics.

#### Sparse model

Under the sparse model, our central conclusions remain robust (Figures S10–S21).

Figures S10 and S11 show the proportion of individuals for sparse and non-sparse model. In the sparse model, each ROH class contains a larger fraction of individuals whose ROH carry zero contribution to the genotypic score. This is an expected consequence of the reduced causal variant density.

Figures S12 and S13 show the conditional per-cM genotypic score contribution for the sparse and non-sparse models. The absolute values decreased, as expected from the reduced causal variant density. However, the relative ordering between ROH and non-ROH regions remains largely unchanged. Under the sparse model, the advantage of all ROH classes over non-ROH regions in the fully recessive model remains robust, with long ROH retaining the highest per-cM contribution under most scenarios. In the other three architectures, the advantage of short ROH over non-ROH regions also remains robust.

Figures S14 and S15 show the unconditional per-cM genotypic score contribution for the sparse and non-sparse models. A similar decrease in absolute values was observed. The relative ordering between ROH and non-ROH regions is nearly identical between the sparse and non-sparse models. Under the fully recessive model, all ROH classes contribute more per cM than non-ROH regions in both the sparse and non-sparse settings. In the other three architectures, short ROH consistently show higher per-cM contributions than non-ROH regions across both models.

For the difference in per-cM genotypic score contribution between ROH and non-ROH regions, both conditional and unconditional analyses show entirely consistent trends and nearly identical values between the sparse and non-sparse models (Figures S16–S19).

Figures S20 and S21 show the proportion of genotypic score explained by ROH in the sparse and non-sparse model. The curves from both models show highly consistent trends and orderings. Specifically, ROH explain 25–55% of the genotypic score in the non-sparse model and 23–50% in the sparse model.

The minor differences between the sparse and non-sparse models are mainly observed in the relative ordering of different ROH classes in per-cM genotypic score contribution. For example, in the African population under the fully recessive model, the conditional analysis shows medium ROH with slightly higher per-cM contributions than long ROH. Similarly, under the unconditional scenario, in the African population under the fully recessive model, when *τ* ≥ 1.0, the non-sparse model yields the ordering long ROH > medium ROH > short ROH, whereas the sparse model yields medium ROH > long ROH > short ROH.

We attribute these fluctuations to two factors. First, the reduced number of individuals meeting the conditional criterion (i.e., those carrying non-zero-contribution ROH) introduces additional statistical noise. Second, medium and long ROH are inherently rare in outbred populations, and their deleterious mutation load is further reduced under the sparse model, generating statistical noise sufficient to perturb the ordering of results. Together, these two factors produce the observed ordering fluctuations under both conditional and unconditional scenarios. These fluctuations do not affect our central conclusions. All specific patterns and orderings are detailed in Figures S10–S21.

#### Allele frequency spectrum and effect frequency spectrum

We observed no obvious differences between the sparse and non-sparse models in either spectrum (Figure S22 vs S23 for AFS; Figure S24 vs S25 for the effect frequency spectrum). This is expected, as sparse-model sampling is fully random. Taking the AFS under the non-sparse model (Figure S23) as an illustrative example: under the same genetic architecture, the European and East Asian populations, which experienced the shared bottleneck, exhibit an excess of rare variants compared to the African population. This is consistent with the expected rare variant excess following a population bottleneck. Under the same population history, the distribution under the fully recessive scenario contains more alleles at medium to high frequencies and fewer alleles at low frequencies. This is because, in the three non-recessive architectures, selection acts even on heterozygotes, making it more difficult for alleles to reach high frequencies. For the same reason, the effect frequency spectrum under the non-sparse model (Figure S25) shows that, under the same population history, the fully recessive scenario exhibits elevated frequencies at medium-to-high effect sizes. We also note that increasing τ does not alter the shape of the effect frequency spectrum but only rescales the effect sizes. This is also expected, as a foundational assumption of the Eyre-Walker model is that selection acts on fitness rather than on phenotype directly, unless the phenotype is fitness itself.

#### Diagnostic of causal alleles

For both the proportion of causal variants in ROH regions and the proportion of homozygous causal variants in ROH regions, we observed no obvious differences between the sparse and non-sparse models (Figure S26B vs Figure S27B; Figure S26D vs Figure S27D). This is expected, as sparse-model sampling is fully random and should not affect proportions.

For both the average number of causal variants and the average number of homozygous causal variants per individual, we observed a corresponding decrease in the sparse model due to the random down-sampling (Figure S26A vs Figure S27A; Figure S26C vs Figure S27C). Taking the non-sparse model as an illustrative example: under the same genetic architecture, the African population carries more causal variants than the European and East Asian populations (Figure S27A). This is expected, given the higher genetic diversity in the African population. Under the same population history, the fully recessive scenario shows the highest number of causal variants and the fully dominant scenario shows the fewest. This is because selection acts most efficiently under the fully dominant scenario. For homozygous causal variants, under the same genetic architecture, the European and East Asian populations carry more homozygous causal variants than the African population (Figure S27C), reflecting the homozygotes produced by their shared bottleneck history.

For the proportion of causal variants located in ROH regions, we first observed that, regardless of population history or genetic architecture, this proportion is always higher than *F_R_*_0*H*_ (i.e., the genomic fraction covered by ROH). This indicates that causal variants are not randomly distributed across the genome. Second, under the same genetic architecture, the European and East Asian populations show higher proportions than the African population. This is because they carry more ROH segments. In contrast, under the same population history, the proportions across the four genetic architectures are nearly identical. This is because different genetic architectures only change the total number of causal variants, without altering their distribution between ROH and non-ROH regions.

For the proportion of homozygous causal variants located in ROH regions, under the same population history, the fully recessive scenario shows the highest proportion and the fully dominant scenario shows the lowest. The underlying mechanism likely involves the interaction of several evolutionary forces, but one possible explanation operates at the haplotype level. ROH arise when two IBD haplotypes co-occur in an individual. Under the fully recessive scenario, deleterious mutations on these haplotypes are not exposed to selection in the heterozygous state, allowing more deleterious variants to accumulate on common haplotypes. When two such haplotypes pair up to form a ROH, the deleterious variants they carry are simultaneously homozygous. In contrast, under the fully dominant scenario, selection already acts on heterozygotes, so deleterious variants on these haplotypes have been more efficiently purged, leading to a lower enrichment of homozygous causal variants within ROH.

#### Genetic variance explained by rare versus common alleles

For the proportion of genetic variance explained by rare versus common variants, we observed no obvious differences between the sparse and non-sparse models (Figure S28 vs Figure S29). This is expected, as sparse-model sampling is fully random and should not affect proportions.

Taking the non-sparse model as an illustrative example: under the fully recessive scenario, variants with allele frequencies ≥1% explained the majority of the genetic variance, with this proportion slightly decreasing as τ increased. The remaining genetic variance was explained primarily by variants with frequencies between 0.1% and 1%, and this proportion slightly increased as τ increased. In contrast, under the other three architectures, variants with frequencies below 0.1% explained the majority of the genetic variance, with this proportion increasing as τ increased.

These results are consistent with our expectations: τ behaves as intended. Moreover, under the fully recessive scenario, the choice of τ values did not produce an extreme outcome in which rare variants explain the vast majority of the genetic variance. For the other three architectures, although less common variants explained the majority of the genetic variance, this is not a consequence of the τ values chosen. Rather, as shown in Figures S22–S25, purifying selection is highly efficient in these three architectures: a variant has difficulty rising from low-frequency rare status to common-variant status at medium or high frequencies, and these architectures rarely accumulate variants with high effect sizes. We therefore consider our choice of τ range to be reasonable.

#### Genetic variance explained by ROH regions

For the proportion of genetic variance explained by ROH regions, we observed no obvious differences between the sparse and non-sparse models (Figure S30 vs Figure S31). This is expected, as sparse-model sampling is fully random and should not affect proportions. Under the fully recessive scenario, ROH regions explained the majority of the genetic variance. This is expected, as deleterious variants under the fully recessive scenario contribute to the genotypic score only when homozygous. In contrast, under the other three scenarios, non-ROH regions explained the majority of the genetic variance. This is primarily because deleterious variants under these three scenarios already contribute in the heterozygous state.

#### Additive and dominance variance

For the proportions of additive variance and dominance variance within the total genetic variance, we observed no obvious differences between the sparse and non-sparse models (Figure S32 vs Figure S33; Figure S34 vs Figure S35). This is expected, as sparse-model sampling is fully random and should not affect proportions.

Taking the non-sparse model as an illustrative example (Figure S33): under the fully recessive scenario, dominance variance accounts for approximately 75% of the genetic variance. This is expected, as all causal variants are fully recessive. For the fully additive scenario, the proportion of dominance variance is zero (Figure S34). For the remaining two architectures, the proportion of dominance variance is also very low. These results are consistent with our expectations, as they directly reflect the dominance coefficients simulated for deleterious variants in each scenario.

#### GARLIC versus GERMLINE

As shown in Figure S36, the conditional per-cM genotypic scores under the fully recessive scenario obtained from GARLIC and from GERMLINE are highly consistent. In European and East Asian populations, the contribution values and orderings of ROH classes are identical between the two methods. In the African population, the contribution values and orderings of ROH classes are also nearly identical. The only difference is that, when τ < 1, the value of medium ROH is slightly higher than that of long ROH; when τ ≥ 1, the values of medium and long ROH become equal. We attribute this more likely to statistical noise between the two methods than to a biologically meaningful difference. We therefore consider GARLIC to be a reliable ROH caller for our purposes.

#### cM versus Mb

As shown in Figure S37, the conditional per-unit genotypic scores under the fully recessive scenario obtained using cM and Mb as units are highly consistent. In the African population, the contribution values and orderings of ROH classes are nearly identical between the two unit choices. In the European and East Asian populations, the contribution values and orderings of ROH classes are also nearly identical between the two unit choices. The only difference is that, when τ < 1, the value of medium ROH is slightly higher than that of short ROH when Mb is used as the unit. We attribute this more likely to statistical noise between the two unit choices than to a biologically meaningful difference. We therefore consider our conclusions to be robust to the choice of Mb as the measurement unit.

As shown in Figure S38, under the fully recessive scenario, the length distributions of short and medium ROH across the three populations are nearly identical between cM and Mb. For long ROH, the distributions across the three populations are more similar to each other when measured in cM, whereas in Mb the African population shows slightly higher long ROH lengths than the European and East Asian populations. This difference between the two unit choices likely reflects measurement noise introduced by recombination rate heterogeneity in Mb-based units.

## Discussion

Runs of homozygosity (ROH), as widely observed genomic features, are gradually emerging as a key factor in understanding the genetic basis of complex traits.^7, 28, 40, 42, 43, 48^ However, the specific mechanisms by which ROH influence complex trait phenotypes, particularly how these mechanisms are systematically modulated by the combined effects of population history and genetic architecture, remain largely unknown. To address this gap, our study systematically investigated the interactions between these factors using large-scale realistic simulations. Our results reveal that, under a selection-coupled architecture, the contribution of ROH to complex traits genotypic score is important but variable. Under the recessive model, ROH, especially long ROH, play a significant and central role, but under other genetic architectures, the contribution patterns differ based on the underlying model.

### Proportion of individuals with ROH

As observed in Figures 1 and 3, many individuals lack ROH regions that contribute to genotypic score. While overall levels of ROH are generally determined by the demographic history, the contrast between Figures 1 and 3 indicate that when the phenotype is determined by deleterious alleles, ROH that have a phenotype effect are subject to purging. Importantly, functional alleles were simulated only within exon regions, which induce a clustering of functional alleles along the genome. If intronic and intergenic alleles can also contribute to phenotypes than we may expect ROH to contain further enrichments of functional variation beyond coding variation. As the three populations we simulated followed random mating with only two populations experiencing a historical bottleneck, long ROH were relatively rare across populations. However, we note that modern human populations are more complex than these demographic models, and founder effects,^8^ recent bottlenecks,^49^ and localized fine structure^50^ can influence long ROH levels in the genome.

Previous studies have focused on the relationship between the total length of ROH (sROH) and total number of ROH (nROH) of sampled individuals in relation to phenotypes.^48^ Our study shows that ROH, especially the long ROH with non-zero genotypic score contribution, exist in only a small proportion of individuals. If we combine the genotypic score contributed by these ROH across all sampled individuals, it will inevitably lead to substantial downward bias in the relationship between ROH and phenotype. We propose that future work on the relationship between ROH and phenotype attempt to model these two types of ROH, those with no genotypic score contribution and those with genotypic score contribution.

### The recessive model

Among all the genetic architectures tested, ROH contribute the most to the complex trait genotypic score under the recessive model. Under this model, our results demonstrate significant cross-population applicability: regardless of population history, in all the populations, the per-cM genotypic score contribution of all ROH regions, and especially long ROH, is systematically higher than that of non-ROH regions (Figure 2, S1). Furthermore, this pattern exacerbates when the weight of rare-allele contribution to genotypic score is increased (Figures 2, 5, S1).

The core mechanism behind this lies in the difference in the probability of recessive alleles forming homozygotes in different genomic regions. In non-ROH regions, for an effect allele with frequency *p*, homozygotes are expected to form at a rate proportional to *p*^2^. However, within ROH regions homozygotes are expected to form at a rate proportional to *p*, and, for a rare allele, *p* ≫ *p*^2^. Furthermore, changing the weight of rare-allele contributions to the phenotype underscore this pattern. As rare alleles become relatively more important contributors to genotypic score (increasing tau), the difference between the per-cM genotypic score contribution of ROH regions compared to non-ROH regions increases, with long ROH increasing the most (Figure 5). Under the recessive model with deleterious effect alleles, only homozygotes are exposed to negative selection, and it is these factors that drive the significant differences between the per-cM genotypic score contribution between ROH and non-ROH regions.

These mechanisms also explain the pattern that long ROH have a significantly higher per-cM genotypic score contribution than medium and short ROH. This is closely related to the age of the underlying haplotype and the effect sizes of deleterious mutations they carry. Long ROH are comprised of young haplotypes and are the direct result of recent inbreeding or bottlenecks and other recent evolutionary events. Due to their age, natural selection has not had time to eliminate any deleterious mutations they may carry. This leads to long ROH having a higher chance of being enriched for the deleterious mutations with high effect. Thus, they have the highest per-cM genotypic score contribution. In contrast, the older short and medium ROH are comprised of haplotypes that have segregated for a longer time, with more opportunity to be exposed to negative selection. As a result, deleterious mutations with strong effects would be mostly purged by natural selection, resulting in their relatively low per-cM genotypic score contribution.

However, when we shift focus from per-cM genotypic score contribution to genotypic score contribution proportion (Figure 7), the order of contribution of ROH regions is changed, with short ROH > medium ROH > long ROH. This is mainly because we have simulated outbred populations and long ROH are relatively infrequent and generated due to small effective population size only. Indeed, long ancestral haplotypes are constantly broken up by recombination events, and thus ROH are biased toward being comprised of shorter haplotypes that have persisted to the present day. Therefore, though the per-cM genotypic score contribution of short ROH is the lowest among all the ROH categories, their large number in the genome allows them to have the highest genotypic score contribution proportion. Although long ROH have the highest per-cM genotypic score contribution, they are rarer and younger and thus have the lowest genotypic score contribution proportion (Figure 3). However, these patterns also highlight that, in the presence of a harsh bottleneck or increased consanguinity, increased prevalence of long ROH could account for a substantial proportion of a phenotype. Indeed, these patterns and expectations likely explain the importance of ROH in highly endogamous populations.^41, 51–53^

### The additive and dominant models

When the genetic architecture is changed from the recessive model to the additive and dominant models, the patterns of ROH contribution to genotypic score is significantly altered, with differences observed in per-cM genotypic score contribution, genotypic score contribution proportion, and its response to changing rare-allele weight (tau value).

Whereas, under the recessive model, ROH had higher per-cM genotypic score contribution compared to non-ROH regions, under the additive and dominant models, the pattern of per-cM genotypic score contribution of ROH regions is reversed, with short ROH regions typically showing the most prominent effect (Figures 2, S1). More importantly, under these two models, as the tau value increases, only short ROH regions consistently show a higher effect per-cM over non-ROH regions (Figure 5).

We propose these patterns are due to the fundamental differences in the efficiency of natural selection under different genetic architectures. Under the additive and dominant models, regardless of zygosity, the deleterious effect of mutations is exposed to the pressure of natural selection, resulting in the elimination of mutations with strong effect. In contrast to the recessive model, long ROH, which are comprised of “young” haplotypes, cannot effectively accumulate a large number of deleterious mutations with high effect size because they are efficiently eliminated by selection under the additive and dominant models, leaving only weak effect alleles to persist. Hence, the per-cM genotypic score contribution of long ROH regions exhibits significant decline compared to those under the recessive model. As the ancient haplotypes that comprise short ROH survived in the evolutionary process, the persistent existence of short ROH may itself imply certain characteristics of the regions in which they are located, such as a slightly lower recombination rate. In population genetics, the lower recombination rate decreases the efficiency of natural selection in eliminating weak deleterious mutations. Therefore, these short ROH regions have accumulated more weak deleterious mutations over a long period of time, which may explain the higher per-cM genotypic score contribution compared to long ROH and non-ROH regions.

Notably, the relative per-cM genotypic score contribution of long and medium ROH is dependent on the tau value. When tau is small, emphasizing common variants, the long and medium ROH have higher per-cM genotypic score contribution than non-ROH regions across all populations under additive and dominant models, but as tau increases, emphasizing rare variants, this advantage reverses. Thus, when tau is low, the genotypic score of long and medium ROH is highly dependent on the common alleles they carry. When tau increases, because they lack the long-term accumulation of rare alleles, their genotypic scores decrease sharply. Although non-ROH also experience a decline in genotypic scores due to the increasing tau, their genetic diversity allows rare variants to buffer the decline in genotypic scores. It is precisely this difference in vulnerability to changes in tau that leads to the reversal of contribution differences between long/medium ROH and non-ROH. However, we also noticed an exception: under the dominant model, the medium ROH in European and East Asian populations didn’t show the advantage in per-cM genotypic score contribution over non-ROH even when tau is small. This significant population-specific pattern is discussed in a subsequent section.

Additionally, we observed that the value of tau for which longer ROH explain less genotypic score than non-ROH under the dominant model is lower than that under the additive model. This is likely because of the different efficiency of natural selection under these models. Under the dominant model, the full effect of a deleterious allele is fully exposed in heterozygotes, which leads to the higher purging efficiency. Thus, the genotypic score per-cM under the dominant model is more sensitive to tau values.

The additive and dominant models show consistent patterns with increasing tau leading to a decrease in per-cM genotypic score contribution for both ROH and non-ROH regions, which is in contrast to the recessive model. On the other hand, the genotypic score contribution proportion of ROH regions under the additive and dominant models is not only lower than that under the recessive model, but also remains generally unchanged with the increasing tau. The former is because the deleterious heterozygotes in non-ROH regions can contribute to genotypic score too, thereby increasing the total genotypic score and decreasing the proportion of ROH regions. The latter is because, under these two models, as shown in Figure 5, the relative per-cM genotypic score contribution between ROH and non-ROH regions does not change much. Therefore, their genotypic score contribution proportion remains stable even when tau increases. These results clearly demonstrate that the contribution of ROH regions to polygenic genotypic scores is highly dependent on the underlying genetic architecture of the trait.

### Population history: a key factor influencing ROH genotypic score contribution paflerns

Population history is another factor that influences the ROH genotypic score contribution patterns. Based on the classic out-of-Africa model, we systematically investigated its effect on “African”, “European”, and “East Asian” populations. These effects are mainly reflected in two aspects: per-cM genotypic score contribution and the genotypic score contribution proportion of each ROH and non-ROH region.

First, regarding the per-cM genotypic score contribution, the results under the recessive model stand out the most. The per-cM genotypic score contribution of ROH regions in the African population is significantly higher than those in the European and East Asian populations (Figure 2), while the European and East Asian populations have similar patterns. These results directly reflect the distinct population histories. Due to its long-term stable effective population size, the African population has the highest genetic diversity. This includes more rare deleterious mutations that are masked as heterozygotes but can potentially be enriched as homozygotes when paired identical-by-descent within ROH. In contrast, European and East Asian populations experienced a historical bottleneck event, which greatly reduced their genetic diversity and likely induced purging of strong deleterious alleles.

These patterns are exacerbated when changing tau to modify the relative influence of rare and common alleles. We observed that under the recessive model, with an increase in tau values, the relative contribution of ROH to the genotypic score compared to non-ROH regions increases significantly more for African populations compared to European and East Asian populations (Figure 5). This underscores our expectation that the African population has a more abundant pool of rare variants. This means when the weight of rare alleles in genotypic score calculation increases, compared to the other two populations, the ROH regions in the African population are enriched for rarer/more deleterious variation.

Notably, an interesting pattern arises under the dominant model when examining the influence of population history. In the African population, when tau is low, medium ROH have significantly higher per-cM genotypic score contribution compared to non-ROH regions (Figure 2). However, across all tested tau, in the European and East Asian populations these ROH have significantly less per-cM genotypic score contribution compared to non-ROH regions. This pattern likely reflects the population bottlenecks experienced by these two populations, along with the accompanying founder effect, may have created a preponderance medium ROH that carry a lower-than-average load of dominant deleterious mutations.

Population history also shows a notable influence under the mixed dominance model. The mixed model is special because it includes three types of deleterious mutations with different genetic structures in proportion, approximately 88.4% additive, 7.1% recessive, and 4.5% dominant (see Methods). As noted in the Methods, the mixed model is a heuristic scenario rather than a definitive account of the true dominance landscape; its results should therefore be interpreted as exploratory. However, even though additive mutations have such a high proportion, the per-cM genotypic score contribution of ROH and non-ROH regions is only similar to those under the additive model in European and East Asian populations, but in the African population they similar to the recessive model (Figure 2). Once again, this is likely the result of high genetic diversity of the African population with many rare recessive alleles masked as heterozygotes, that get paired into homozygotes within ROH. In contrast with the European and East Asian populations for which many of these alleles were purged during their bottlenecks. Therefore, although the proportion of recessive alleles is low, in the African population, the impact they produce on genotypic scores when enriched in ROH is so great that their overall effect is disproportionately amplified, even enough to override the effects of additive mutations that dominate in terms of absolute numbers.

Finally, population history also has a consistent influence on the proportion of genotypic score explained by different ROH regions. Specifically, we observed that the proportion of genotypic score explained by short and medium ROH under all the models follows the order of East Asian > European > African (Figure 7A-B). However, the genotypic score contribution proportion of long ROH shows the reverse pattern, African > European > East Asian (Figure 7C). The pattern exacerbates when tau increases. These results highlight the importance of long ROH for explaining phenotypes in populations of high genetic diversity.

### Phenotypes from neutral mutations as a comparison

To contextualize the results from above where the phenotype is derived from the fitness effects of individual mutations, we also considered a scenario where neutral mutations were the causal alleles as a reference. In this scenario, when we calculate the genotypic score of neutral mutations, the effect size of mutations is sampled from the marginal distribution of selection coefficients while the actual selection coefficient of the mutations remains 0. To control for any effects of background selection, we continue to simulate deleterious alleles in these cases, but they do not have any phenotype effect. Therefore, in Figure 3, the observation that all the individuals from the European and East Asian populations have at least one medium ROH with non-zero genotypic score is due to the bottleneck event and the random distribution of neutral mutations in exons. The random distribution also explains the increased proportion of individuals with non-zero contribution long ROH across all three populations, as well as the increased proportion of individuals with non-zero medium ROH from the African population.

Furthermore, under the neutral scenario, compared to non-ROH regions, the higher per-cM genotypic score contribution observed in ROH regions (especially long ROH) under the recessive model disappeared. Similarly, all the key patterns observed in the deleterious model, such as the advantage of long ROH, the distinct patterns of African populations in mixed models, or the systematic effects of tau values, are either completely absent or greatly weakened in the neutral mutation scenario (Fig 4, 6, 8). These contrasts strongly demonstrate that the observations we obtained from deleterious scenarios are the results of the combined effects of natural selection, genetic architecture, and population history.

### Limitations and Future Directions

Although these simulations are designed to be realistic, this study still has several limitations. Although this study systematically considered multiple different genetic architectures, our model did not include epistasis or gene-environment interactions (GxE^54, 55^), which are wide-spread in biology. The out-of-Africa model we used is widely used in forward-time simulations, but it is also an idealized and simplified model that does not fully capture the more complicated migration and admixture events in the actual human evolution history. In this case, the complicated evolutionary history^56^ underlying the real human genomic data may further influence the distribution of ROH and their genotypic score contribution patterns. Explicitly varying individual demographic parameters in simplified toy models would help disentangle the contributions of specific demographic events and remains a direction for future work. The genomic structure we simulated in our model is based on the first 100 Mbps of human chromosome 1, which cannot fully represent the entire human genome. Although our comparison with the 1000 Genomes data (Figures S3–S9) demonstrates that this region reproduces genome-wide ROH distributions, exploring additional genomic regions beyond the first 100Mbps of chr1 remains worthwhile. Furthermore, we only simulate causal alleles within exon regions and assume all alleles contribute to the genotypic score. All simulated populations are outbred; incorporating explicit consanguinity would extend our framework to inbred populations. Additionally, sampling dominance coefficients from a continuous distribution and exploring a broader range of recessive fractions for mixed model remain promising directions. However, despite these limitations, we believe that the core conclusions of our study, that ROH regions contribute robustly and significantly to polygenic genotypic score under different genetic architectures and population histories, particularly in recessive models and populations with high genetic diversity, remain valid.

Our study provides new mechanistic insights into existing work in the field. Previous studies have found that long ROH can enrich deleterious mutations.^42, 43^ In this project, we have extended their work to broader systems with more complicated genetic architectures and population histories. In previous studies, ROH has often been used to investigate its association with diseases such as schizophrenia^4, 28, 29, 37^ and Alzheimer’s disease.^28, 57–59^ However, these associations have shown inconsistent conclusions across different populations,^7, 60–63^ leading to doubts about the utility of ROH. Our work shows that such inconsistencies are actually expected, as the effect of ROH is closely related to the population history of the sampled population and the genetic architectures of the trait.

In the future, the conclusions and observations from this study can be further verified in real human genome data. At the same time, the sensitivity of ROH to evolutionary history suggests its potential to be incorporated into inference methods. We also believe that the contribution patterns of ROH, especially the long ROH under the recessive model and short ROH under other models, can be helpful for method development in finding rare variants in genome data and exploring the missing heredity. In addition, future simulations can build upon our work to integrate more biological realities to explore the role of ROH in more realistic genetic landscapes.

This study systematically shows that, under a selection-coupled architecture, ROH in the human genome are an underestimated but powerful source for understanding the genetic basis of complex traits. The significance of ROH goes far beyond that of an inbreeding indicator. ROH are dynamic features shaped by population history, whose function is defined by the genetic structure of traits. Our study demonstrates that the polygenic genotypic score contribution of ROH is the result of both natural selection and the accumulation of deleterious mutations, with the relative importance of these two mechanisms determined by both genetic architecture and population history.

## Methods

### Simulating Genetic Data

We simulated genetic data using SLiM 4.1^64^ and msprime^65^ under a three-population Out-of-Africa demographic model from Gravel et al.^66^ (for demographic parameters see SLiM recipe 5.4). For a realistic genome structure, we simulated 100 Mbps genome segment with an exon structure based on the CCDS^67^ coordinates (GRCh37) from the first 100 Mbps of human chromosome 1. We used a variable recombination rate map based on the HapMap3^68^ recombination map. Deleterious mutations were restricted to exon regions; thus, the deleterious mutation rate was set to 2.36×10^-9^, which is 10% of the mutation rate of 2.36×10^-8^ for the entire region. Deleterious selection coefficients were sampled from a gamma distribution with shape 0.2 and mean –0.03.^69, 70^ We considered four different scenarios for dominance: fully recessive (h = 0), fully additive (h = 0.5), fully dominant (h = 1), and a mixed model including all three. All simulated populations follow random mating without consanguinity; this study therefore focuses exclusively on outbred populations.

To determine the relative proportions of recessive, additive, and dominant mutations in the mixed dominance model, we proceeded as follows. First, we downloaded the GRCh37 gene annotation file^71, 72^ and restricted it to only autosomal genes. Next, we downloaded two gene lists^73, 74^ generated by the MacArthur group (https://github.com/macarthur-lab/gene_lists) corresponding to genes that are deemed to follow autosomal dominant inheritance or recessive inheritance based on Online Mendelian Inheritance in Man (OMIM) database. We downloaded a third list (“universal”^75^) and assumed that genes present in the universal list but absent from both the dominant list and the recessive list can be classified as additive. For each gene, we calculated the maximum transcript length across the different transcript versions. Then, we summed the total transcript length among all genes classified as dominant, recessive, or additive, and then divided by the total transcript length of all the genes considered. For the simulations, the relative proportion of dominance coefficients for new mutations is given by these proportions, calculated as dominant:additive:recessive = 0.04549:0.8839:0.07059. We note that OMIM inheritance-pattern annotations at the gene level are not directly equivalent to dominance coefficients at the variant level. The mixed model is therefore treated as a heuristic scenario, constructed from currently available biological information, rather than an exact representation of the true variant-level dominance coefficient distribution in humans, which remains unknown.

To speed up our simulations, we did not use SLiM to simulate neutral mutations. Instead, we generated tree sequences and used msprime to recapitate and overlay neutral mutations at a mutation rate of 2.124 × 10^•8^. For each population, we sampled 500 diploid individuals, and we simulated 500 replicates per parameter combination.

### Simulating Phenotypes

We simulated the genetic component of phenotypes under two separate scenarios, one where causal mutations are deleterious and one where causal mutations are neutral. For phenotypes generated from deleterious mutations, we assumed all deleterious mutations contribute to the genotypic score. We used a quantitative model derived from Eyre-Walker, Adam ^76^and Uricchio, Lawrence H et al,^44^ which explicitly relates the selection coefficient of the mutation to an effect size. We compute genotypic scores for each individual to represent the genetic contribution to the phenotype. For a given selection coefficient, *s*, the genotypic score *Z_s_* is given by

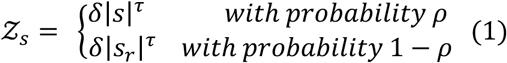

In this model, *δ* and *τ* allow the marginal distribution of effects to differ from the marginal distribution of selection coefficients. As we are interested in directional effects, we set *δ* = 1. When *τ* increases, it up-weights the significance of rare-allele contributions to genotypic scores, and, when *τ* decreases, it up-weights the significance of common-allele contributions to genotypic scores. The parameter *ρ* allows for the introduction of pleiotropy without altering the overall marginal distribution of effects, as *s_r_* is generated from a new draw from the marginal distribution of selection coefficients. For phenotypes generated from deleterious alleles we set *ρ* = 1. For each sampled individual, the total genotypic score, *Z,* was computed as follows.

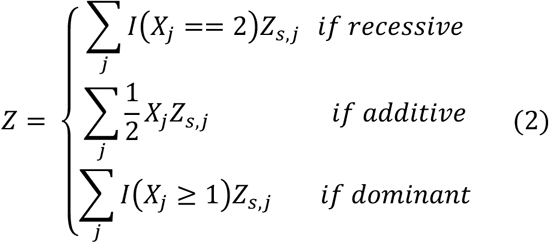

where *X*_j_ is the number of effect alleles at locus j, and *I*(*A*) is an indicator function that takes the value 1 when proposition A is true and 0 when false. The sum is across all effect loci. The inheritance pattern of the phenotype effect (i.e., recessive, additive, or dominant), is determined by the dominance coefficient of the selection coefficient. We considered *τ* ∈ 0.7, 0.8, 0.9, 1.0, 1.1, 1.2, 1.3} to examine the influence of changing the significance of rare alleles.

For phenotypes generated from only neutral alleles, we used the above phenotype model but set *ρ* = 0, so that we randomly drew effects from the marginal distribution of selection coefficients and transform accordingly. However, as there are many more neutral alleles than deleterious ones in the genome, we down sampled the number of neutral effect alleles that contribute to the genotypic score as follows.

For a given parameter combination and dominance regime, we calculated the distribution of the total number of deleterious mutations for all populations combined across simulation replicates. We fit a normal distribution and employed the Shapiro-Wilk test for testing goodness of fit. The results of this statistical test confirmed that the data fits well. (See Supplemental material. Table S7) Then, we sampled from this distribution to determine the number of neutral effect alleles for each replicate with a matching parameter and dominance regime. Using this number of effect alleles, we randomly choose neutral mutations that fall within exons and assign an effect size using the above model. Although most loci are biallelic, occasionally a locus with >1 derived allele is encountered. In this case, we randomly choose a derived allele at that locus to be the effect allele. Importantly, the dominance of phenotype for neutral alleles follows the dominance pattern of the deleterious alleles that were simulated. In the case of mixed dominance simulations, we randomly assign a neutral allele’s phenotype dominance based on the mixed proportions described above.

Finally, for each parameter set and replicate, we standardize each individual’s genotypic score by dividing by the standard deviation of the distribution of genotypic scores across all sampled individuals in all three populations. By placing the genotypic scores from all sampled individuals on a common, global variation-based scale, standardization facilitates meaningful comparisons between simulations under different parameter sets.

### Runs of Homozygosity

To identify runs of homozygosity in the genotype data, we first generated tped files from VCF file produced by msprime using PLINK^77^ (v1.90b6.21) (www.cog-genomics.org/plink/1.9/). Then, we called runs of homozygosity by using GARLIC.^6, 78, 79^ Allele frequencies for GARLIC were estimated separately for each population within each replicate using the 500 sampled individuals.

Based on the output of GARLIC, we obtained the size class and location of all the ROH regions in all the sampled individuals of all the replicates. We also calculated the total length of ROH for each size class in all the sampled individuals of all the replicates. These results are used for analyzing the relationship between ROH and genotypic score. The parameters used in GARLIC are set as follows: –-map plink.chr1.GRCh37.map, –-size-bounds 0.25 1, –-centromere dummy.centromeres.txt, –-winsize 10, –-auto-winsize, –-auto-overlap-frac. By using this software, we classified the ROH in genotype data generated from each simulation into short ROH (shorter than 0.25cM), medium ROH (longer than 0.25cM but shorter than 1cM), and long ROH (longer than 1cM). We adopted cM as the unit for ROH length in this study because it directly reflects the genealogical age of the underlying haplotype by incorporating local recombination rate variation (see Figures S37–S38 for Mb-based robustness analysis).

Because GARLIC uses allele frequencies, its ROH classification may be differentially affected by the allele frequency spectrum of bottlenecked versus high-diversity populations as low-diversity populations require more statistical evidence to identify true autozygosity. Our robustness analysis using frequency-independent GERMLINE yielded consistent results (Figure S36).

### Statistical Analysis

For each replicate, we place individuals into three categories: individuals with no ROH, individuals with at least one ROH that contributes to the genotypic score, and individuals with ROH but none contribute to the genotypic score. Each of these categories are further stratified by ROH size class, and then we calculate the mean across replicates.

For all population models and genetic architectures, we calculated the genotypic score per-cM contributed by all ROH and non-ROH regions which contribute to the genotypic score. The per-cM contribution (*C*) for a specific genomic region type (*k*, where *k* ∈ *long ROH*, *medium ROH*, *s*ℎ*ort ROH*, *non* − *ROH*}) in an individual (*i*) was calculated as the ratio of the total genotypic score contributed by that region (*P_i_*_,*k*_) to the total length of region in cM (*L_i_*_,*k*_):

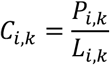

To obtain a single summary value for each replicate (*r*), we then calculated the mean of the per-cM contributions from all 500 sampled individuals of each population (*N* = 500). When calculating the mean, we only considered individuals that have ROH of the given size class and only ROH that contributed to the genotypic score. This mean value, denoted as 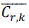, represents the result for that replicate:

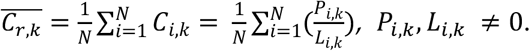

Where N is the total number of individuals satisfying the conditions above and is not necessarily 500. Moreover, we calculated the difference in per-cM contribution between different categories of ROH and non-ROH regions under various scenarios as a function of tau values. The difference in per-cM contribution (*D_r_*) between different categories of ROH and non-ROH regions is calculated as follows:

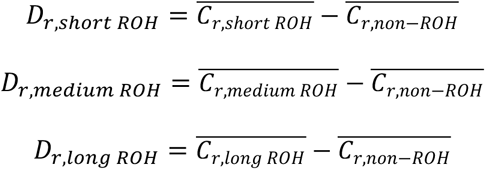

Finally, we examined the proportion (*Prop_i_*_,*k*_) of the total genotypic score attributed to different categories of ROH and non-ROH regions across these simulations. We first calculated the total genotypic score for each individual (*P_i_*_,*total*_) by following:

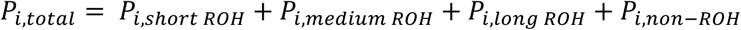

Then, the proportion of genotypic scores attributed to different ROH and non-ROH regions are calculated as follows:

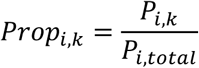

The summary value for each replicate is calculated by following:

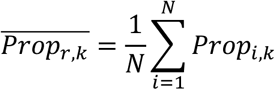

To statistically evaluate the impact of different genomic regions on genotypic scores, for each combination of population model, dominance scenario, and tau value, we applied a series of pairwise comparisons on the distributions of per-cM contributions generated from the 500 replicates. We used the Wilcoxon signed-rank test to account for the paired nature of the data within each replicate. This test was applied to compare: (i) the contribution of each ROH category (Short, Medium, and Long) against that of the non-ROH regions, and (ii) the contributions between different ROH categories themselves (e.g., Long ROH vs. Medium ROH). To correct for the multiple comparisons performed across all pairwise tests and experimental conditions, the resulting p-values were adjusted using the Holm-Bonferroni method. (See Supplemental material. Table S1 and S2.)

### Robustness analyses

#### Comparison with 1000 Genomes

To assess whether the length distribution and abundance of ROH segments generated by our simulations are biologically reasonable, we called ROH in the African (AFR), European (EUR), and East Asian (EAS) populations from the 1000 Genomes Project (phase 3).^80^ The AFR population was constructed by merging the following subpopulations: ACB, ASW, ESN, GWD, LWK, MSL, and YRI, comprising 661 individuals in total. The EAS population was constructed by merging subpopulations CDX, CHB, CHS, JPT, and KHV, comprising 504 individuals. The EUR population was constructed by merging subpopulations CEU, FIN, GBR, IBS, and TSI, comprising 503 individuals. The procedure was as follows. First, we retrieved the sample IDs of all sub-populations belonging to these three super-populations. Second, we used bcftools^81^ and PLINK (v1.90b6.21) (www.cog-genomics.org/plink/1.9/)^77^ to filter for autosomal biallelic loci and extracted genome-wide autosomal genotypes for each sub-population. Third, we called ROH on each sub-population using GARLIC^78^ and then pooled the resulting ROH lengths by super-population. Finally, we computed *F_R_*_0*H*_, the number of ROH per individual (nROH), and the ROH length distribution for each super-population. The parameters used for ROH calling were: –-cm –-map chraut.map –-build hg19 –-size-bounds 0.25 1 –-winsize 10 –-auto-winsize –-auto-overlap-frac –-error 0.0001 –-lod-cutoff 0 –-tped-missing 9. The ROH size-class thresholds were the same as those used elsewhere in this study.

#### Genetic architecture diagnostics

To assess whether our simulations are biologically reasonable, we conducted the following diagnostics.

#### Sparse model

To assess the robustness of our conclusions, we introduced a sparse model. In the sparse model, for each replicate, we randomly sampled 30% of the deleterious mutations simulated in SLiM as causal variants. In contrast, in the non-sparse model, all simulated deleterious mutations were considered causal. All other procedures were identical to those described above.

For the sparse model, we computed the proportion of individuals, per-cM genotypic score contribution (under both conditional and unconditional scenarios), the proportion of genotypic score explained by ROH, and the difference in per-cM contribution between ROH and non-ROH regions as a function of τ. Under the conditional scenario, we considered only individuals with ROH contributing non-zero genotypic scores; under the unconditional scenario, we considered all individuals. For the sparse model, we performed the same statistical significance tests for both the conditional and unconditional per-cM genotypic score contributions. Detailed results are presented in Supplementary Tables S8–S11.

#### Allele frequency spectrum and effect frequency spectrum

To further validate the simulation setup, we computed the allele frequency spectrum and the effect frequency spectrum for all populations across all genetic architectures. Specifically, the effect frequency spectrum refers to the distribution of effect sizes assigned to individual causal variants under each regime. We considered both the sparse and non-sparse models.

#### Diagnostic of causal alleles

We computed the average number of causal variants per individual, the average number of homozygous causal variants per individual, the proportion of causal variants located in ROH regions, and the proportion of homozygous causal variants located in ROH regions, across all populations and genetic architectures. Both the sparse and non-sparse models were considered.

#### Genetic variance explained by rare versus common alleles

To assess whether the range of τ values we selected (0.7, 0.8, 0.9, 1.0, 1.1, 1.2, 1.3) is reasonable, we computed the proportions of genetic variance explained by rare versus common variants under each τ value within each regime. The calculation proceeded as follows. Each individual’s total genotypic score is the sum of effect sizes across all causal variant sites. We partitioned each individual’s genotypic score into three components according to the allele frequency of each contributing causal variant in the population: variants with frequencies below 0.1%, variants with frequencies between 0.1% and 1%, and variants with frequencies above 1%. For each population, we then computed the genetic variance across the 500 sampled individuals for each of these three components, and calculated the proportion of total genetic variance attributable to each. Both the sparse and non-sparse models were considered.

#### Genetic variance explained by ROH regions

To further explore the relationship between ROH and genetic variance, we computed the proportion of genetic variance explained by ROH regions under each regime. The procedure was similar to that described above. We partitioned each individual’s genotypic score into the contribution from causal variants located within ROH regions and the contribution from causal variants located in non-ROH regions. For each population, we then computed the genetic variance across the 500 sampled individuals for each of these two components, and calculated the proportion of total genetic variance attributable to each. Both the sparse and non-sparse models were considered.

#### Additive and dominance variance

We also computed the proportions of additive variance and dominance variance within the total genetic variance under each regime. For this calculation, we followed the definitions of breeding value and dominance value as in Lynch et al.^82^ Based on these definitions, we performed Fisher variance decomposition at each causal locus across the 500 sampled individuals per population in each replicate. Under this decomposition, each individual’s genotypic score can be partitioned into a breeding value and a dominance value. For each population, we then computed the genetic variance across the 500 sampled individuals for each of these two components and calculated the proportion of total genetic variance attributable to each. Both the sparse and non-sparse models were considered.

#### GARLIC versus GERMLINE

To assess the robustness of the results obtained from the ROH caller GARLIC,^78^ we used another fully frequency-independent ROH caller, GERMLINE,^83^ to call ROH independently. Specifically, we used GERMLINE to obtain ROH segments in units of bp and then interpolated the segment lengths using the genetic distance map of the first 100 Mb of human chromosome 1 to obtain ROH segments in units of cM. We then classified these segments using the same ROH size-class thresholds applied elsewhere in this study. Finally, we used the ROH segments obtained from GERMLINE to recompute the conditional per-cM genotypic score under the fully recessive scenario. The GERMLINE parameters used were: –homozonly-min_m 0.0000001 –bits 128. The minimum segment length was set so low that the resulting ROH segments were not filtered in practice. All other parameters were set to the GERMLINE manual defaults.

#### cM versus Mb

To examine the potential effect of unit choice on the robustness of our conclusions, we computed the Mb lengths of the ROH segments obtained from GARLIC.^78^ We then reclassified the ROH segments using the following criteria: segments longer than 1 Mb were classified as long ROH, segments shorter than 0.25 Mb as short ROH, and segments in between as medium ROH. We computed the conditional per-Mb genotypic score under the fully recessive scenario. In addition, under the current cM-based ROH classification, we computed the length distributions of the three ROH classes under the fully recessive scenario for all three populations. These distributions were calculated and presented in both cM and Mb (Figure S38).

## Acknowledgement

This work was supported by the National Institute of General Medical Sciences of the National Institutes of Health under Award Number R35GM146926 (ZAS and MP). This work was also supported by Eberly College of Science Startup Fund (ZAS and MP). Computations for this research were performed using the Pennsylvania State University’s Institute for Computational Data Sciences’ Roar supercomputer.

## Code and data availability

All simulation scripts and analytical pipelines described in this study are available at the GitHub repository (https://github.com/Mingzuyu-Pan/ROH_phenotype_simulation).

