## Supplemental Data for "The Influence of Demographic History and Genetic Architecture on Complex Traits via Runs of Homozygosity"

### Supplemental Material

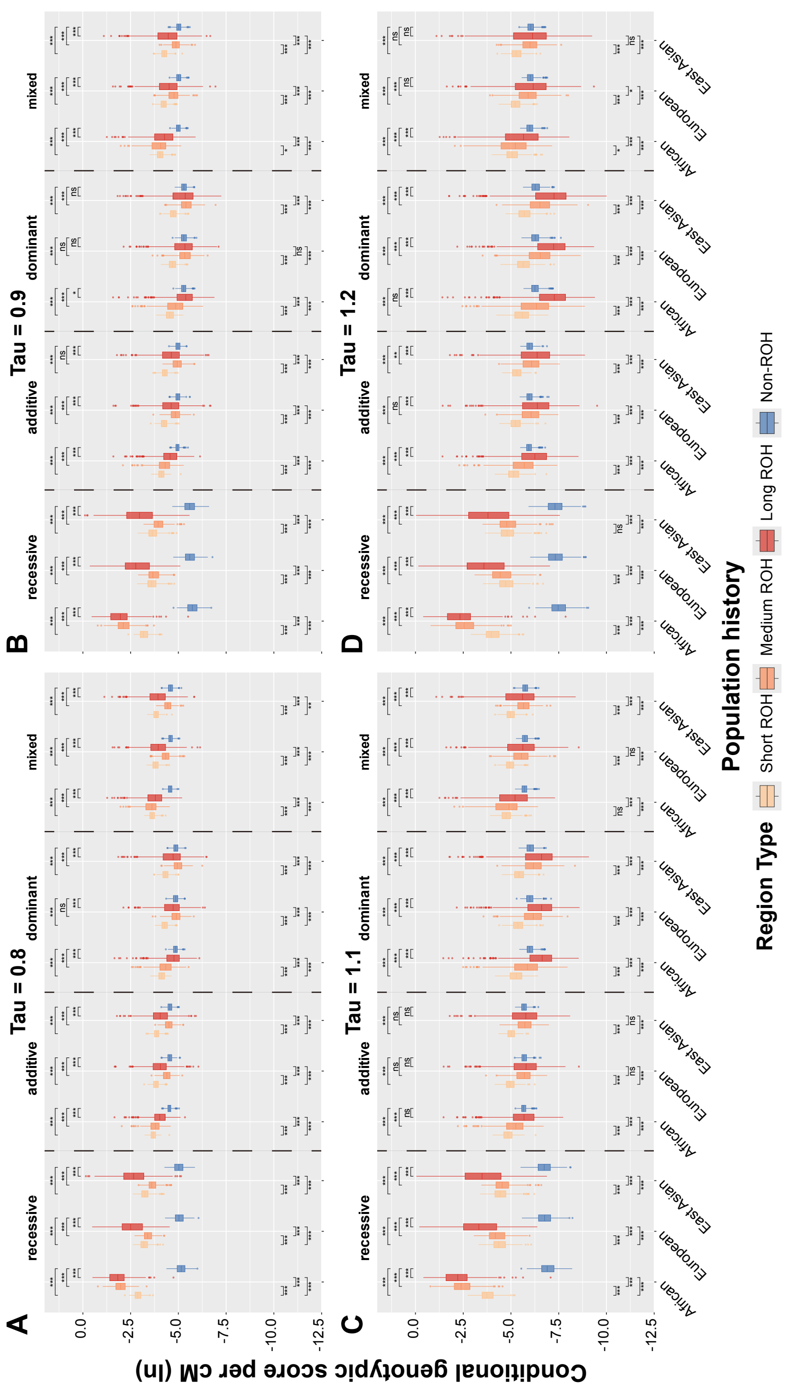


Figure S1. Conditional per-cM genotypic score contribution (log-transformed; deleterious mutations) for various regions, population histories, and genetic architectures. Regions are: Short ROH (< 0.25 cM); Medium ROH (0.25-1 cM); Long ROH (> 1 cM); Non-ROH. Causal genotypes are generated from deleterious mutations. The parameter tau is varied for different weighting of rare alleles in phenotype score calculation. (A) $\tau = 0.8$, (B) $\tau = 0.9$, (C) $\tau = 1.1$, (D) $\tau = 1.2$. Significance levels for adjusted P-values: *** P_adj < 0.001, ** P_adj < 0.01, * P_adj < 0.05, ns P_adj ≥ 0.05.

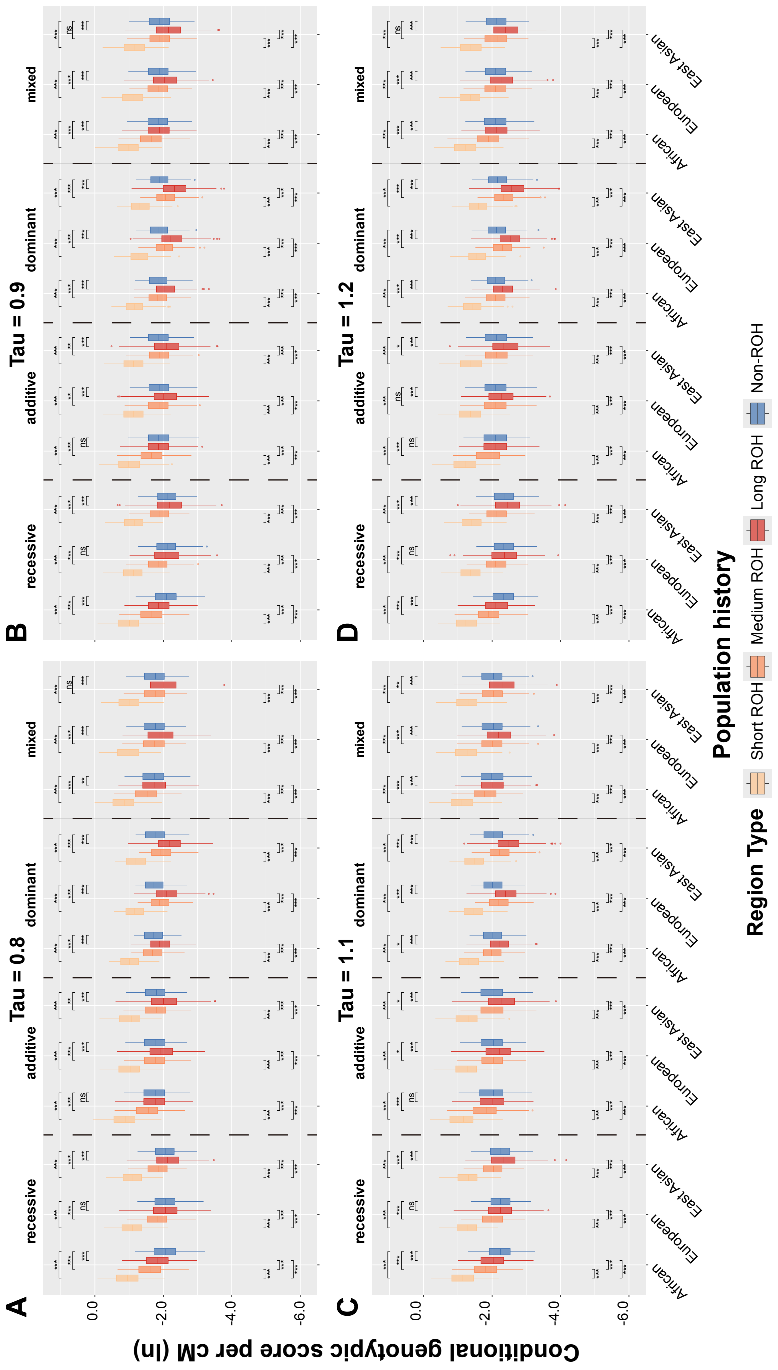


Figure S2. Conditional per-cM genotypic score contribution (log-transformed; neutral mutations) for various regions, population histories, and genetic architectures. Regions are: Short ROH (< 0.25 cM); Medium ROH (0.25-1 cM); Long ROH (> 1 cM); Non-ROH. Causal genotypes are generated from neutral mutations. The parameter tau is varied for different weighting of rare alleles in phenotype score calculation. (A) $\tau = 0.8$, (B) $\tau = 0.9$, (C) $\tau = 1.1$, (D) $\tau = 1.2$. Significance levels for adjusted P-values: *** P_adj < 0.001, ** P_adj < 0.01, * P_adj < 0.05, ns P_adj ≥ 0.05.

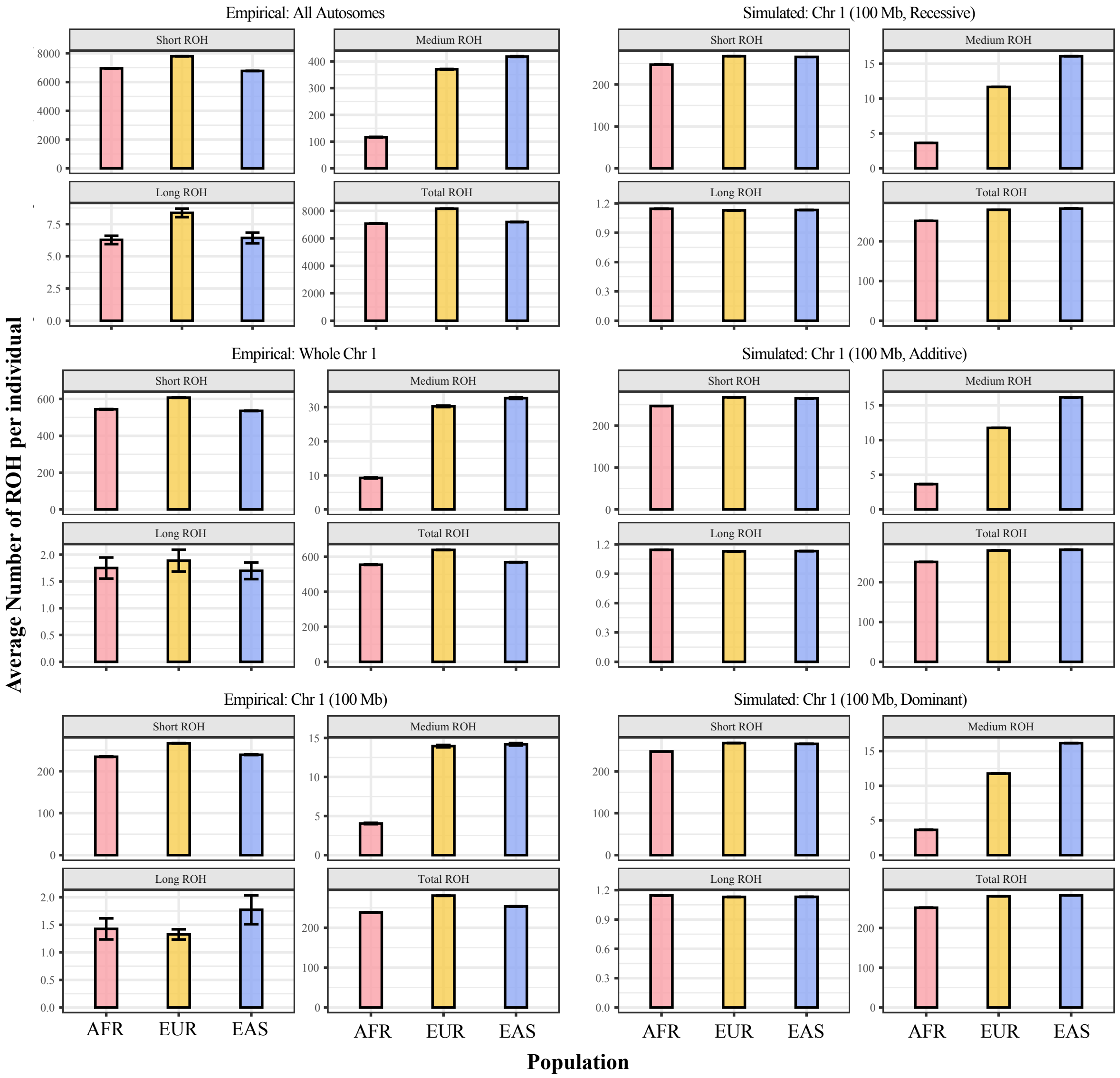


Figure S3. Average number of ROH per individual by ROH class across empirical and simulated scenarios. Bars show the average number of short (< 0.25 cM), medium (0.25–1 cM), long (> 1 cM), and total ROH per individual for African (AFR), European (EUR), and East Asian (EAS) populations, under three empirical scenarios from the 1000 Genomes Project (ROH called across all autosomes, full chromosome 1, and the first 100 Mb of chromosome 1) and three simulated dominance models (fully recessive, fully additive, fully dominant).

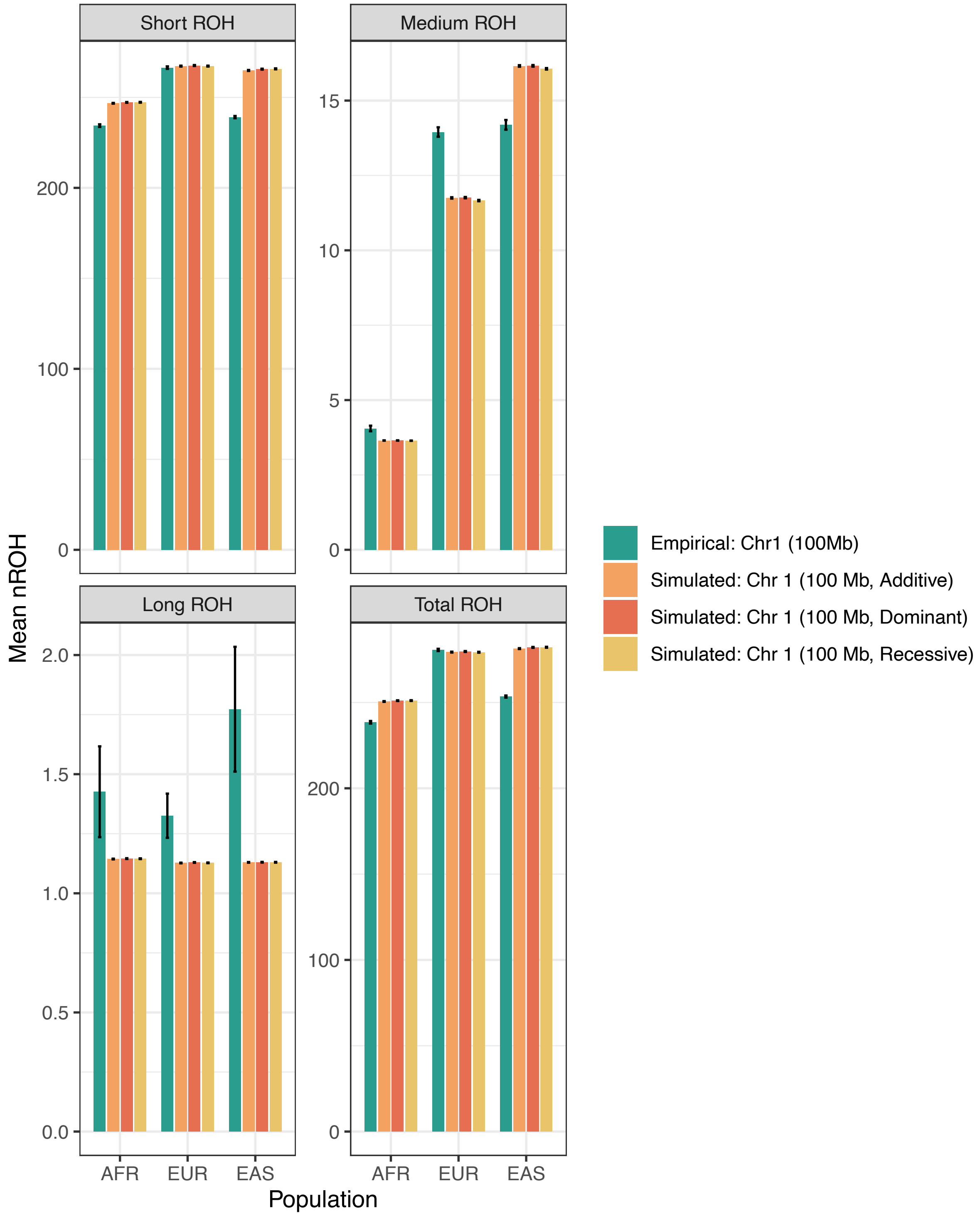


Figure S4. Average number of ROH per individual by ROH class, comparing the empirical chr 1 (100 Mb) scenario with simulated 100 Mb scenarios. Bars show the average number of short (< 0.25 cM), medium (0.25–1 cM), long (> 1 cM), and total ROH per individual for African (AFR), European (EUR), and East Asian (EAS) populations, under the empirical Chr 1 (100 Mb) scenario from the 1000 Genomes Project and three simulated dominance models (fully recessive, fully additive, fully dominant).

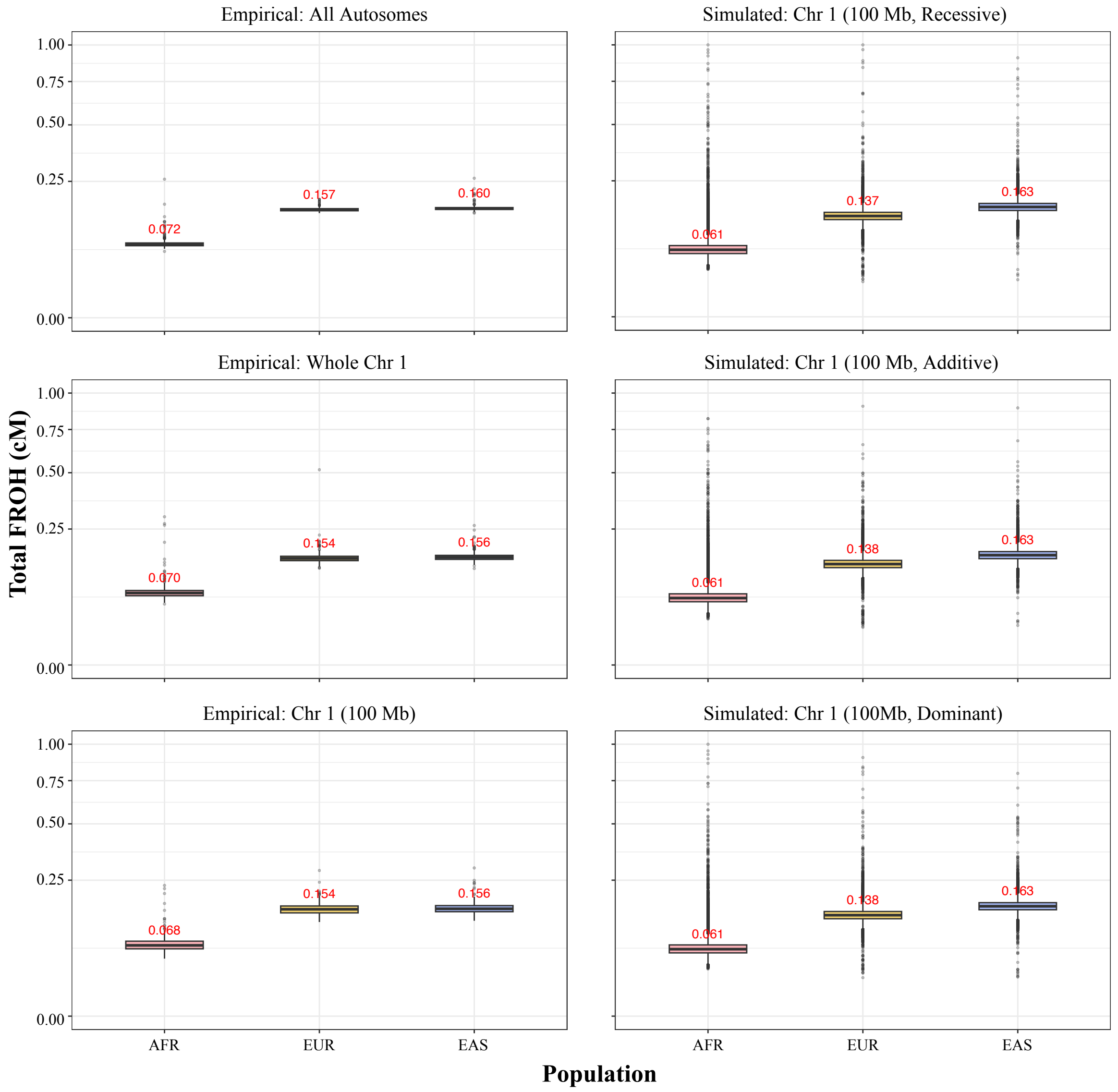


Figure S5. $F_{ROH}$ per individual across empirical and simulated scenarios. Box plots show per-individual $F_{ROH}$ (the proportion of the analyzed genomic region covered by ROH) for African (AFR), European (EUR), and East Asian (EAS) populations, under three empirical scenarios from the 1000 Genomes Project (ROH called across all autosomes, full chromosome 1, and the first 100 Mb of chromosome 1) and three simulated dominance models (fully recessive, fully additive, fully dominant). Red numerals indicate the median $F_{ROH}$ for each population–scenario combination.

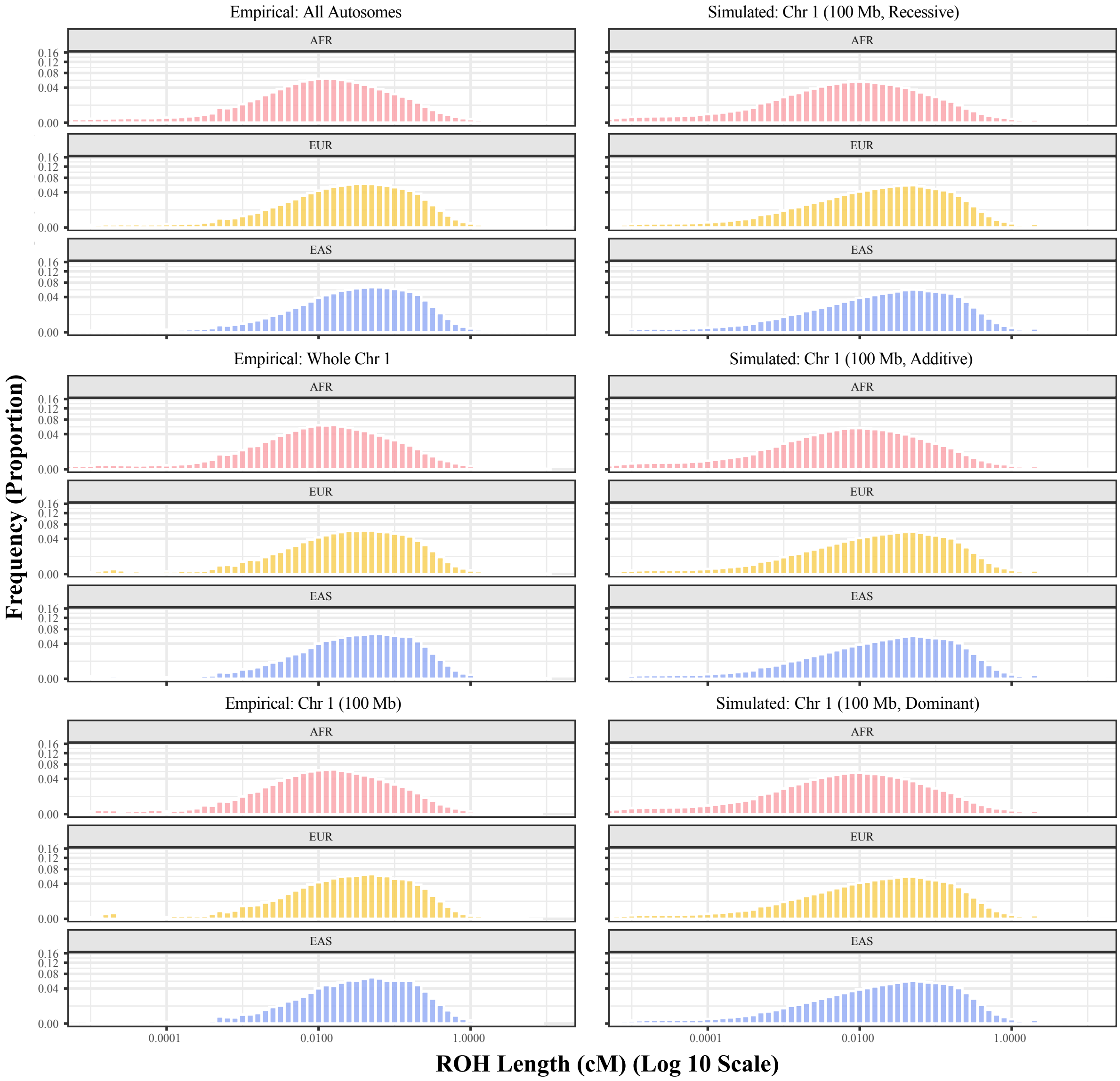


Figure S6. ROH length distribution (cM) across empirical and simulated scenarios. Histograms show the distribution of ROH segment lengths (in cM, log-scaled x-axis; square-root-scaled y-axis) for African (AFR), European (EUR), and East Asian (EAS) populations, under three empirical scenarios from the 1000 Genomes Project (ROH called across all autosomes, full chromosome 1, and the first 100 Mb of chromosome 1) and three simulated dominance models (fully recessive, fully additive, fully dominant). For simulated data, ROH segments were pooled across replicates within each population.

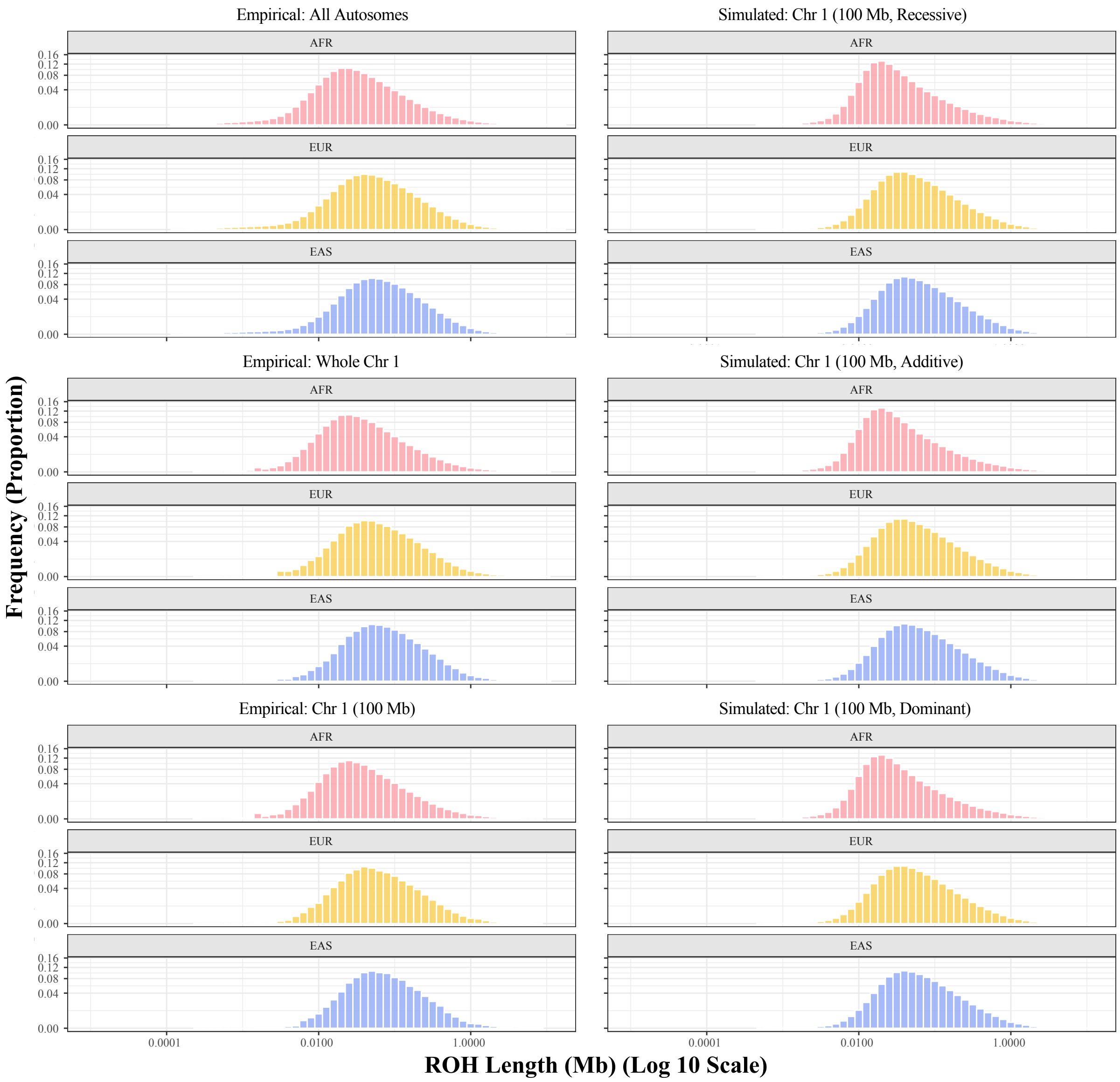


Figure S7. ROH length distribution (Mb) across empirical and simulated scenarios. Histograms show the distribution of ROH segment lengths (in cM, log-scaled x-axis; square-root-scaled y-axis) for African (AFR), European (EUR), and East Asian (EAS) populations, under three empirical scenarios from the 1000 Genomes Project (ROH called across all autosomes, full chromosome 1, and the first 100 Mb of chromosome 1) and three simulated dominance models (fully recessive, fully additive, fully dominant). For simulated data, ROH segments were pooled across replicates within each population.

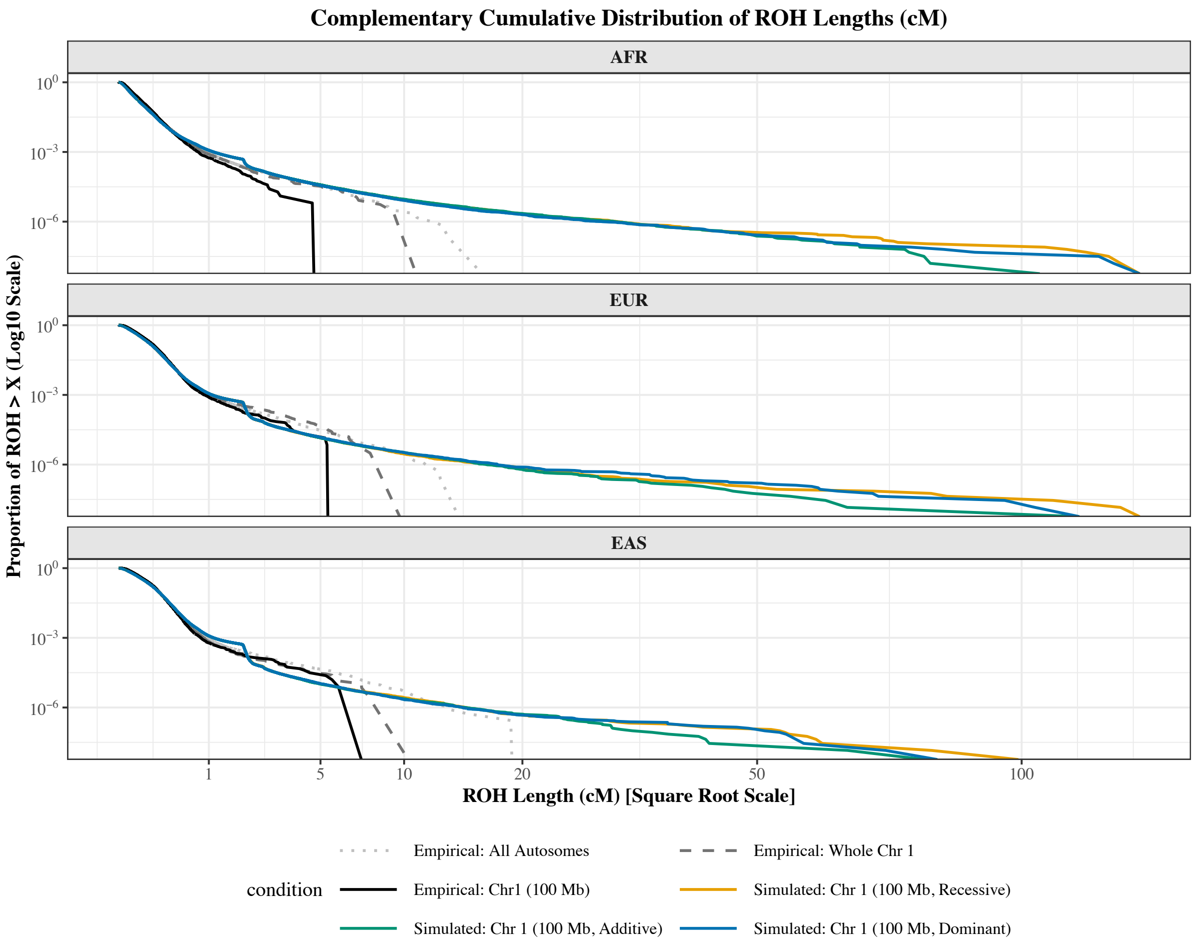


Figure S8. Complementary cumulative distribution function (CCDF) of ROH lengths in cM. CCDF of ROH segment lengths in cM (x-axis on square-root scale; y-axis on log scale to highlight the long-length tail) for African (AFR), European (EUR), and East Asian (EAS) populations, under three empirical scenarios from the 1000 Genomes Project (ROH called across all autosomes, full chromosome 1, and the first 100 Mb of chromosome 1; gray dotted, dashed, and solid lines, respectively) and three simulated dominance models (fully recessive, fully additive, fully dominant; colored solid lines). Each empirical curve truncates at the maximum ROH length observable under its corresponding scenario. The three simulated dominance models produce nearly identical CCDFs, indicating that the long-tail behavior is not affected by the choice of dominance model.

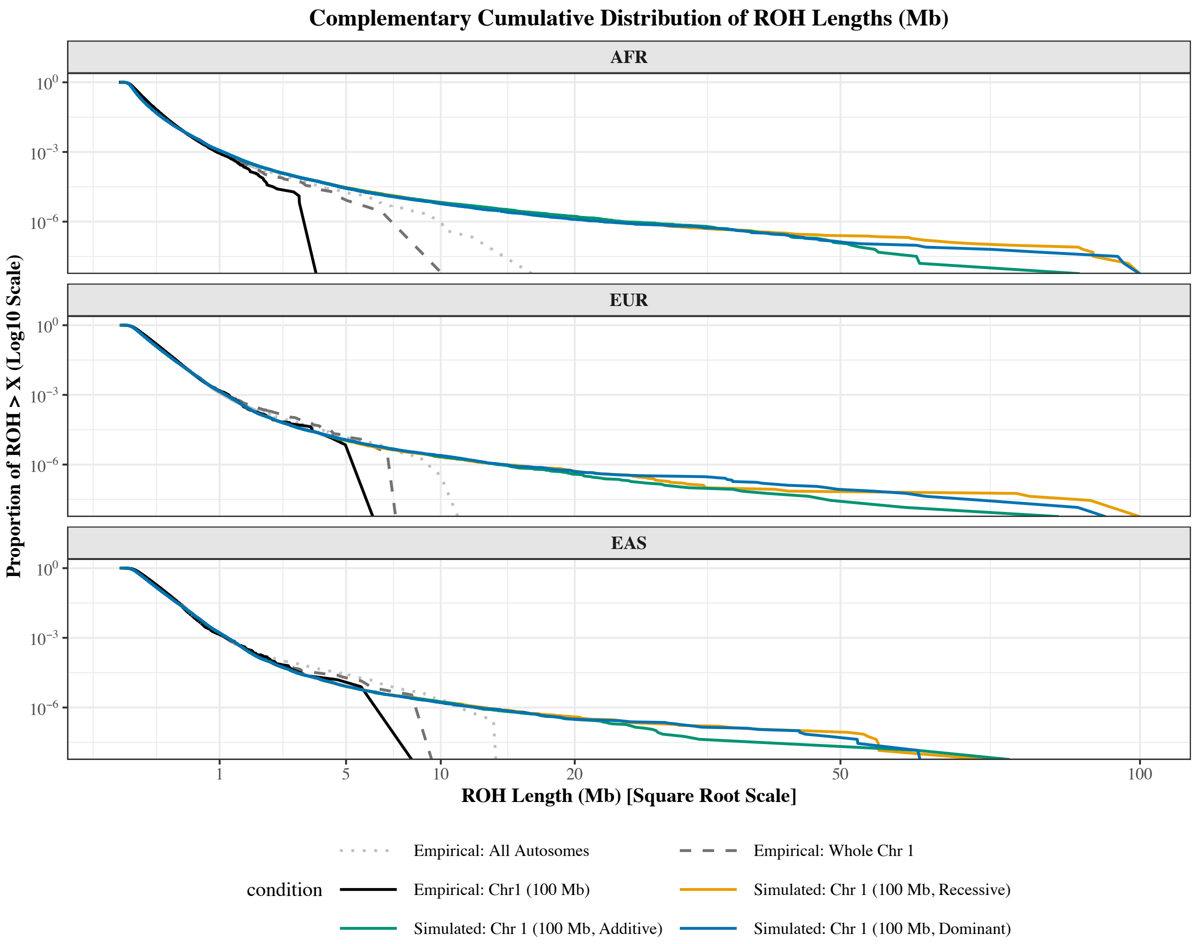


Figure S9. Complementary cumulative distribution function (CCDF) of ROH lengths in Mb. CCDF of ROH segment lengths in Mb (x-axis on square-root scale; y-axis on log scale to highlight the long-length tail) for African (AFR), European (EUR), and East Asian (EAS) populations, under three empirical scenarios from the 1000 Genomes Project (ROH called across all autosomes, full chromosome 1, and the first 100 Mb of chromosome 1; gray dotted, dashed, and solid lines, respectively) and three simulated dominance models (fully recessive, fully additive, fully dominant; colored solid lines). Each empirical curve truncates at the maximum ROH length observable under its corresponding scenario. The three simulated dominance models produce nearly identical CCDFs, indicating that the long-tail behavior is not affected by the choice of dominance model.

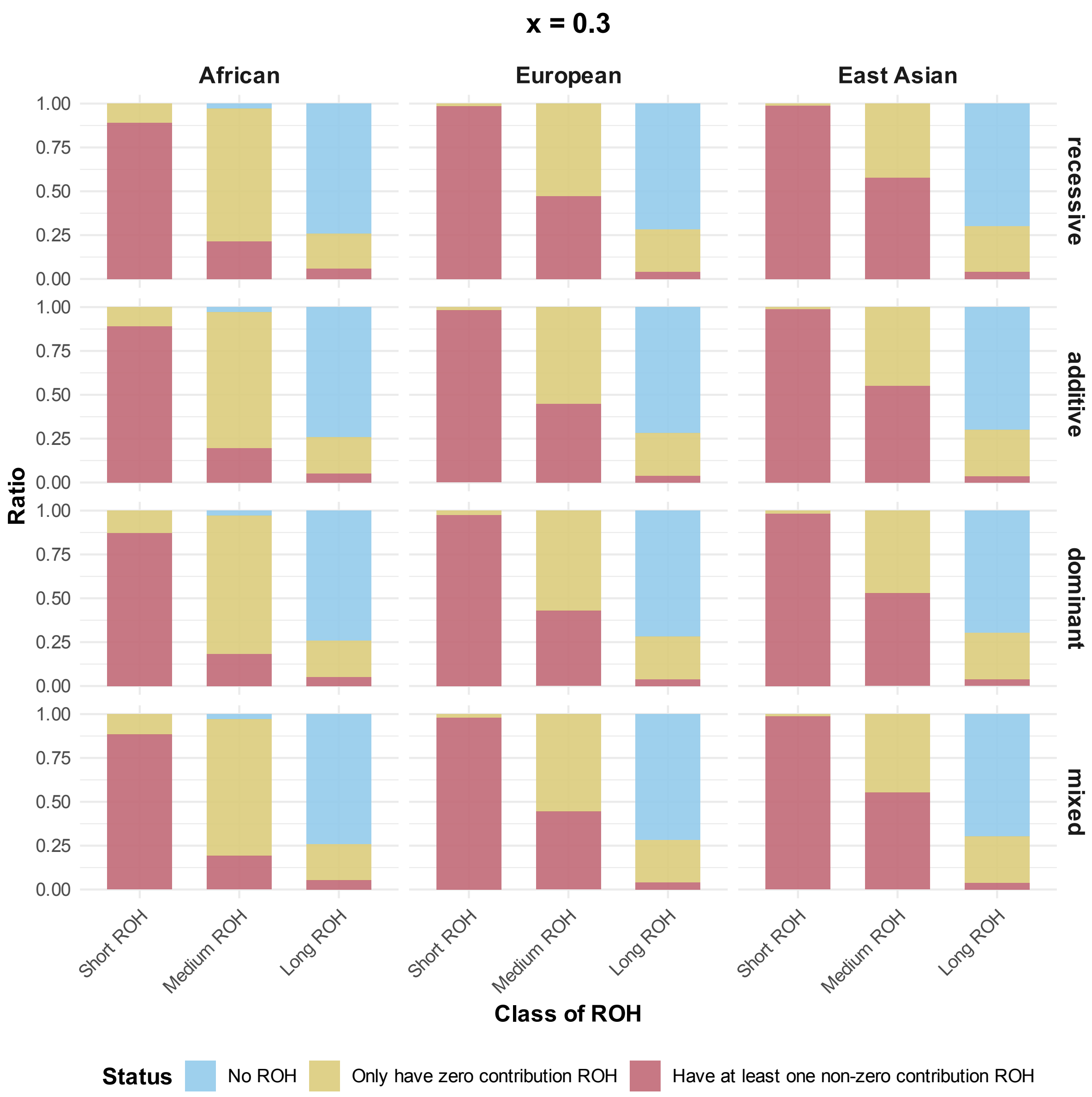


Figure S10. (Sparse model) Proportion of individuals with ROH contributing to genotypic score across various population histories and genetic architectures (x = 0.3, the fraction of simulated deleterious mutations treated as causal). Genotypic scores are only contributed by deleterious mutations. Blue bars represent the individuals lacking ROH (thus no genotypic score contribution); red bars represent the individuals having at least one ROH with a genotypic score contribution; yellow bars represent the individuals having ROH with no genotypic score contribution. Region definitions: Short ROH (< 0.25 cM); Medium ROH (0.25-1 cM); Long ROH (> 1 cM).

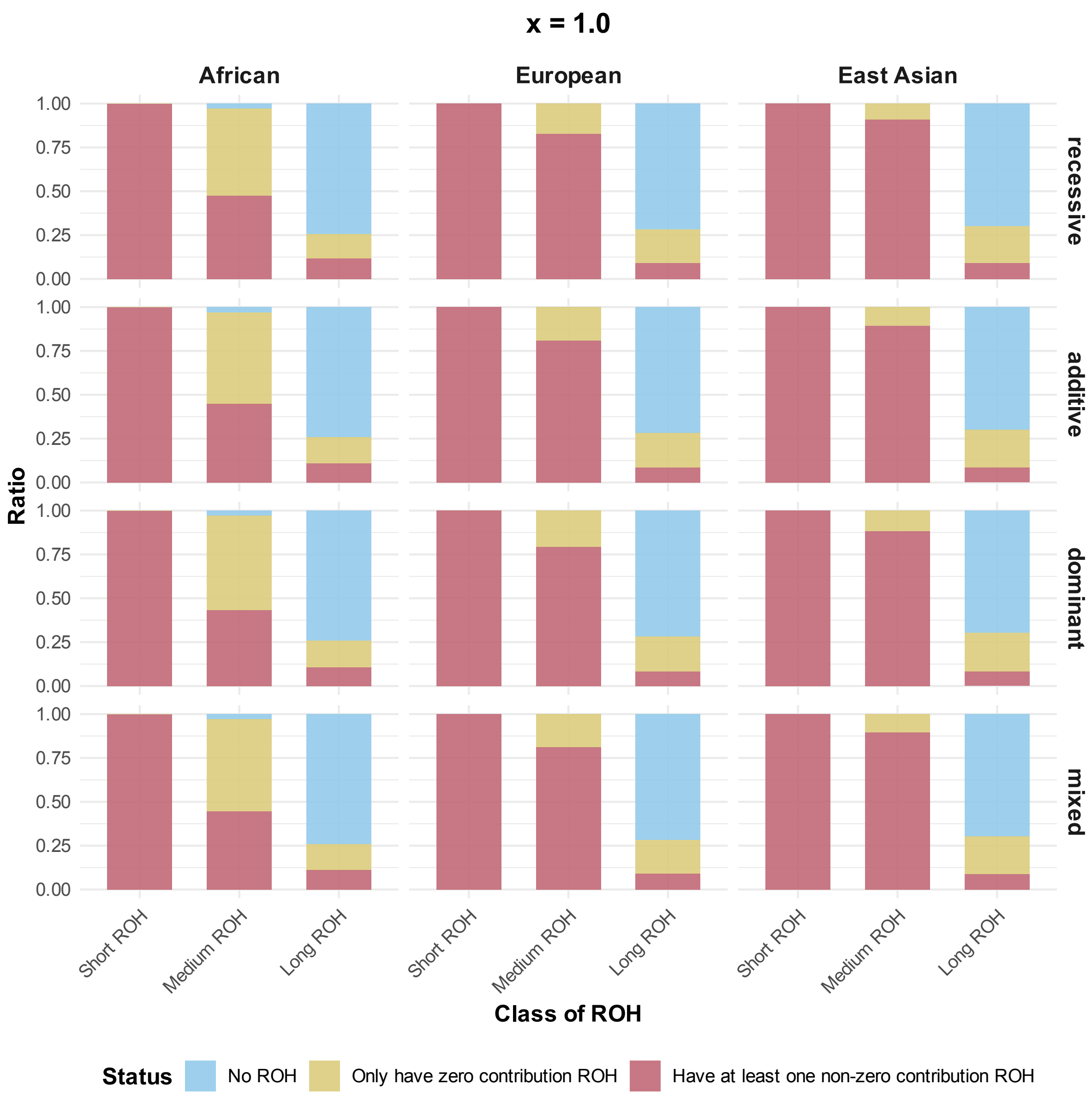


Figure S11. (Non-sparse model) Proportion of individuals with ROH contributing to genotypic score across various population histories and genetic architectures (x = 1.0, the fraction of simulated deleterious mutations treated as causal). Genotypic scores are only contributed by deleterious mutations. Blue bars represent the individuals lacking ROH (thus no genotypic score contribution); red bars represent the individuals having at least one ROH with a genotypic score contribution; yellow bars represent the individuals having ROH with no genotypic score contribution. Region definitions: Short ROH (< 0.25 cM); Medium ROH (0.25-1 cM); Long ROH (> 1 cM).

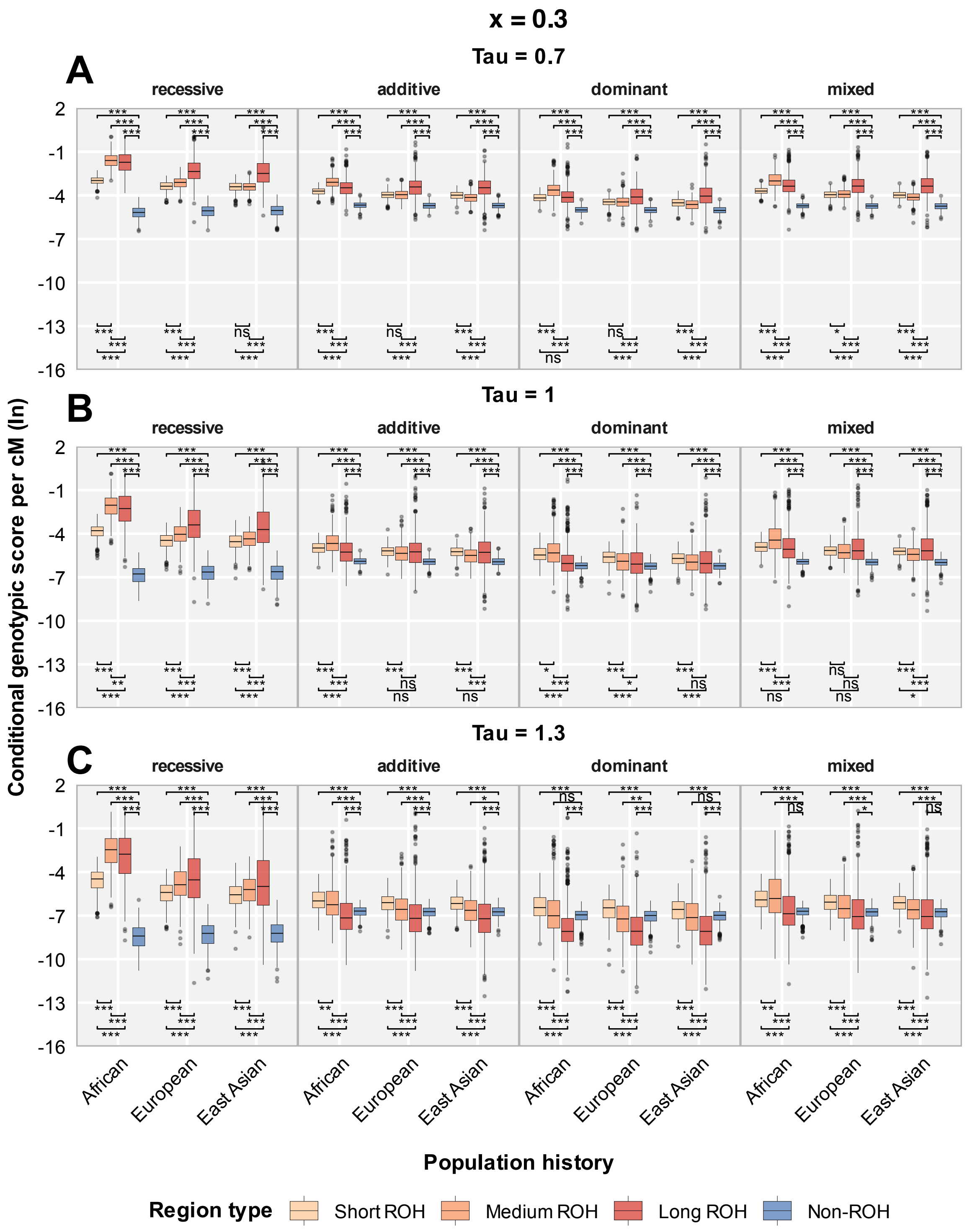


Figure S12. (Sparse model) Conditional per-cM genotypic score contribution (log-transformed; deleterious mutations) for various regions, population histories, and genetic architectures (x = 0.3, the fraction of simulated deleterious mutations treated as causal). Regions are: Short ROH (< 0.25 cM); Medium ROH (0.25-1 cM); Long ROH (> 1 cM); Non-ROH. Causal genotypes are generated from deleterious mutations. The parameter tau is varied for different weighting of rare alleles in phenotype score calculation. (A) τ = 0.7, (B) τ = 1.0, (C) τ = 1.3. Significance levels for adjusted P-values: *** P_adj < 0.001, ** P_adj < 0.01, * P_adj < 0.05, ns P_adj ≥ 0.05.

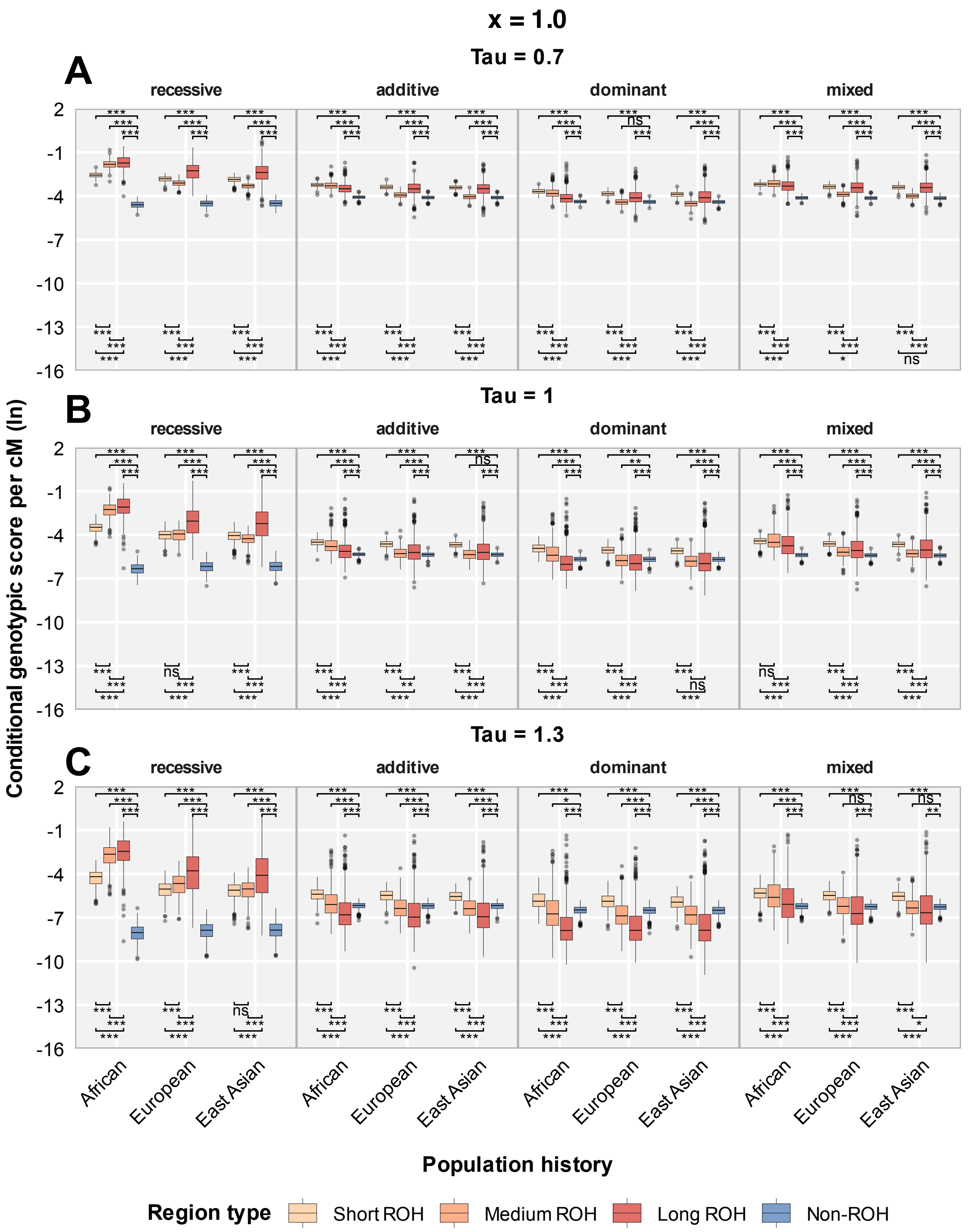


Figure S13. (Non-sparse model) Conditional per-cM genotypic score contribution (log-transformed; deleterious mutations) for various regions, population histories, and genetic architectures (x = 1.0, the fraction of simulated deleterious mutations treated as causal). Regions are: Short ROH (< 0.25 cM); Medium ROH (0.25-1 cM); Long ROH (> 1 cM); Non-ROH. Causal genotypes are generated from deleterious mutations. The parameter tau is varied for different weighting of rare alleles in phenotype score calculation. (A) $\tau = 0.7$, (B) $\tau = 1.0$, (C) $\tau = 1.3$. Significance levels for adjusted P-values: *** P_adj < 0.001, ** P_adj < 0.01, * P_adj < 0.05, ns P_adj ≥ 0.05.

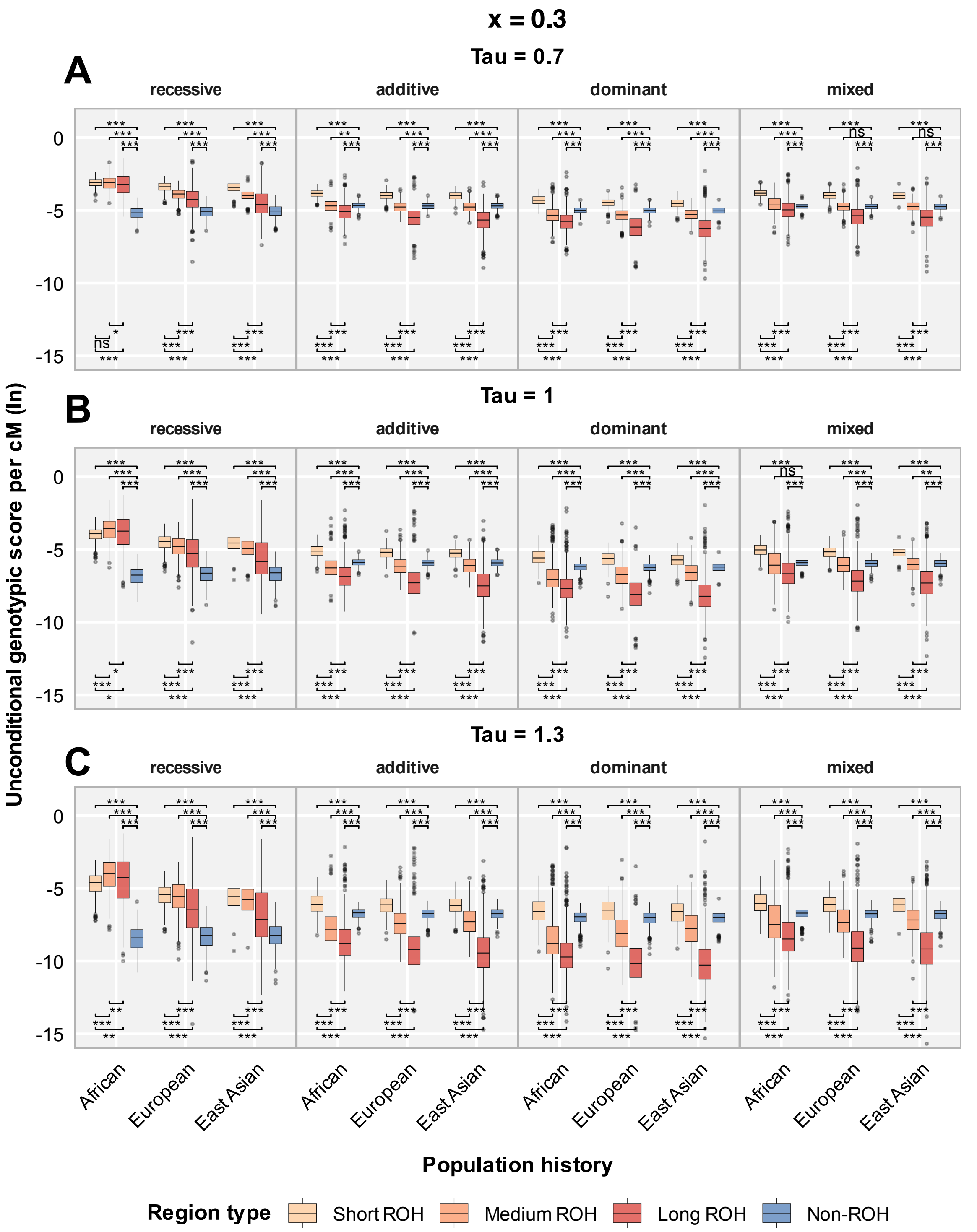


Figure S14. (Sparse model) Unconditional per-cM genotypic score contribution (log-transformed; deleterious mutations) for various regions, population histories, and genetic architectures (x = 0.3, the fraction of simulated deleterious mutations treated as causal). Regions are: Short ROH (< 0.25 cM); Medium ROH (0.25-1 cM); Long ROH (> 1 cM); Non-ROH. Causal genotypes are generated from deleterious mutations. The parameter tau is varied for different weighting of rare alleles in phenotype score calculation. (A) τ = 0.7, (B) τ = 1.0, (C) τ = 1.3. Significance levels for adjusted P-values: *** P_adj < 0.001, ** P_adj < 0.01, * P_adj < 0.05, ns P_adj ≥ 0.05.

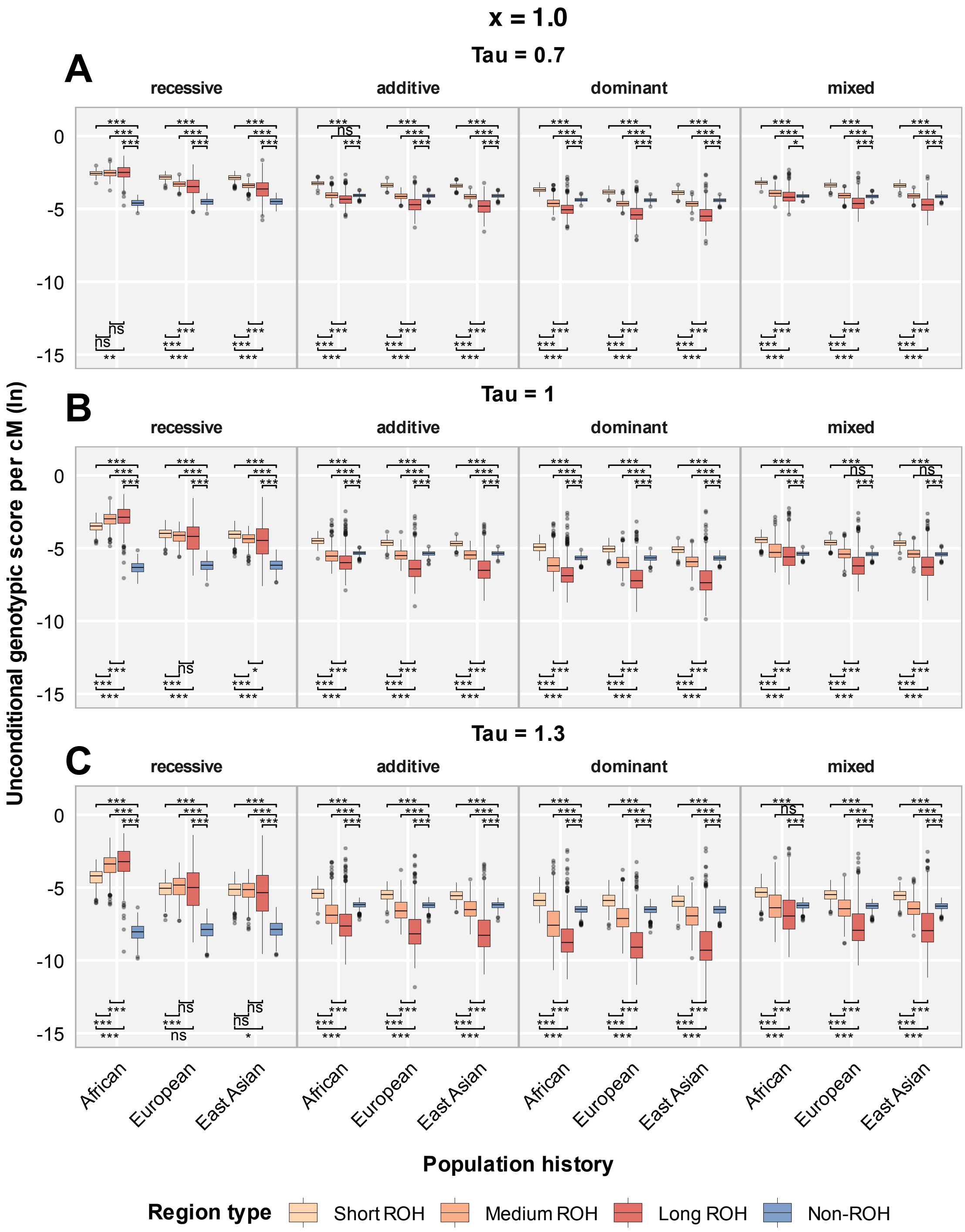


Figure S15. (Non-sparse model) Unconditional per-cM genotypic score contribution (log-transformed; deleterious mutations) for various regions, population histories, and genetic architectures (x = 1.0, the fraction of simulated deleterious mutations treated as causal). Regions are: Short ROH (< 0.25 cM); Medium ROH (0.25-1 cM); Long ROH (> 1 cM); Non-ROH. Causal genotypes are generated from deleterious mutations. The parameter tau is varied for different weighting of rare alleles in phenotype score calculation. (A) $\tau = 0.7$, (B) $\tau = 1.0$, (C) $\tau = 1.3$. Significance levels for adjusted P-values: *** P_adj < 0.001, ** P_adj < 0.01, * P_adj < 0.05, ns P_adj ≥ 0.05.

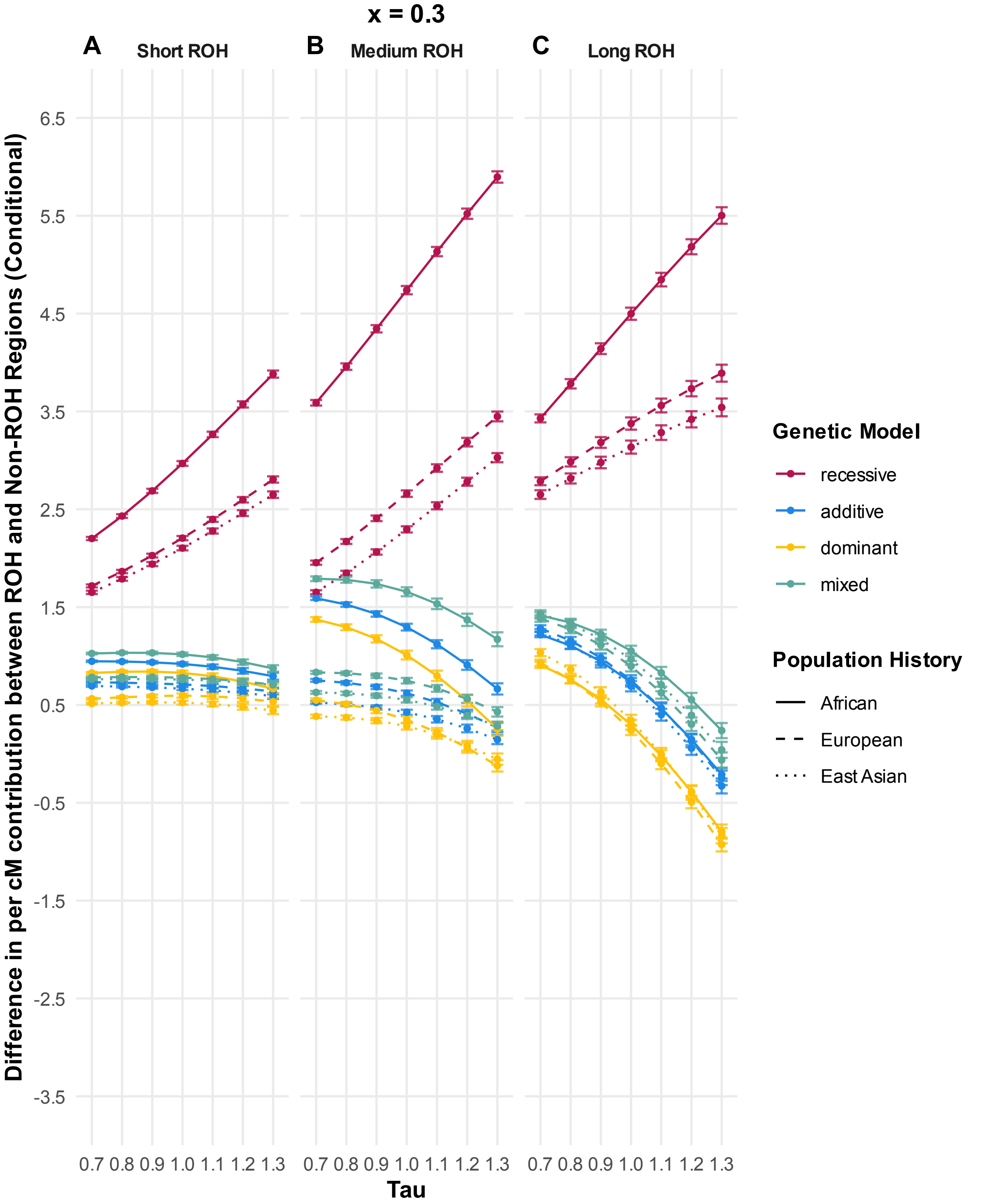


Figure S16. (Sparse model) Difference (in log space) in conditional per-cM genotypic score contribution between ROH and non-ROH regions with varied tau, various population histories and genetic architectures (x = 0.3, the fraction of simulated deleterious mutations treated as causal). Causal genotypic scores are only contributed by deleterious mutations. Panels show the difference in contribution between the specific ROH regions and non-ROH regions: (A) Short ROH regions (< 0.25 cM); (B) Medium ROH regions (0.25-1 cM); (C) Long ROH regions (> 1 cM).

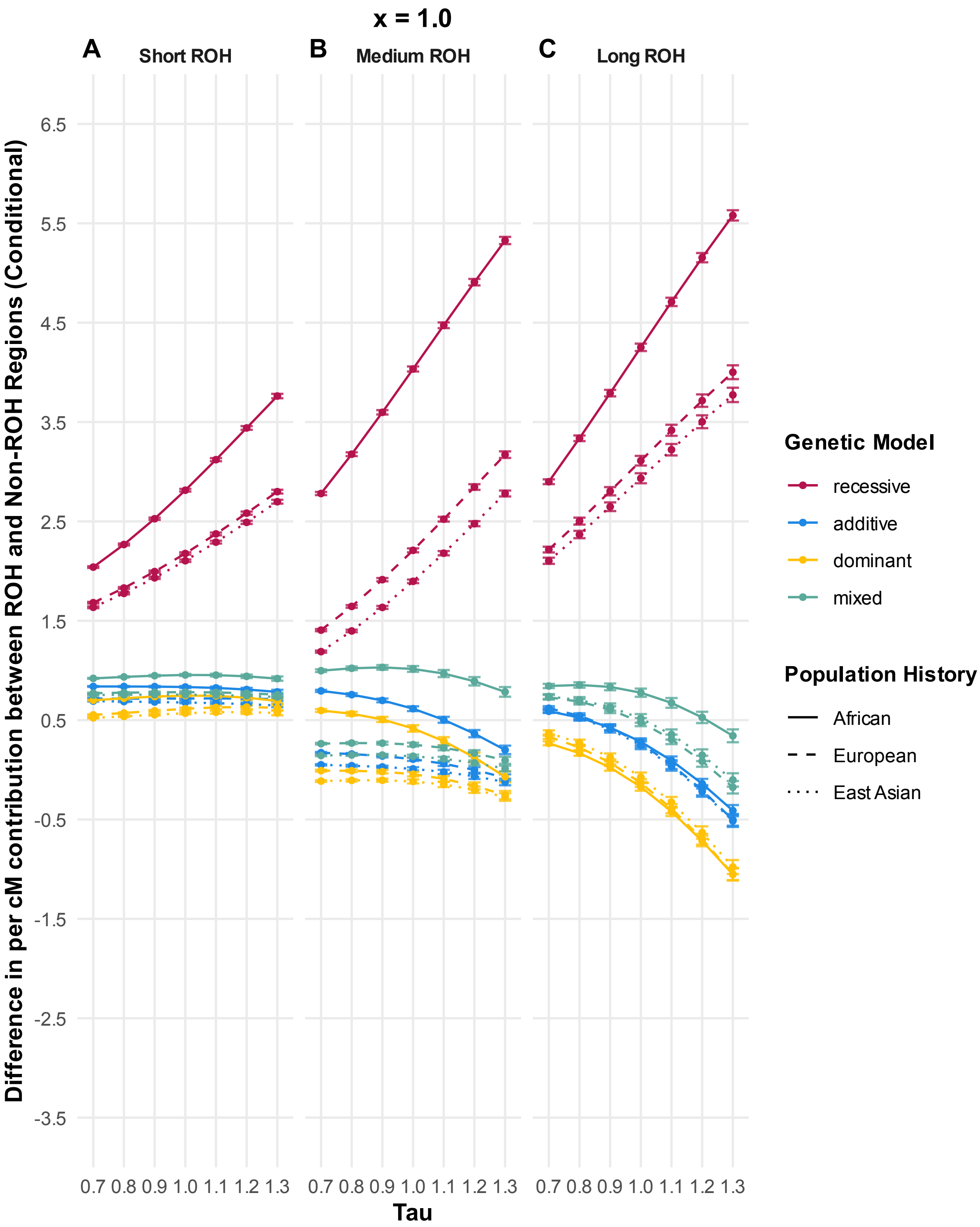


Figure S17. (Non-sparse model) Difference (in log space) in conditional per-cM genotypic score contribution between ROH and non-ROH regions with varied tau, various population histories and genetic architectures (x = 1.0, the fraction of simulated deleterious mutations treated as causal). Causal genotypic scores are only contributed by deleterious mutations. Panels show the difference in contribution between the specific ROH regions and non-ROH regions: (A) Short ROH regions (< 0.25 cM); (B) Medium ROH regions (0.25-1 cM); (C) Long ROH regions (> 1 cM).

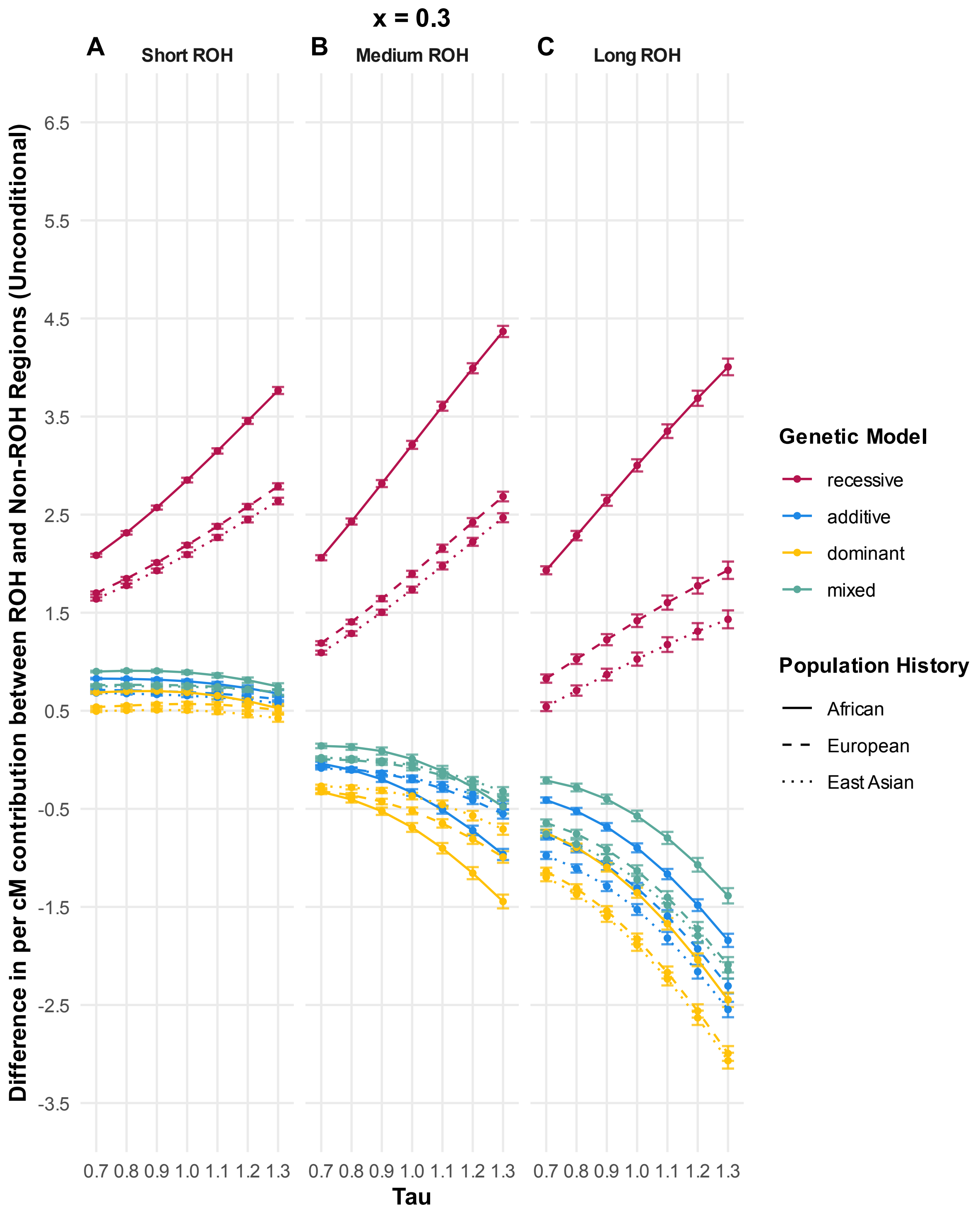


Figure S18. (Sparse model) Difference (in log space) in unconditional per-cM genotypic score contribution between ROH and non-ROH regions with varied tau, various population histories and genetic architectures (x = 0.3, the fraction of simulated deleterious mutations treated as causal). Causal genotypic scores are only contributed by deleterious mutations. Panels show the difference in contribution between the specific ROH regions and non-ROH regions: (A) Short ROH regions (< 0.25 cM); (B) Medium ROH regions (0.25-1 cM); (C) Long ROH regions (> 1 cM).

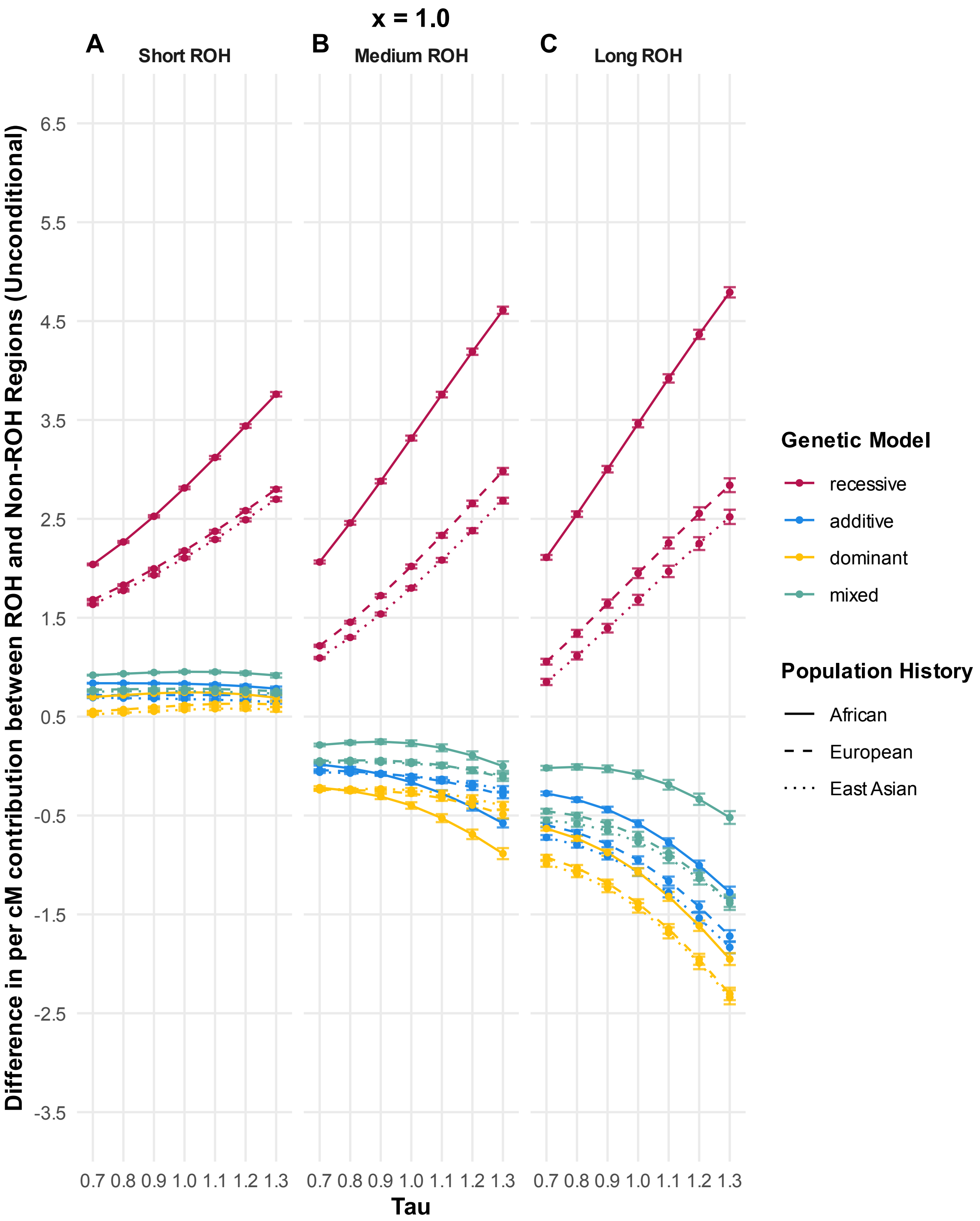


Figure S19. (Non-sparse model) Difference (in log space) in unconditional per-cM genotypic score contribution between ROH and non-ROH regions with varied tau, various population histories and genetic architectures (x = 1.0, the fraction of simulated deleterious mutations treated as causal). Causal genotypic scores are only contributed by deleterious mutations. Panels show the difference in contribution between the specific ROH regions and non-ROH regions: (A) Short ROH regions (< 0.25 cM); (B) Medium ROH regions (0.25-1 cM); (C) Long ROH regions (> 1 cM).

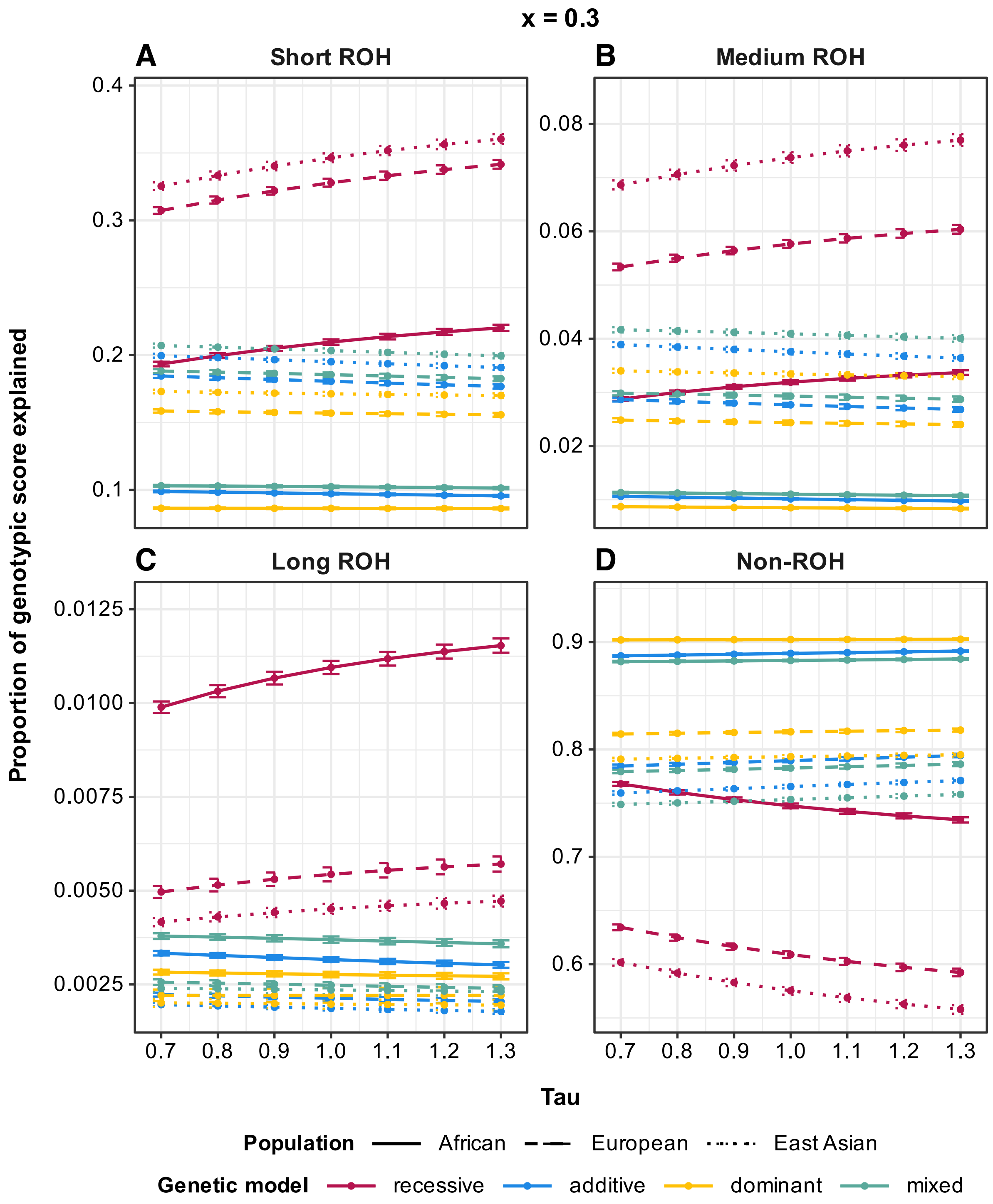


Figure S20. (Sparse model) Proportion of total genotypic score per individual explained by different types of regions with varied tau, various population histories and genetic architectures (x = 0.3, the fraction of simulated deleterious mutations treated as causal). Causal genotypic scores are only contributed by deleterious mutations. Panels show the proportion of genotypic score explained by (A) Short ROH regions (< 0.25 cM); (B) Medium ROH regions (0.25-1 cM); (C) Long ROH regions (> 1 cM); (D) Non-ROH.

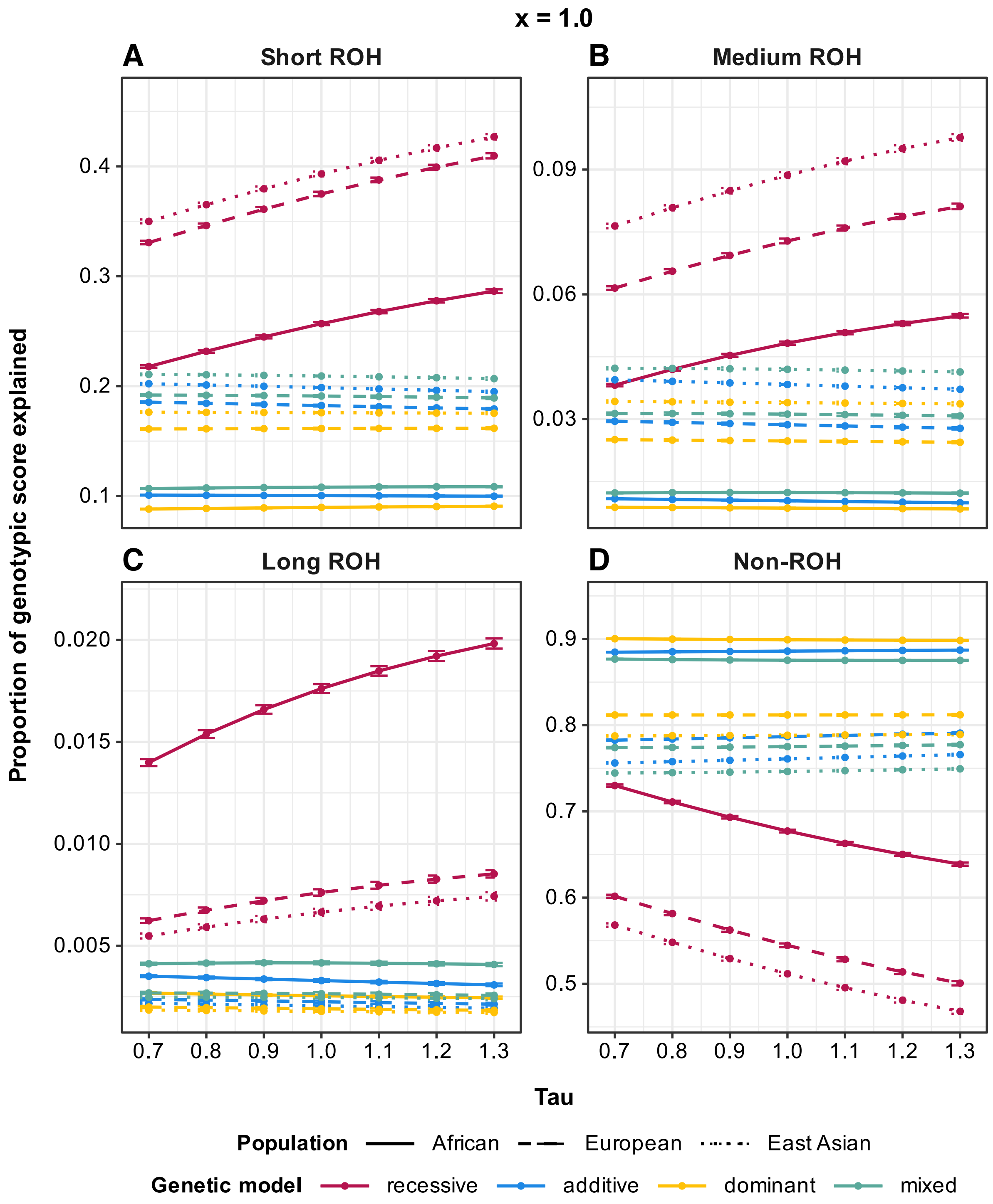


Figure S21. (Non-sparse model) Proportion of total genotypic score per individual explained by different types of regions with varied tau, various population histories and genetic architectures (x = 1.0, the fraction of simulated deleterious mutations treated as causal). Causal genotypic scores are only contributed by deleterious mutations. Panels show the proportion of genotypic score explained by (A) Short ROH regions (< 0.25 cM); (B) Medium ROH regions (0.25-1 cM); (C) Long ROH regions (> 1 cM); (D) Non-ROH.

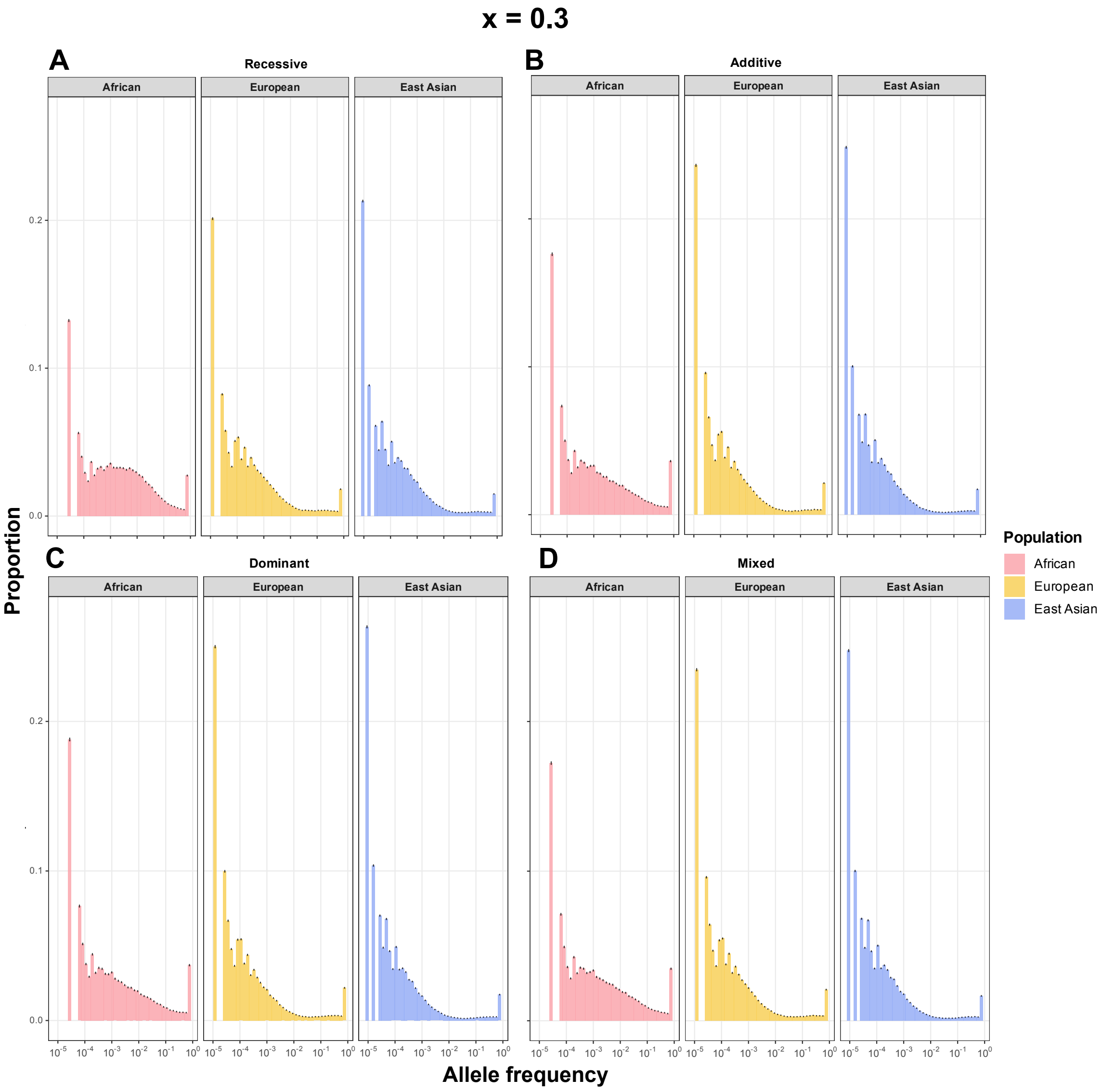


Figure S22. (Sparse model) Allele frequency spectrum of causal variants across populations and dominance models (x = 0.3, the fraction of simulated deleterious mutations treated as causal). Spectra are shown for African (AFR), European (EUR), and East Asian (EAS) populations under (A) fully recessive, (B) fully additive, (C) fully dominant, and (D) mixed models. x-axis on log scale.

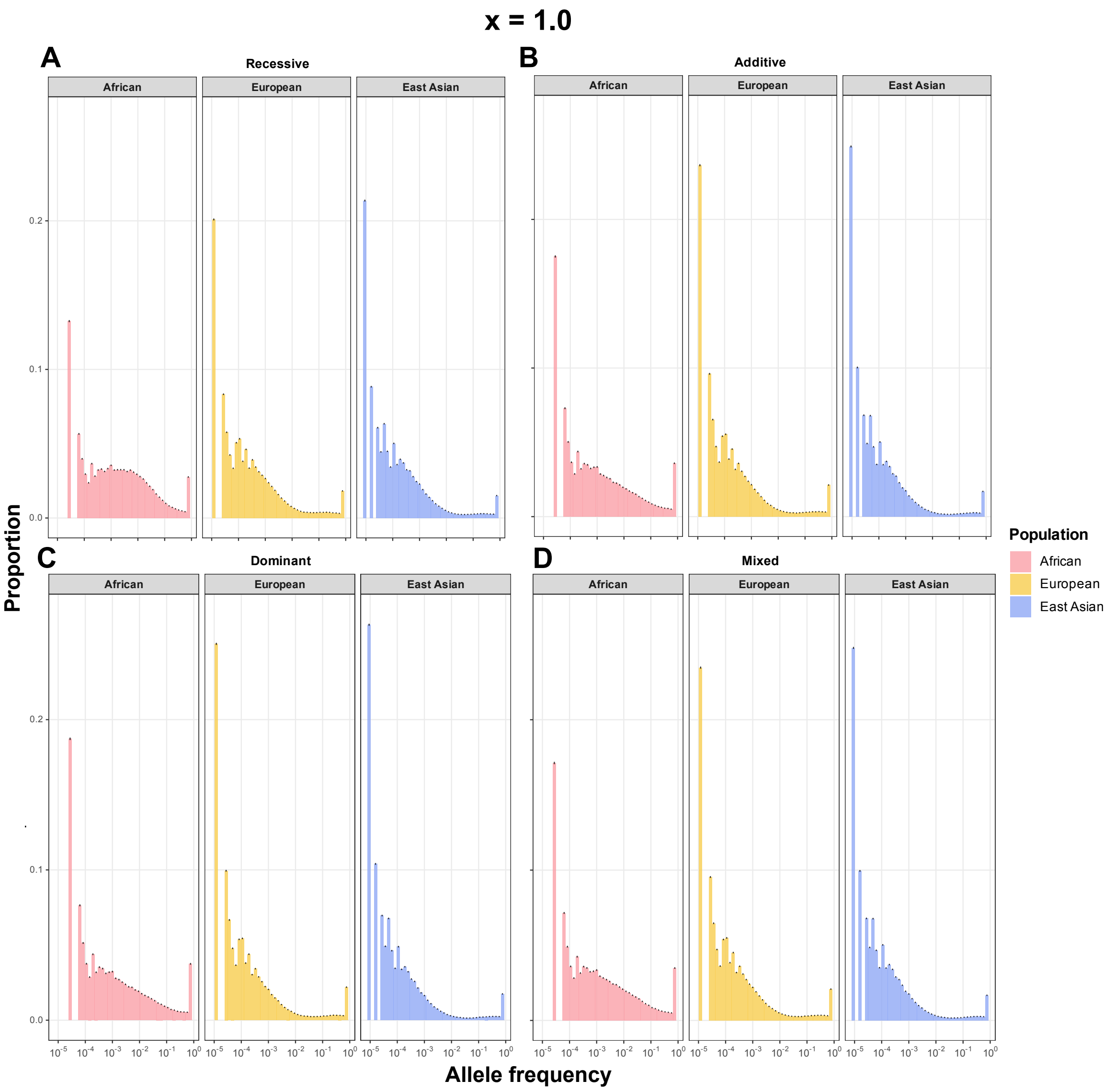


Figure S23. (Non-sparse model) Allele frequency spectrum of causal variants across populations and dominance models (x = 1.0, the fraction of simulated deleterious mutations treated as causal). Spectra are shown for African (AFR), European (EUR), and East Asian (EAS) populations under (A) fully recessive, (B) fully additive, (C) fully dominant, and (D) mixed models. x-axis on log scale.

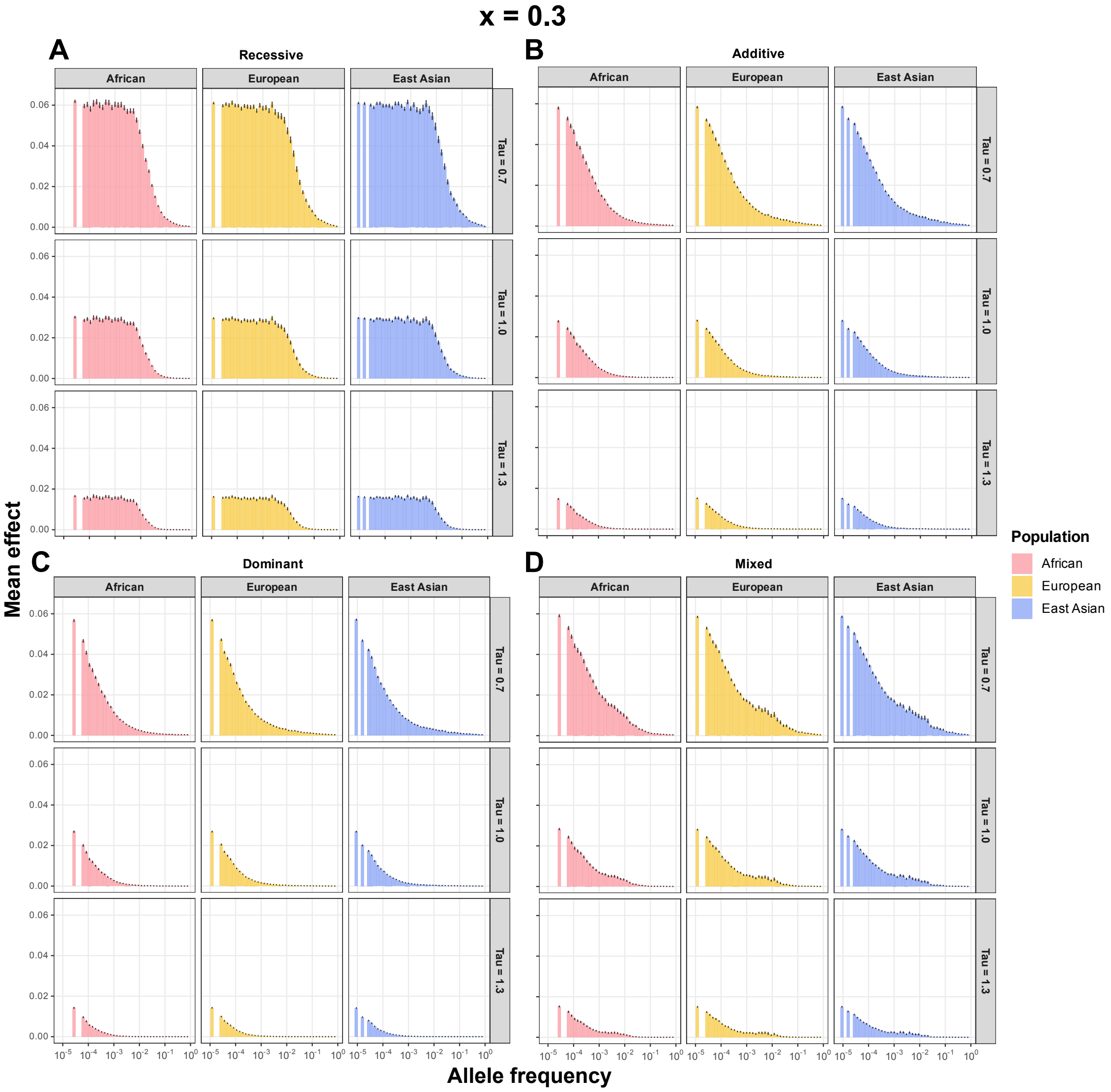


Figure S24. (Sparse model) Effect frequency spectrum of causal variants across populations and dominance models (x = 0.3, the fraction of simulated deleterious mutations treated as causal). Spectra are shown for African (AFR), European (EUR), and East Asian (EAS) populations under (A) fully recessive, (B) fully additive, (C) fully dominant, and (D) mixed models. x-axis on log scale.

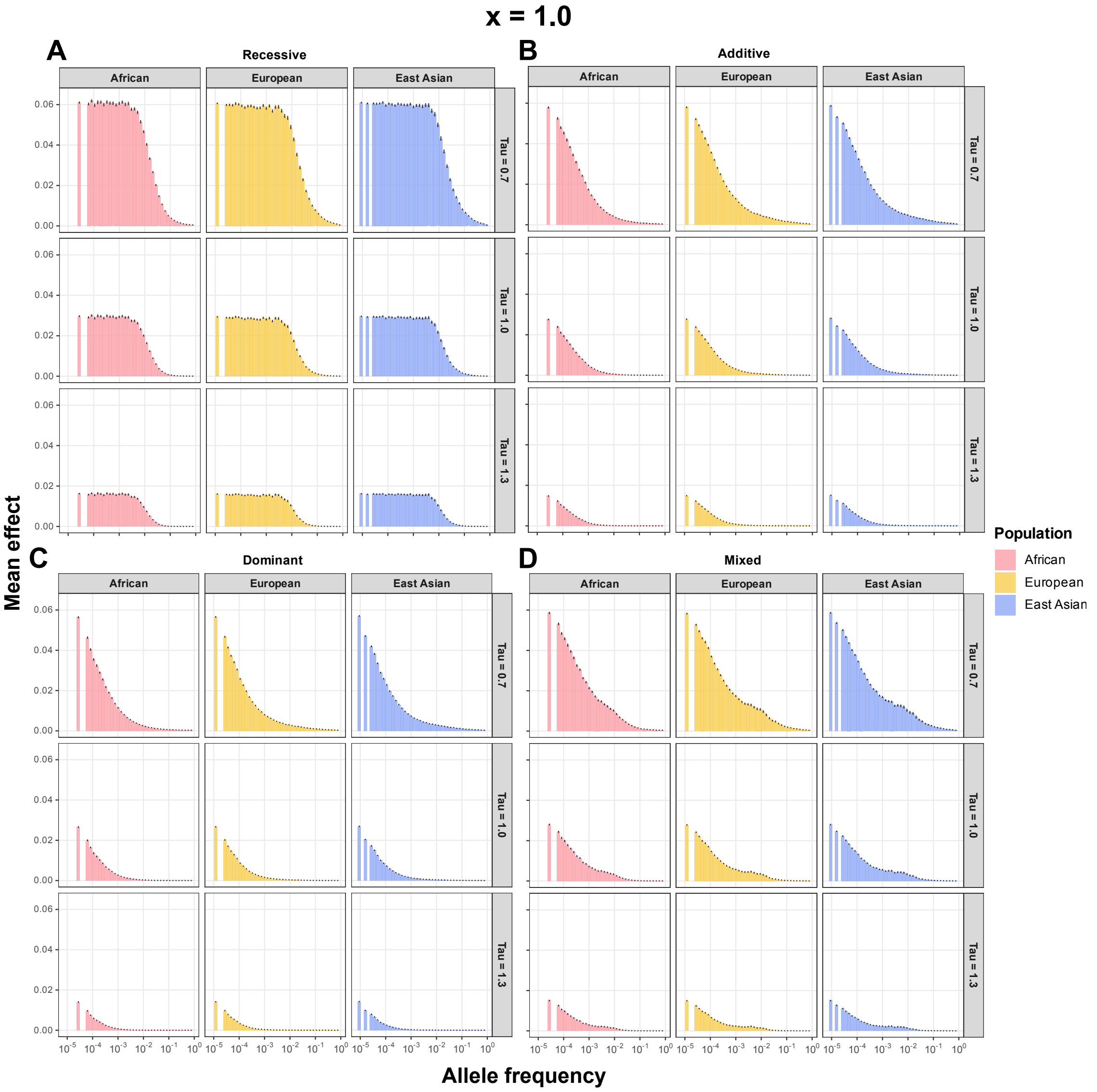


Figure S25. (Non-sparse model) Effect frequency spectrum of causal variants across populations and dominance models (x = 1.0, the fraction of simulated deleterious mutations treated as causal). Spectra are shown for African (AFR), European (EUR), and East Asian (EAS) populations under (A) fully recessive, (B) fully additive, (C) fully dominant, and (D) mixed models. x-axis on log scale.

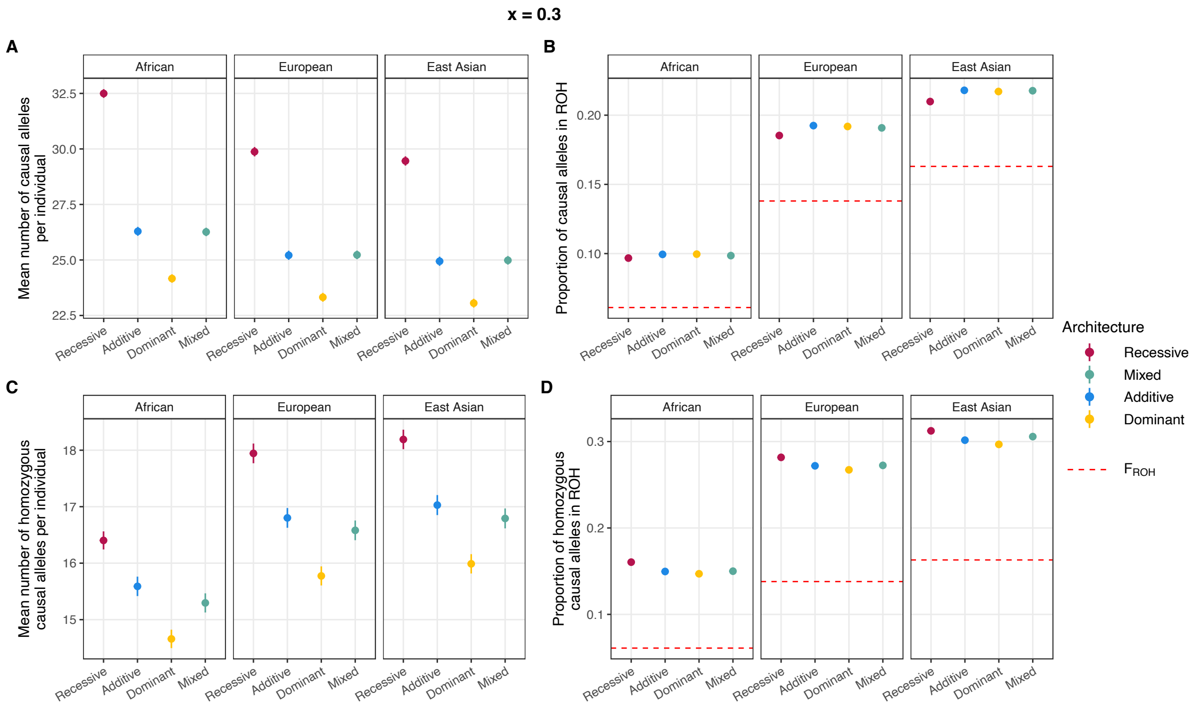


Figure S26. (Sparse model) Diagnostics of causal alleles across populations and dominance models (x = 0.3, the fraction of simulated deleterious mutations treated as causal). (A) Mean number of causal alleles per individual. (B) Proportion of causal alleles located in ROH regions. (C) Mean number of homozygous causal alleles per individual. (D) Proportion of homozygous causal alleles located in ROH regions. All quantities are shown for African (AFR), European (EUR), and East Asian (EAS) populations under fully recessive, fully additive, fully dominant, and mixed models. Red dashed lines in (B) and (D) indicate FROH, the expected proportion under random distribution of causal alleles across the genome.

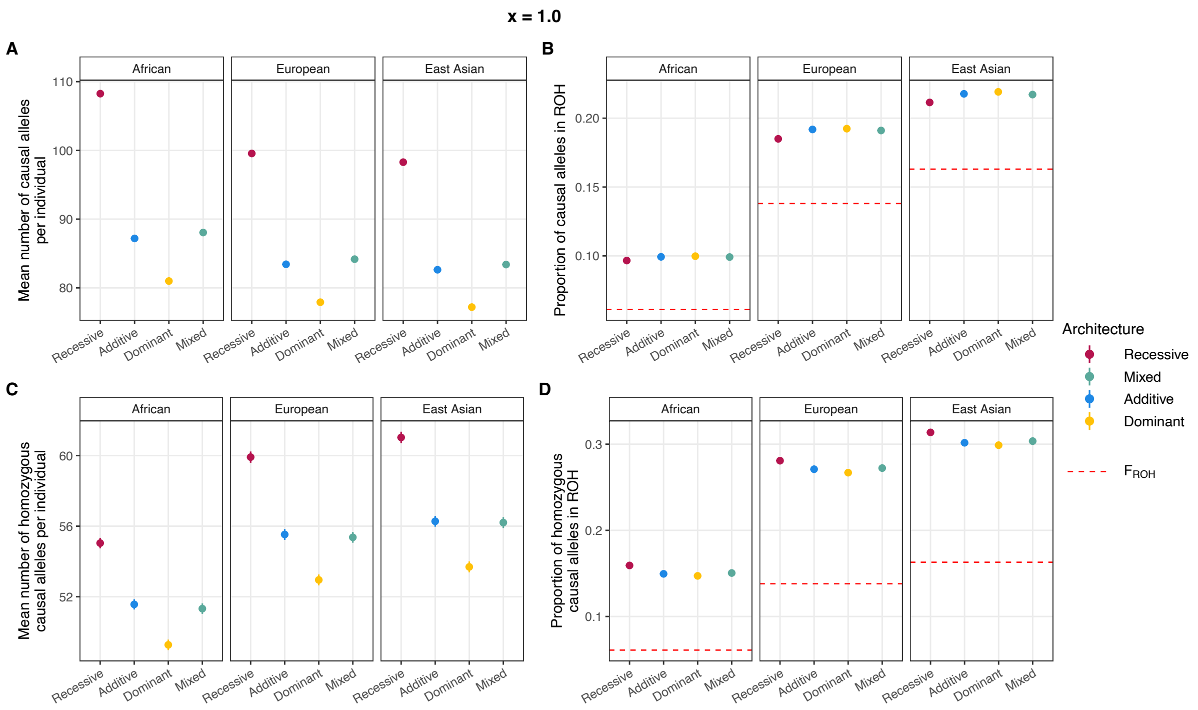


Figure S27. (Non-sparse model) Diagnostics of causal alleles across populations and dominance models (x = 1.0, the fraction of simulated deleterious mutations treated as causal). (A) Mean number of causal alleles per individual. (B) Proportion of causal alleles located in ROH regions. (C) Mean number of homozygous causal alleles per individual. (D) Proportion of homozygous causal alleles located in ROH regions. All quantities are shown for African (AFR), European (EUR), and East Asian (EAS) populations under fully recessive, fully additive, fully dominant, and mixed models. Red dashed lines in (B) and (D) indicate FROH, the expected proportion under random distribution of causal alleles across the genome.

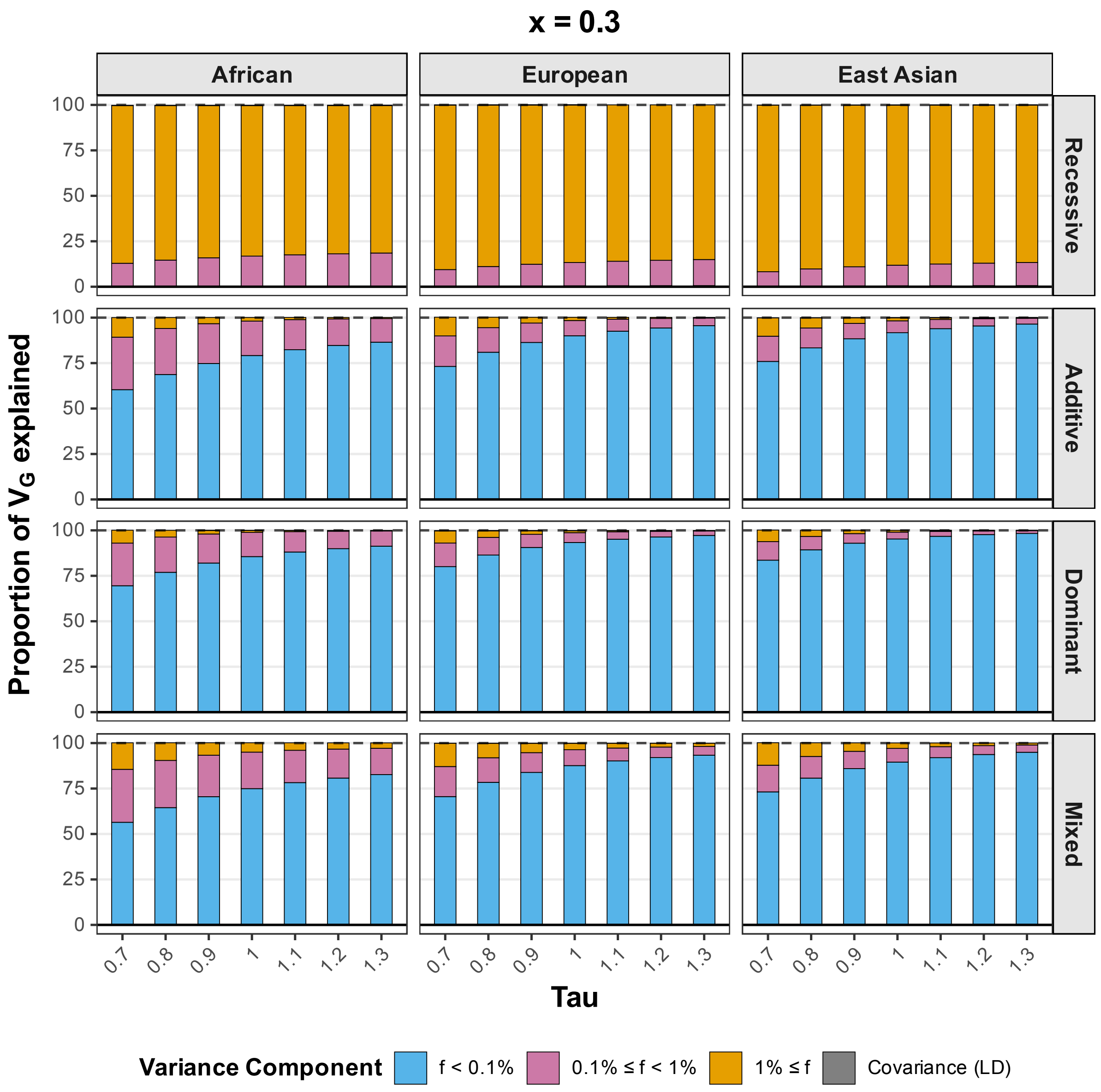


Figure S28. (Sparse model) Proportion of genetic variance explained by rare versus common alleles across populations and dominance models (x = 0.3, the fraction of simulated deleterious mutations treated as causal). Stacked bars show the proportion of total genetic variance ($V_G$) attributable to causal variants in three allele frequency bins (f < 0.1%; 0.1% ≤ f < 1%; 1% ≤ f) and to LD-induced covariance, as a function of τ (0.7–1.3), for African (AFR), European (EUR), and East Asian (EAS) populations under fully recessive, fully additive, fully dominant, and mixed models.

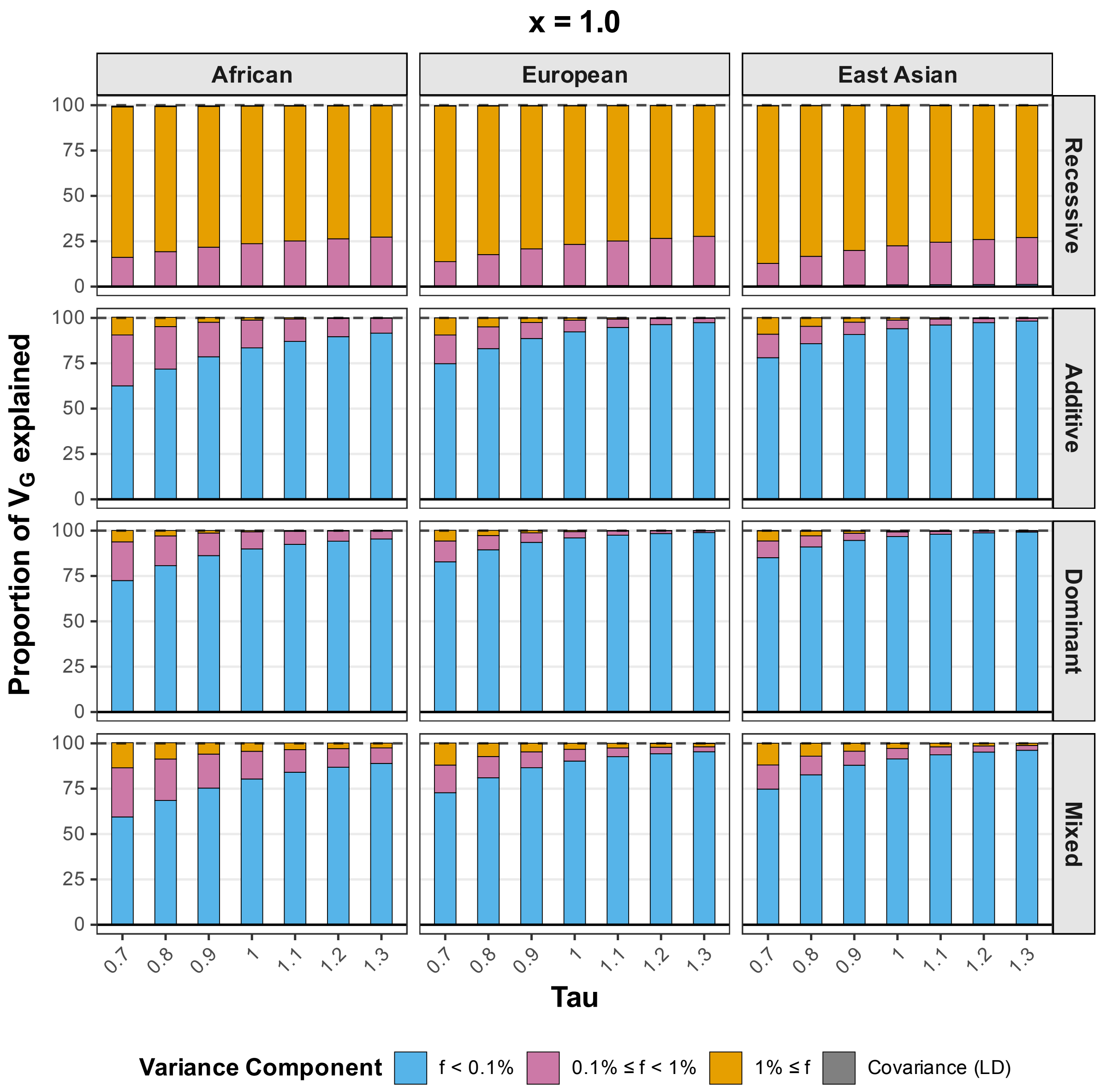


Figure S29. (Non-sparse model) Proportion of genetic variance explained by rare versus common alleles across populations and dominance models (x = 1.0, the fraction of simulated deleterious mutations treated as causal). Stacked bars show the proportion of total genetic variance ($V_G$) attributable to causal variants in three allele frequency bins (f < 0.1%; 0.1% ≤ f < 1%; 1% ≤ f) and to LD-induced covariance, as a function of τ (0.7–1.3), for African (AFR), European (EUR), and East Asian (EAS) populations under fully recessive, fully additive, fully dominant, and mixed models.

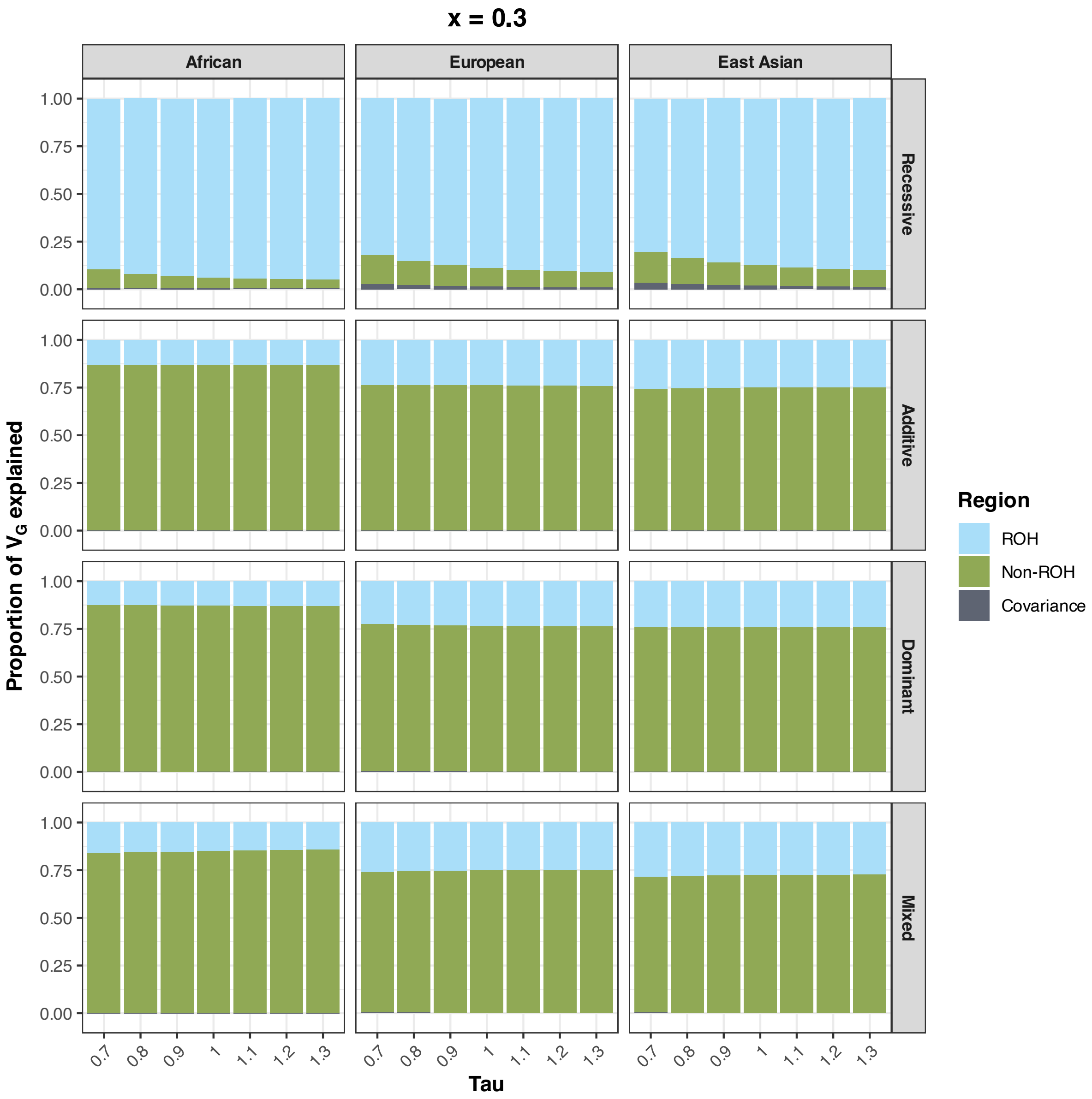


Figure S30. (Sparse model) Proportion of genetic variance ($V_G$) explained by ROH versus non-ROH regions across populations and dominance models (x = 0.3, the fraction of simulated deleterious mutations treated as causal). Stacked bars show the proportion of total genetic variance attributable to causal variants located in ROH regions, in non-ROH regions, and to the covariance between the two, as a function of τ (0.7–1.3), for African (AFR), European (EUR), and East Asian (EAS) populations under fully recessive, fully additive, fully dominant, and mixed models.

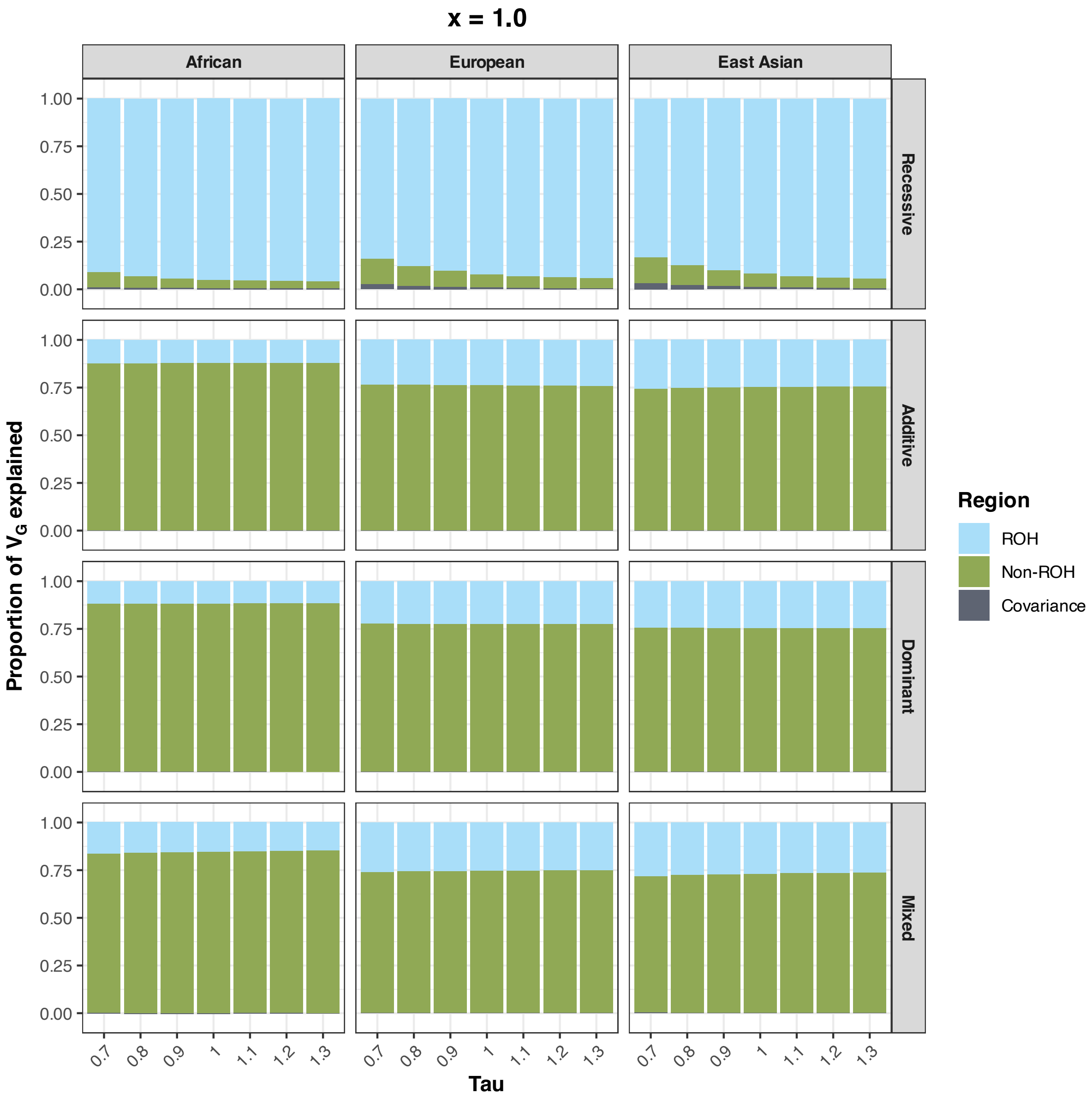


Figure S31. (Non-sparse model) Proportion of genetic variance ($V_G$) explained by ROH versus non-ROH regions across populations and dominance models (x = 1.0, the fraction of simulated deleterious mutations treated as causal). Stacked bars show the proportion of total genetic variance attributable to causal variants located in ROH regions, in non-ROH regions, and to the covariance between the two, as a function of τ (0.7–1.3), for African (AFR), European (EUR), and East Asian (EAS) populations under fully recessive, fully additive, fully dominant, and mixed models.

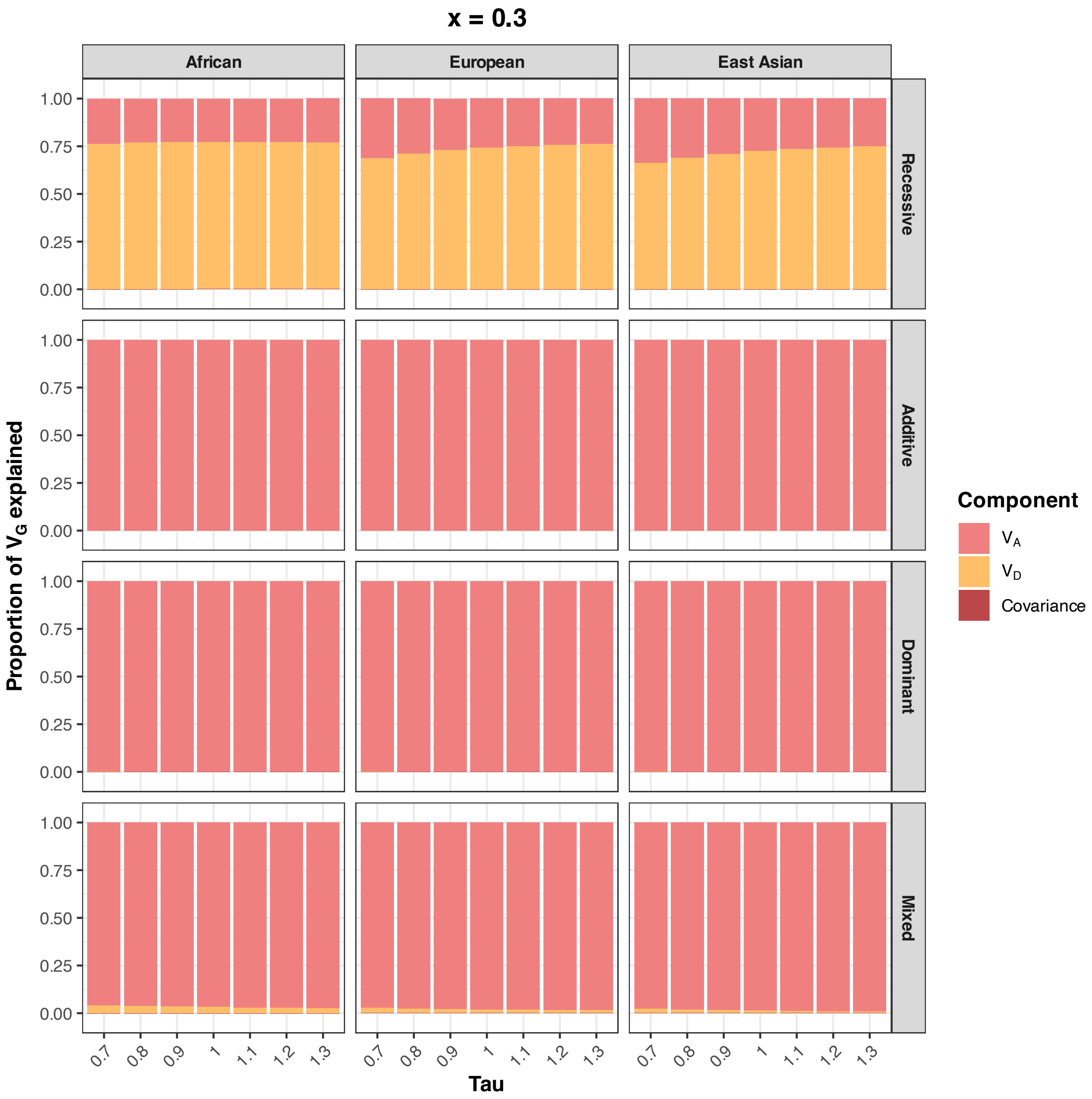


Figure S32. (Sparse model) Proportion of genetic variance ($V_G$) explained by additive ($V_A$), dominance ($V_D$), and covariance components across populations and dominance models (x = 0.3, the fraction of simulated deleterious mutations treated as causal). Stacked bars show the proportion of total genetic variance ($V_G$) attributable to $V_A$, $V_D$, and the $V_A$-$V_D$ covariance, as a function of τ (0.7–1.3), for African (AFR), European (EUR), and East Asian (EAS) populations under fully recessive, fully additive, fully dominant, and mixed models.

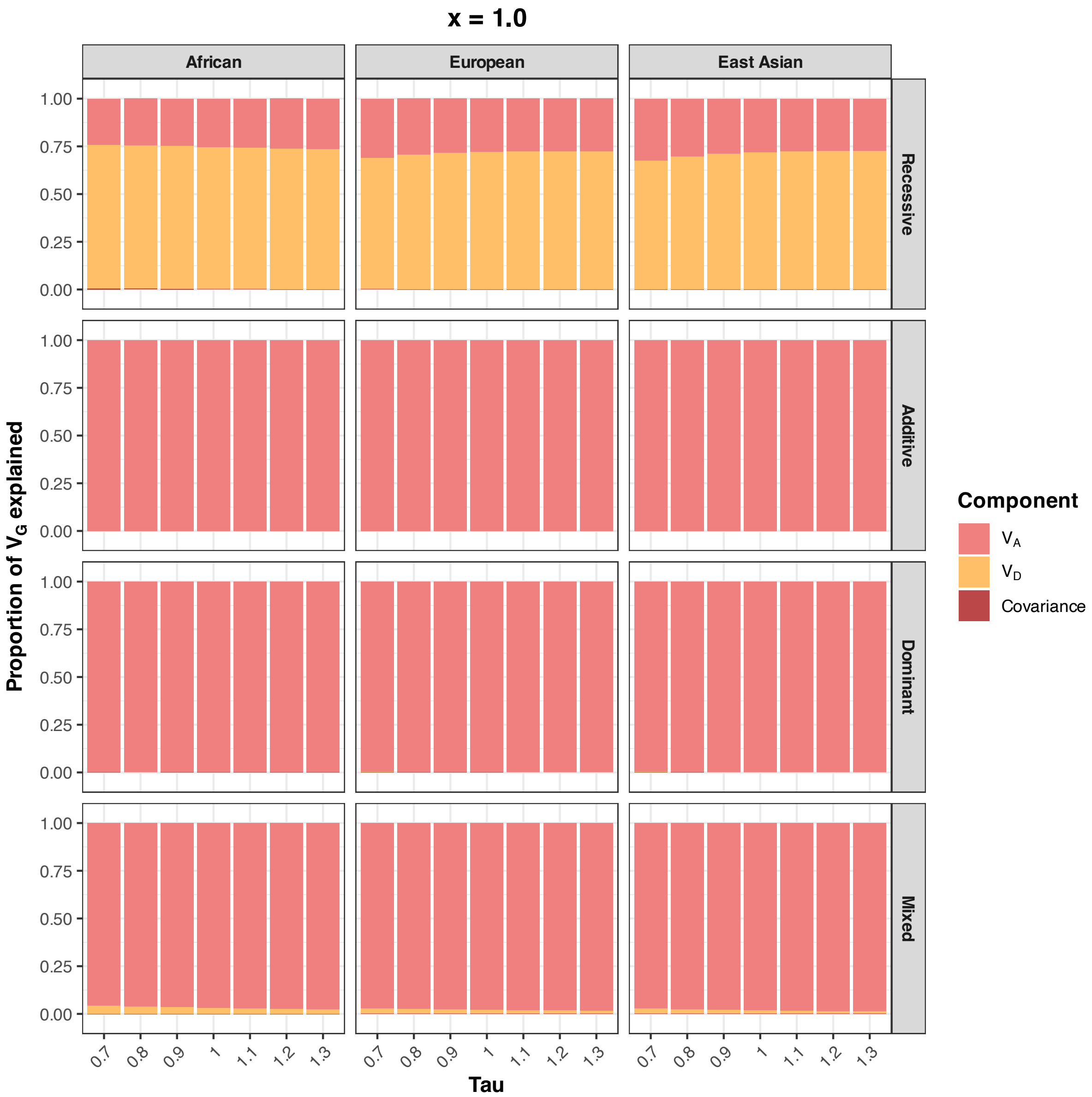


Figure S33. (Non-sparse model) Proportion of genetic variance ($V_G$) explained by additive ($V_A$), dominance ($V_D$), and covariance components across populations and dominance models (x = 1.0, the fraction of simulated deleterious mutations treated as causal). Stacked bars show the proportion of total genetic variance ($V_G$) attributable to $V_A$, $V_D$, and the $V_A$-$V_D$ covariance, as a function of τ (0.7–1.3), for African (AFR), European (EUR), and East Asian (EAS) populations under fully recessive, fully additive, fully dominant, and mixed models.

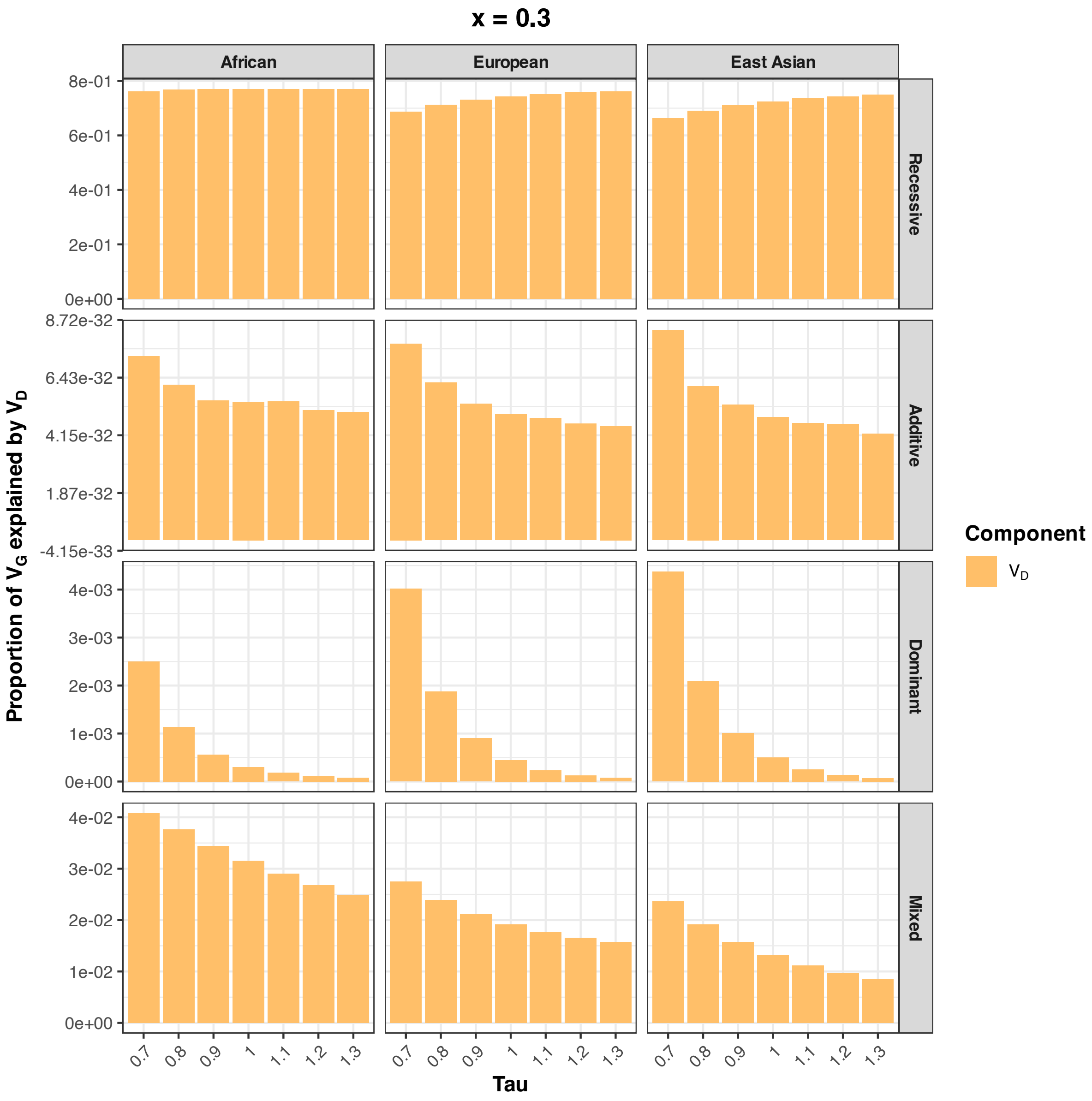


Figure S34. (Sparse model) Proportion of genetic variance explained by dominance variance ($V_D$) across populations and dominance models (x = 0.3, the fraction of simulated deleterious mutations treated as causal). Bars show the proportion of total genetic variance ($V_G$) attributable to $V_D$ alone, as a function of τ (0.7–1.3), for African (AFR), European (EUR), and East Asian (EAS) populations under fully recessive, fully additive, fully dominant, and mixed models. y-axis scales differ across dominance models to make the small $V_D$ proportions under fully additive, fully dominant, and mixed models visible; values under the fully additive model are at machine-precision zero (~$10^{-32}$).

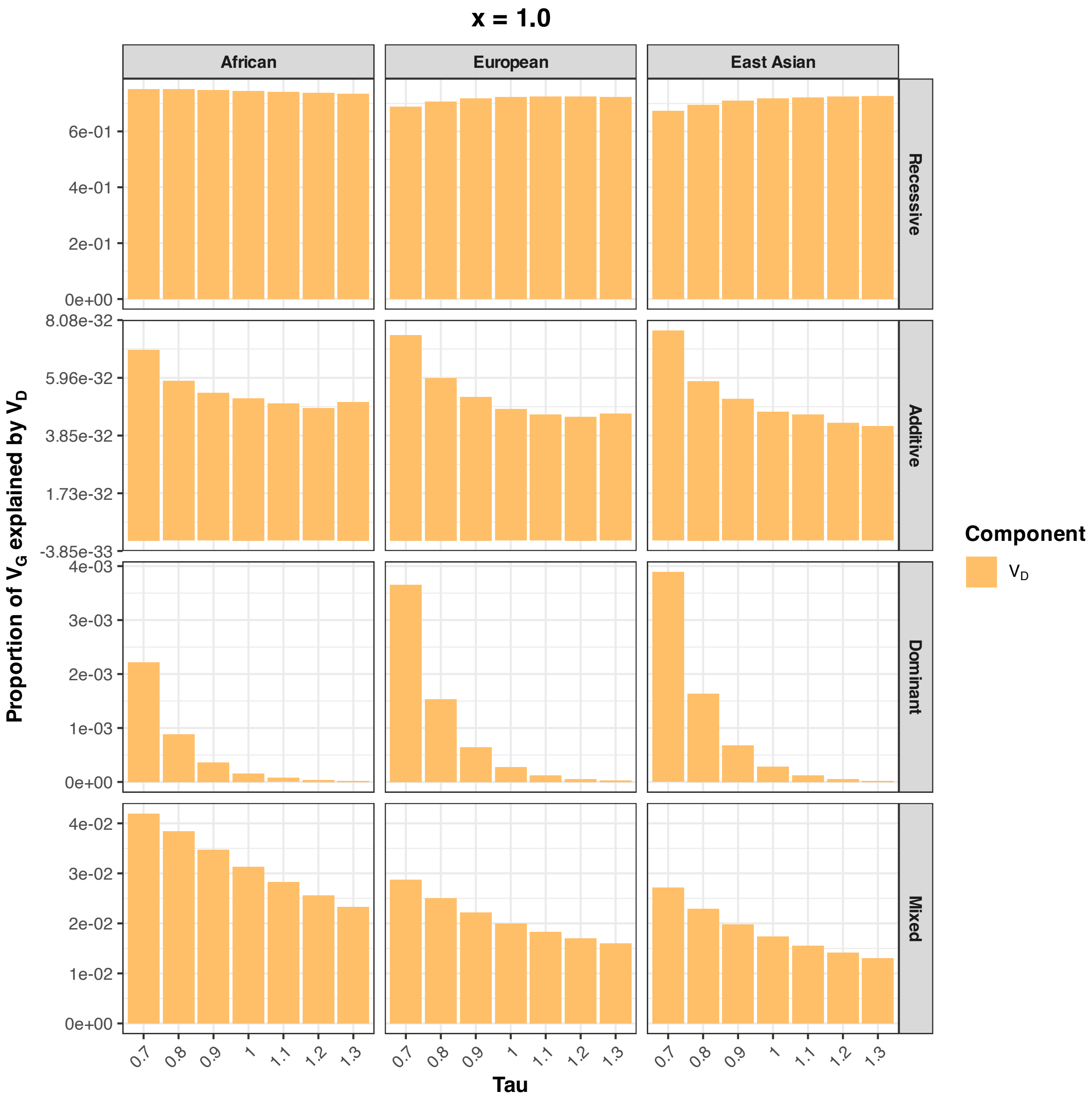


Figure S35. (Non-sparse model) Proportion of genetic variance explained by dominance variance ($V_D$) across populations and dominance models (x = 1.0, the fraction of simulated deleterious mutations treated as causal). Bars show the proportion of total genetic variance ($V_G$) attributable to $V_D$ alone, as a function of τ (0.7–1.3), for African (AFR), European (EUR), and East Asian (EAS) populations under fully recessive, fully additive, fully dominant, and mixed models. y-axis scales differ across dominance models to make the small $V_D$ proportions under fully additive, fully dominant, and mixed models visible; values under the fully additive model are at machine-precision zero (~$10^{-32}$).

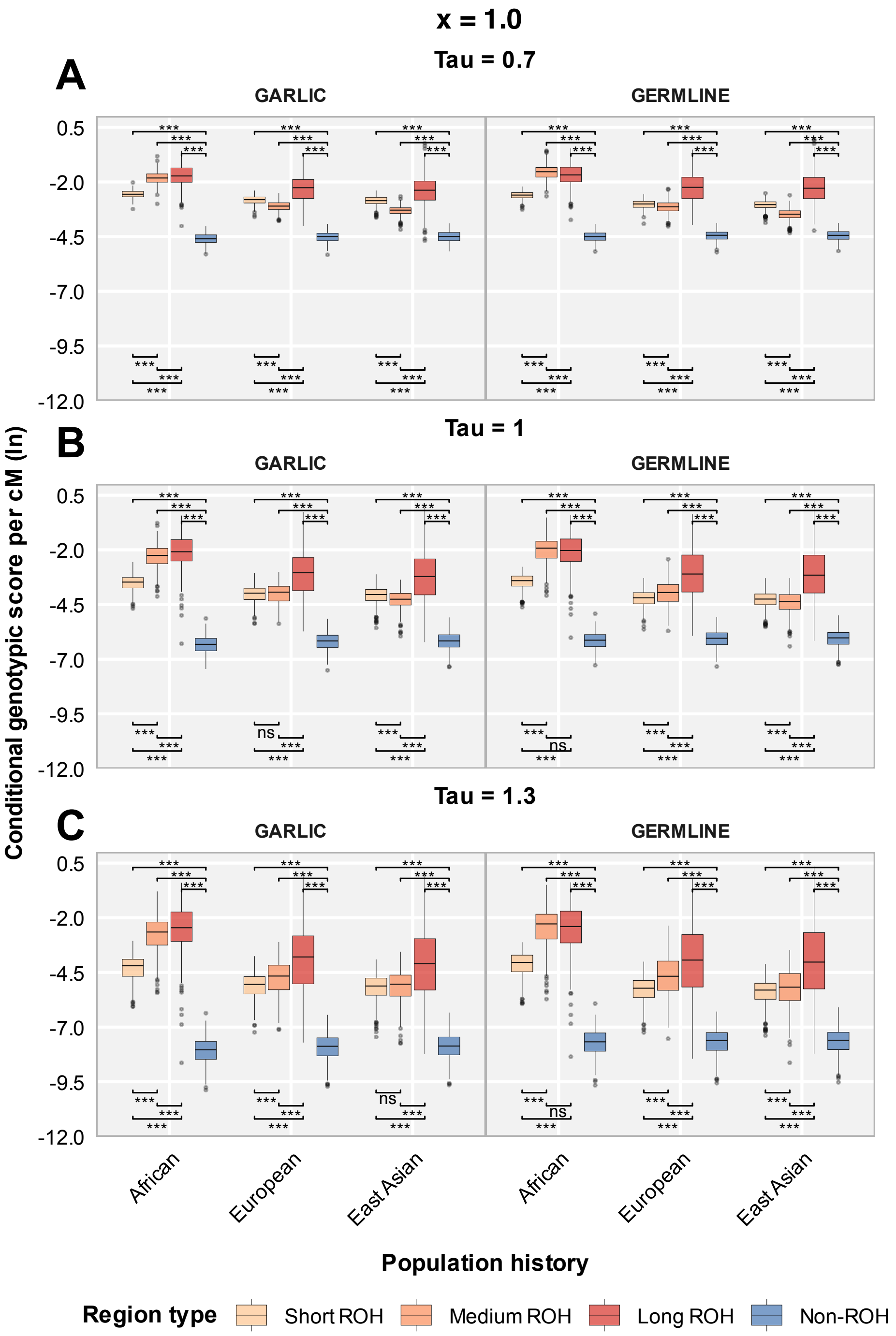


Figure S36. (Non-sparse model) Conditional per-cM genotypic score contribution (log-transformed; deleterious mutations) under the fully recessive model, comparing GARLIC and GERMLINE ROH callers across population histories (x = 1.0, the fraction of simulated deleterious mutations treated as causal). Regions are: Short ROH (< 0.25 cM); Medium ROH (0.25–1 cM); Long ROH (> 1 cM); Non-ROH. Causal genotypes are generated from deleterious mutations. The parameter tau is varied for different weighting of rare alleles in phenotype score calculation. (A) $\tau = 0.7$, (B) $\tau = 1.0$, (C) $\tau = 1.3$. Each panel compares ROH called by GARLIC (left) and GERMLINE (right). Significance levels for adjusted P-values: *** P_adj < 0.001, ** P_adj < 0.01, * P_adj < 0.05, ns P_adj ≥ 0.05.

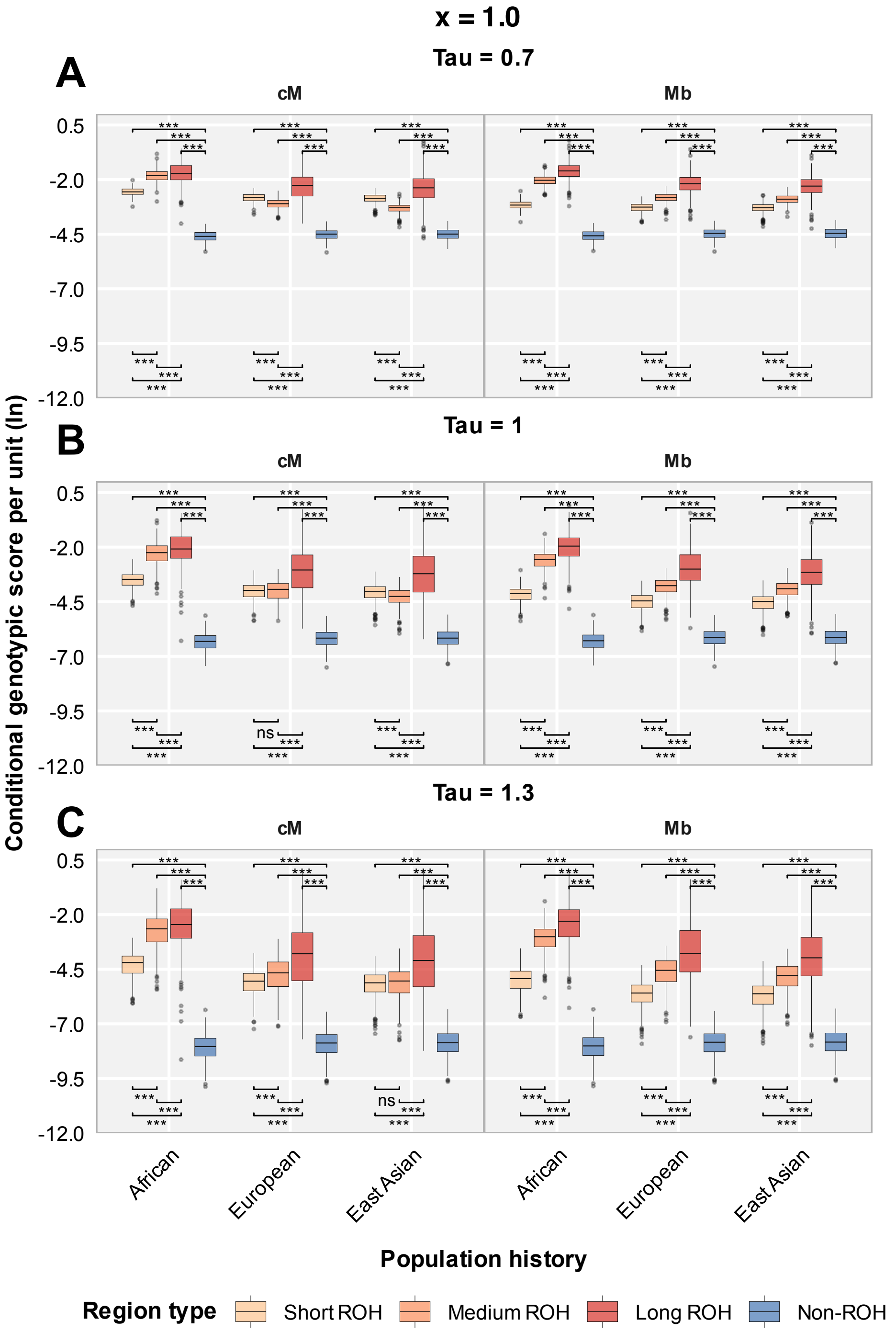


Figure S37. (Non-sparse model) Conditional per-unit genotypic score contribution (log-transformed; deleterious mutations) under the fully recessive model, comparing cM-based and Mb-based ROH classifications across population histories (x = 1.0, the fraction of simulated deleterious mutations treated as causal). Regions are: Short ROH (< 0.25 cM or < 0.25 Mb); Medium ROH (0.25–1 cM or 0.25–1 Mb); Long ROH (> 1 cM or > 1 Mb); Non-ROH. Causal genotypes are generated from deleterious mutations. The parameter tau is varied for different weighting of rare alleles in phenotype score calculation. (A) $\tau = 0.7$, (B) $\tau = 1.0$, (C) $\tau = 1.3$. Each panel compares cM-based classification (left) and Mb-based classification (right). Significance levels for adjusted P-values: *** $P_{adj} < 0.001$, ** $P_{adj} < 0.01$, * $P_{adj} < 0.05$, ns $P_{adj} \geq 0.05$.

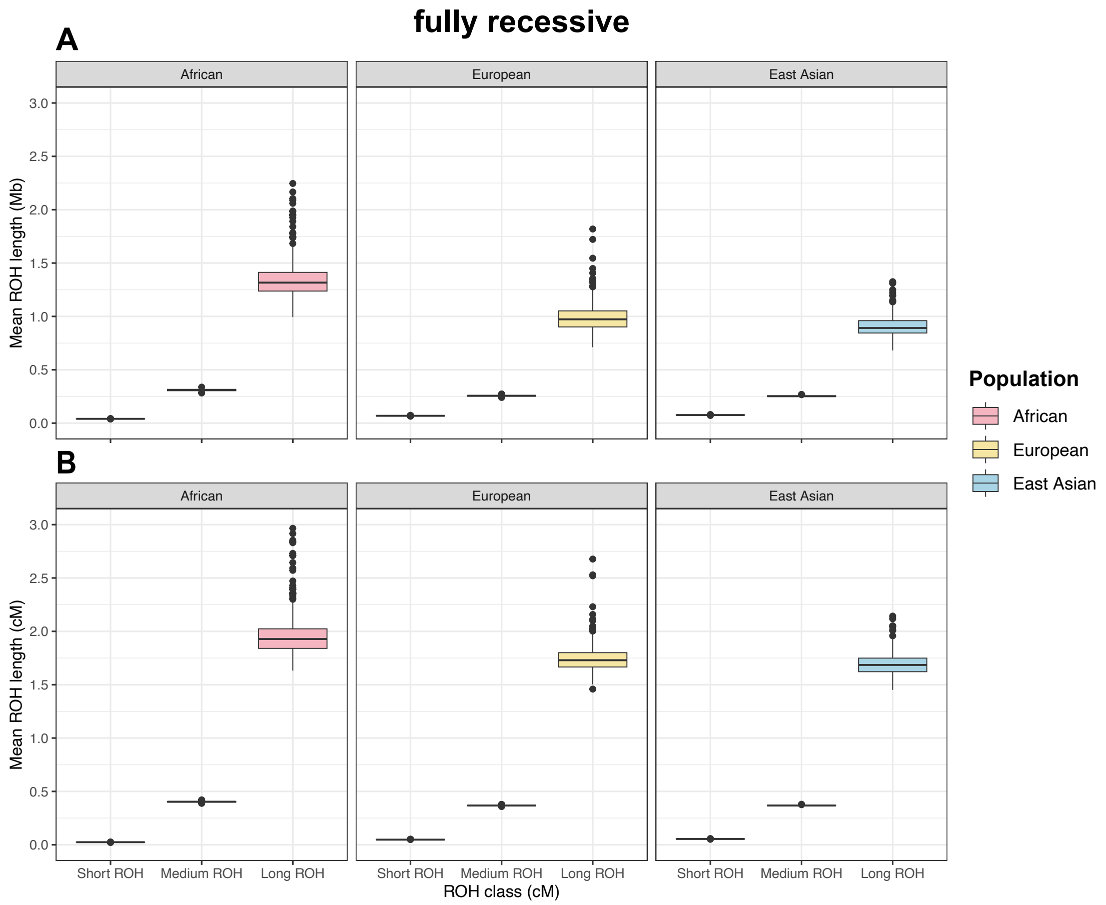


Figure S38. Length distribution of ROH classes under the fully recessive model, in Mb and cM. Box plots show the mean ROH length per individual in (A) Mb and (B) cM, for short (< 0.25 cM), medium (0.25–1 cM), and long (> 1 cM) ROH classes (cM-based classification), across African (AFR), European (EUR), and East Asian (EAS) populations.
